# SnoBIRD: A tool to identify C/D box snoRNAs and refine their annotation across all eukaryotes

**DOI:** 10.1101/2025.04.01.646650

**Authors:** Étienne Fafard-Couture, Cédric Boulanger, Laurence Faucher-Giguère, Vanessa Sinagoga, Mélodie Berthoumieux, Jordan Hedjam, Virginie Marcel, Sébastien Durand, Mark A. Bayfield, François Bachand, Sherif Abou Elela, Pierre-Étienne Jacques, Michelle S. Scott

## Abstract

Small nucleolar RNAs (snoRNAs), a group of noncoding RNAs present amongst all eukaryotes, are most extensively characterized for their regulation of ribosome biogenesis and splicing. Despite their central roles, current snoRNA annotations remain incomplete. Several eukaryote genome annotations contain few or no snoRNAs, and none distinguish expressed snoRNAs from their pseudogenes—a recently characterized snoRNA subclass with distinct features and expression levels. To address this, we developed SnoBIRD, a BERT-based C/D box snoRNA predictor trained on snoRNAs spanning all eukaryote kingdoms. We show that SnoBIRD outperforms existing tools and is the only predictor capable of identifying snoRNA pseudogenes using biologically relevant signal. Applied on the fission yeast and human genomes, we demonstrate that only SnoBIRD scales well with genome size in terms of runtime, and we identify and experimentally validate several new SnoBIRD-predicted C/D box snoRNAs. By running SnoBIRD on multiple eukaryote genomes, we identify hundreds of novel snoRNA candidates and highlight SnoBIRD’s usefulness to determine the evolutionary paths of snoRNAs distributed across different species. Overall, SnoBIRD represents a user-friendly and efficient tool for reliably predicting C/D box snoRNAs and their pseudogenes across any eukaryote genome.

**Graphical abstract:** 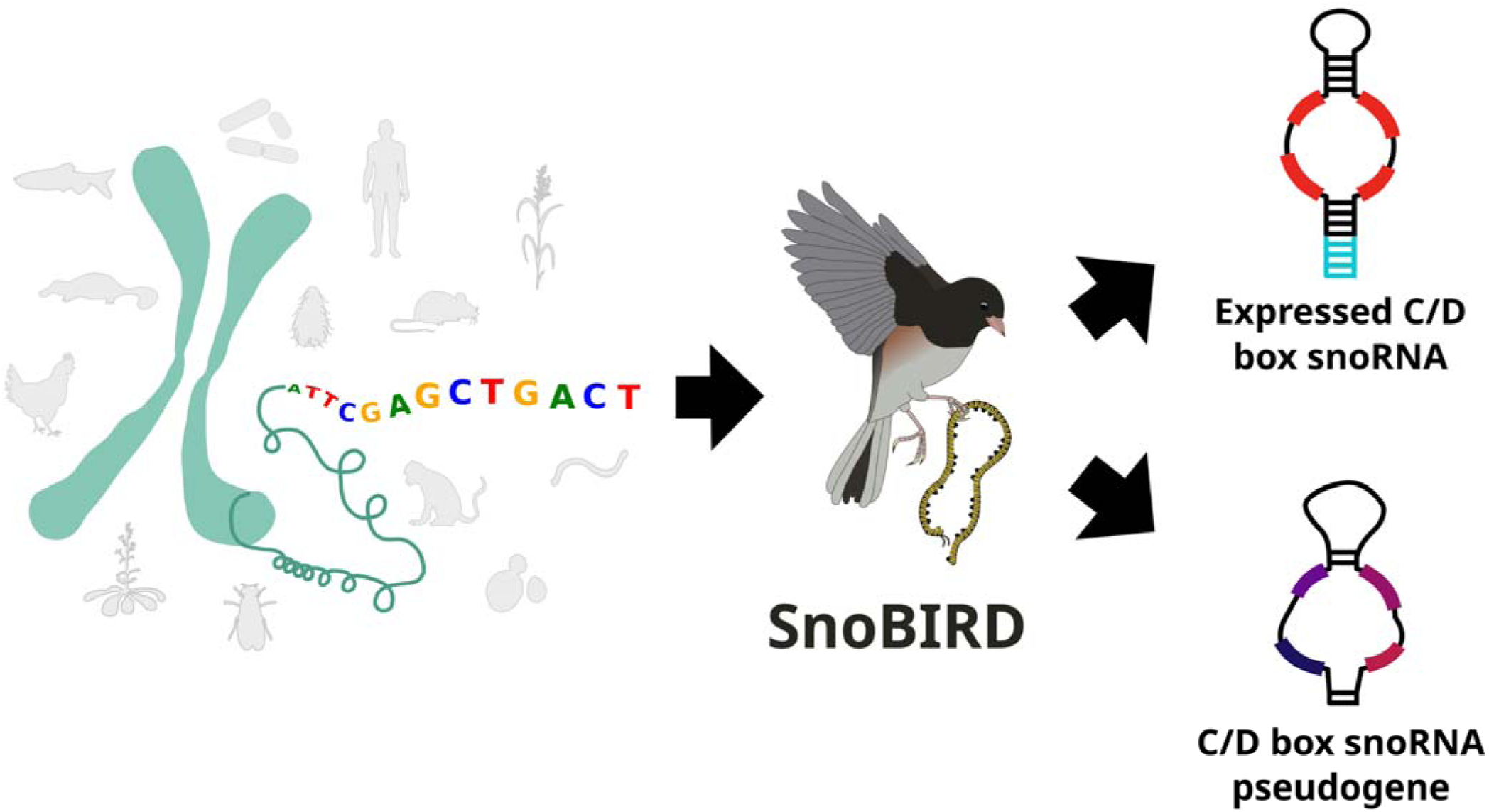

## INTRODUCTION

Small nucleolar RNAs (snoRNAs) represent an ancient group of highly structured noncoding RNAs present in all eukaryotes. SnoRNAs are generally categorized in two types which differ in their sequence motifs, protein interactors and functions: the C/D and H/ACA box snoRNAs, the name of which has been respectively standardized to start with SNORD and SNORA in human. Termed after the canonical motifs they harbor (namely the C/C’ and D/D’ boxes (respectively RUGAUGA and CUGA), and the H and ACA boxes (respectively ANANNA and ACA), where R is a purine and N is any nucleotide), snoRNAs exist in their mature form as ribonucleoprotein complexes (snoRNPs), i.e. bound to their core protein partners which confer the stability and catalytic activity to the snoRNP (1, 2). As a crucial member of the complex, snoRNAs act as guides by base-pairing to a specific target RNA which allows dedicated enzymes to modify specific nucleotides in the recipient RNA (3). The most well-characterized targets of snoRNAs are ribosomal RNAs (rRNA) and small nuclear RNAs (snRNAs) which undergo several important chemical modifications as part of ribosome and spliceosome biogenesis steps, namely C/D-guided 2’-O-methylations and H/ACA-guided pseudouridylations (4, 5). While guiding these chemical modifications was considered for decades to be the main purpose of snoRNAs, a multitude of new functions and noncanonical targets have been associated with snoRNAs in recent years, several of which do not involve RNA modification (reviewed in (6–8)). These include the regulation of splicing (9–13), chromatin remodeling (14, 15), protein secretion (16) as well as transfer RNA (tRNA) and transcript stability (17, 18) through the interaction of snoRNAs with different effectors (messenger RNAs (mRNAs), RNA binding proteins (RBPs), etc.). Overall, these diverse cellular involvements underscore that snoRNAs are increasingly recognized as versatile key players in the regulation of gene expression.

While most snoRNAs in human are embedded within the introns of so-called host genes, the genomic location of snoRNAs varies substantially across species and eukaryote kingdoms (8, 19, 20). For instance, most plants and fungi snoRNAs are expressed as independent units using their own promoter, transcribed either alone or as a polycistronic transcript with other snoRNAs located in a cluster (8, 20). On the other hand, some snoRNAs in vertebrates are also transcribed as intergenic units, but most rely on the transcription and splicing of their host gene locus to be expressed (21). The association between intronic snoRNAs and their host gene appears to be under strong evolutionary pressure, as snoRNA host loci are strongly enriched for genes involved in ribosome biogenesis and RNA processing (19, 22–24). While these genomic locations are often conserved across species, several “new” snoRNA genomic locations arose throughout evolution and across species, thanks to countless rounds of gene duplications (via *cis*-recombination and retrotransposition events) which simultaneously led to the expansion of snoRNA families and copy numbers (25–28).

Interestingly, not all these snoRNAs or their copies are equally expressed. In fact, we and others have recently shown that a significant proportion of annotated snoRNA genes are not expressed (sometimes as many as two-thirds of all annotated snoRNAs) (19, 24, 29, 30). Investigating the reasons explaining this widespread non-expression, it was demonstrated that four features could reliably distinguish the expressed from the non-expressed snoRNAs (which are also called snoRNA pseudogenes) (24). Indeed, snoRNA pseudogenes display increased mutations in their characteristic boxes, an unstable secondary structure and an unstable terminal stem (i.e. a structure formed by the base-pairing of both snoRNA ends) compared to the expressed snoRNAs, as well as a non-expressed host locus (24). These snoRNA pseudogenes were hypothesized to have either transient functions in a rapidly degraded host transcript and/or represent snoRNA remnants of past duplication events in loci unfavorable for their expression (8, 24). Taken together, these observations represent an additional layer of snoRNA complexity (which is yet to be considered in current annotations) since these two subclasses show distinct sequence features and functionalities.

Moreover, the latest gene annotations do not reflect the full range and complexity of snoRNA repertoires for several reasons. Firstly, a recent effort highlighted that snoRNAs are not as evenly and adequately represented in the different eukaryote kingdom annotations (8). Indeed, it was shown that fungus and protist species displayed fewer than expected or sometimes even no snoRNA genes in their current Ensembl annotation (8, 31). Secondly, several species including model species (e.g. *Drosophila melanogaster*, *Caenorhabditis elegans*, *Arabidopsis thaliana*, etc.) display incomplete snoRNA annotation, many lacking information about the snoRNA type (8). Finally, and as mentioned earlier, the quite recent characterization of snoRNA pseudogenes brings the challenge that they are not yet considered in the latest gene annotations. Therefore, these observations clearly underline that current gene annotations lack the comprehensiveness and granularity to fully investigate the multiple facets of snoRNA biology.

One way to help alleviate these shortcomings is to create accurate snoRNA-specific annotation tools. Snoscan (32), the first C/D box snoRNA predictor to have been released and still one of the most widely used in the literature, is based on the Hidden Markov Model (HMM) architecture and was trained on a few budding yeast snoRNAs. The beginning of the twenty-first century next saw the introduction of several new snoRNA prediction tools including CDseeker (33), Snoreport (34) and SnoStrip (35), none of which are currently supported or installable. The latest snoRNA predictor to be released, Snoreport2 (36), was elaborated almost a decade ago as an optimized version of the original Snoreport, employing a Support Vector Machine (SVM) architecture that was trained on a limited set of different species snoRNAs. Other generalist approaches were also developed, including cmscan from the non-snoRNA-centric Infernal suite (37). This tool, when combined with Rfam covariance models (thereby dubbed Infernal_rfam in this study), searches in an input for sequence and structural homology that fit within a certain Rfam family covariance model, including snoRNA families. Unfortunately, each of the previous tools suffers from major issues which are further discussed in the following sections of this study (**Table 1**). In addition, having been released over a decade ago, their training set is highly limited in terms of the diversity of species considered. Indeed, during this last decade, many new snoRNAs were identified across a vast breadth of species (30, 38, 39), including through the development of novel techniques to specifically detect structured RNAs in a high-throughput manner (e.g., TGIRT-Seq which uses a thermostable reverse transcriptase (40, 41)).

**Table 1.** Comparison between existing C/D snoRNA predictors and SnoBIRD.

|  | *Snoscan* | *Infernal_rfam* | *Snoreport2* | *SnoBIRD* |
| --- | --- | --- | --- | --- |
| **Purpose** | Identify canonical C/D snoRNAs* | Identify C/D snoRNAs that fit in known Rfam families | Identify any C/D snoRNA | Identify any C/D snoRNA |
| **SnoRNA pseudogene detection** | False | False | False | True |
| **Strand prediction** | Both strands | Both strands | + strand only | Both strands |
| **Minimal input** | Input FASTA and rRNA FASTA | Input FASTA and covariance model | Input FASTA | Input FASTA or BED and input FASTA |
| **Output type** | FASTA with multi-line embedded information | Text file and/or table-shaped | FASTA | BED, FASTA, GTF or TSV (default) |
| **Output information** | C/D/D' motifs, genomic location, potential target, score (bits) | Genomic location, Rfam family, score (bits and E-value) | Sequence, C/D motifs, genomic location, score (probability) | Sequence, C/D/C'/D' motifs, genomic location, box conservation, structure and terminal stem stabilities, score (probability) |
| **Installation** | Conda | Conda | Conda with dependency issues** | Virtualenv / conda |
| **Parallelization** | False | Multithreading or Message Passing Interface | False | CPU/GPU multiprocessing |
| **Publication** | Lowe & Eddy, 1999 | Nawrocki & Eddy, 2013 | de Araujo Oliveira *et al.* 2016 | Fafard-Couture *et al.* 2025 (This study) |

\* Snoscan will identify only snoRNAs that present complementarity to the input rRNA sequence, thereby excluding snoRNAs with non-canonical targets
\*\* The ViennaRNA dependency is not installed automatically, and GNU Compiler Collection / C Library versions must be tweaked for Snoreport2 to work

Therefore, to take advantage of these novel snoRNA datasets and considerable advances in machine learning models over the past decade, as well as to tackle the aforementioned issues with current snoRNA annotations and snoRNA prediction tools, we created a new C/D box snoRNA predictor. To do so, we employed the state-of-the-art Bidirectional Encoder Representations from Transformers (BERT) model which was shown to perform well on tasks related to sequential inputs, including nucleotide sequences (42, 43). Our predictor, called SnoBIRD for <u>B</u>ERT-based Identification and <u>R</u>efinement of C/<u>D</u> box <u>sno</u>RNAs, takes as input any genomic sequence to identify C/D box snoRNAs, and refines their annotation by considering if they are pseudogenes or not. SnoBIRD was trained on a large dataset of validated snoRNAs representative of all eukaryote kingdoms. We demonstrate that SnoBIRD outperforms existing tools in a controlled test set environment and at the scale of genome-wide predictions. We also show that SnoBIRD uses relevant biological signal within the input sequence to predict the presence of C/D box snoRNA genes. Finally, we applied SnoBIRD across various species, underlining its discovery of several new snoRNAs and its usefulness in cross-species comparative studies, all of which help pave the way to precisely characterize snoRNAs in any eukaryotic species.

## MATERIAL AND METHODS

### Dataset collection

Expressed C/D box snoRNA genes were collected from different publications (*D. melanogaster* (44), *C. elegans* (45), *Gallus gallus* (27), *Ornithorhynchus anatinus* (26), *Macaca mulatta* (46), *Tetrahymena thermophila* (47), *Leishmania major* (48), *Dictyostelium discoideum* (49), *Giardia lamblia* (50), *A. thaliana* (51, 52), *Oryza sativa* (53), *Ostreococcus tauri* (38), *Aspergillus fumigatus* (54), *Neurospora crassa* (55), *Candida albicans* (56)) and were considered in the initial dataset if they met at least one of the following criteria: detected in a cDNA library, detected by Northern blot, detected by RT-PCR or detected by primer extension. In addition, expressed C/D box snoRNAs and C/D box snoRNA pseudogenes were included in the initial dataset based on their abundance level in TGIRT-Seq datasets for *Saccharomyces cerevisiae*, *D. melanogaster*, *Mus musculus* and *Homo sapiens* (i.e. species with enough datasets to reliably assess snoRNA expression globally). These datasets were either collected from the literature (untreated or retinoic acid-treated embryonic stem cells from *M. musculus* (29); S2R cells, head and ovary samples from *D. melanogaster* (30); brain, breast, ovary, liver, prostate, testis, skeletal muscle, plasma, universal brain and human reference samples, HCT116/PC3/MCF7/SKOV/TOV112D cell lines, high grade and low grade ovarian cancer samples from *H. sapiens* (19, 39–41, 57–59)) or generated in this study (wild type samples from *S. cerevisiae*, brain sample from *M. musculus*). The generation and analysis of these TGIRT-Seq datasets is described in the following subsections. The snoRNAs in *H. sapiens* were retrieved from snoDB (v2.0) (60) whereas the snoRNAs from the other species were extracted from the Ensembl annotations (Animals v108 and Fungi v55). Per species, snoRNAs were considered to be expressed if their abundance was greater than 1 transcript per million (TPM) in at least one average condition (mean across replicates of a same condition, except for human plasma TGIRT-Seq samples, for which we stringently required at least half of the plasma samples at an abundance >1 TPM in order to account for greater variability in these samples); otherwise, they were considered as C/D box snoRNA pseudogenes.

Negative examples included in the initial dataset were constituted of four types of midsize noncoding (mnc)RNAs (transfer RNAs (tRNAs), H/ACA box snoRNAs, small nuclear RNAs (snRNAs) and pre-micro RNAs (pre-miRNAs)), shuffled version of annotated C/D box snoRNA sequences as well as randomly selected 194 nt windows in exons, introns and intergenic regions that did not overlap with any annotated C/D box snoRNA. These negative examples were randomly sampled amongst the genome of species for which we collected C/D box snoRNAs and for which sufficient reliable annotation existed. More specifically, mncRNAs were sampled from the species present in the RNAcentral database (61), whereas exons, introns and intergenic regions were sampled based on the Ensembl (31) GTF annotation and genome FASTA files for the different species (genome and annotation file versions: Protists v55, Animals v108, Plants v55, Fungi v55).

In addition, the following TGIRT-Seq datasets were retrieved from the literature and processed as described in the following subsections for the cross-species inference results to identify relevant C/D box snoRNA predictions (thereby not included in the training of SnoBIRD): wild type and MLP1 immunoprecipitation samples from *T. thermophila* (62), developmental stage samples (egg (0 hour post fertilisation (hpf)), 256-cells (2.5 hpf), 1000-cells (3 hpf), sphere (4 hpf), shield (6 hpf) and bud (10 hpf)) from wild type *Danio rerio* (63) and developmental stage samples (initial infection with complete or amino acids-depleted medium, ring, trophozoites and schizonts stages) from the NF54 *Plasmodium falciparum* strain (64).

### Dataset filtering, data augmentation strategy and dataset split

Positive examples were filtered using the following criteria: exact duplicate window sequences were removed (i.e. snoRNA copies with the same genomic context) and snoRNAs part of an Rfam (65) family with more than 100 members were reduced to 100 members maximum (by randomly removing snoRNA members in order to reduce overrepresentation of some snoRNA sequences). Moreover, based on the length distribution of collected C/D box snoRNAs across species (**Supplementary Figure S1A**), the maximal length for a snoRNA to be considered in the initial dataset was set at 164 nt (since it included 95 % of all collected examples, without shifting too much the length towards a handful of longer nonrepresentative snoRNAs). C/D box snoRNAs of length greater than 164 nt were therefore excluded from all downstream analyses. The chosen window size that is fed to SnoBIRD was set at 194 nt, thereby including the snoRNA length threshold plus a flanking 15 nt on both sides of the snoRNA. These flanking nucleotides were added as it was previously shown that one of the most important determinant of snoRNA expression is the formation of a terminal stem using nucleotides that are internal to and flanking the snoRNA (24). As the 194 nt fixed window length was set, the sequence of examples in the initial dataset was extended to reach that desired length: C/D box snoRNAs and other mncRNA negative example sequences were extended uniformly on both sides of the given gene using the surrounding genomic nucleotides, thereby creating a 194 nt window containing a centered example (**Supplementary Figure S1B**). Random exonic, intronic and intergenic regions (i.e. the other negative examples) were sampled directly as 194 nt windows.

To increase SnoBIRD training inference, positive examples (expressed C/D box snoRNAs and snoRNA pseudogenes) were data augmented through the creation of synthetic new examples by shifting either left or right the window that is initially centered around the snoRNA (**Supplementary Figure S1C**). Two different data augmentation strategies were used, one for each of SnoBIRD’s model (the first model being used for the C/D box snoRNA identification step, the second being used for the refinement step, i.e. the prediction of expressed C/D box snoRNA vs snoRNA pseudogene). For SnoBIRD’s first model dataset, each positive example was data augmented 10 times (five 1-nt shift to the left, and five 1-nt shift to the right of the original window). The overall ratio of negative to positive examples in SnoBIRD’s first model dataset was set at ∼5:1 to reflect the overrepresentation of non-snoRNA sequences that SnoBIRD encounters when running across a whole genome. For SnoBIRD’s second model dataset, each expressed C/D box snoRNA example was data augmented 10 times, whereas each C/D box snoRNA pseudogene was data augmented 30 times (fifteen 1-nt shift to the left, and fifteen 1-nt shift to the right of the original window) to create an equal number of examples of both classes. Both datasets were split into three independent parts: the tuning, training and test sets (respectively, 10%, 70% and 20 % of all examples). The tuning set was created to tune the hyperparameters of the models, whereas the training and test sets were respectively created to train the models and evaluate their performance on examples that were never encountered during training. SnoRNAs of a same Rfam clan and Rfam family were constrained into one set only in order to reduce example redundancy across sets and thereby limit the chances of overfitting. Manual stratification was performed when dispatching negative examples across sets to ensure a consistent distribution of species and negative types across the sets. The datasets used to tune/train/test the first and second model used by SnoBIRD are accessible on Zenodo at https://zenodo.org/records/14927289, respectively containing expressed C/D box snoRNAs, snoRNA pseudogenes and negative examples for the first model, and expressed C/D box snoRNAs and snoRNA pseudogenes for the second model.

### Hyperparameter tuning, training and testing of SnoBIRD

SnoBIRD was built by fine-tuning DNABERT (43), i.e. by optimizing and training a classification layer on top of the pretrained BERT model using the *transformers*, *scikit-learn* and *optuna* libraries (66–68). This fine-tuning was done in parallel for the two DNABERT models that SnoBIRD employs: the first one predicts if there is a C/D box snoRNA gene (in the general sense) in the input sequence, and the second one predicts if the C/D box snoRNA gene is expressed or is a pseudogene. For each BERT model, hyperparameter tuning was performed using the grid search algorithm to find the optimal learning rate and batch size that maximized the F1-score over a stratified 3-fold cross-validation strategy after 4 epochs of training (as recommended in (42)) with their respective tuning set. Using the optimal hyperparameters, each BERT model was next trained over 4 epochs using their respective training set to fit the parameters of the classification layer. To stabilize the learning process, a learning rate schedule was implemented for both models during training by decreasing linearly the learning rate to a tenth of its initial value over the training epochs. After training, the performance of the two fine-tuned BERT models was assessed on their respective test set. SnoBIRD was also later run on the following genomes and chromosome sequence (with default parameters unless specified): *Schizosaccharomyces pombe* (with -cs 2), *H. sapiens* (with -s 1), *D. melanogaster* (with -cs 10 -G V100), *T. thermophila* (with -cs 10 -G V100), *G. gallus* (with -cs 10 -G V100), *M. mulatta* (with -cs 10 -G V100) and chr1 of *H. sapiens* (with -cs 10 -G V100), *P. falciparum* and *D. rerio* (with -cs 100). SnoBIRD was run using one NVIDIA A100 graphics processing units (GPU) and 2 central processing units (CPU) cores for the *S. pombe*, *H. sapiens* and *P. falciparum* genomes; the remaining genomes were predicted on by SnoBIRD using one NVIDIA V100 GPU and 2 CPU cores.

### Implementation of the different SnoBIRD steps

SnoBIRD was written in python (v3.10.14) and implemented under a Snakemake workflow. SnoBIRD was elaborated so that it can take as input either a FASTA file (containing one or multiple chromosome (or smaller) sequences) or a BED file of regions of interest as well as a FASTA file containing the genome sequence (**Supplementary Figure S2**). SnoBIRD was designed to use a sliding window approach to scan the input sequence with more or less sequence redundancy between the different overlapping predicted windows. To define what default step size should be used with SnoBIRD (i.e. the number of nucleotides employed to shift the sliding window; with a step size of 1 meaning all possible overlapping windows are predicted on, a step size of 2 meaning an overlapping window is skipped at every shift, etc.), a step size analysis was generated on *S. pombe* and *H. sapiens* genomes (**Supplementary Figure S3**). After using SnoBIRD with a step size of 1 on both species and thereby predicting on all possible 194 nt windows in these genomes, the impact of increasing the step size was mimicked by synthetically removing the relevant windows for each step size from 2 to 50 (i.e. the windows that would be skipped if SnoBIRD model was actually rerun using the given step size). The conservative step size of 5 was chosen as SnoBIRD’s default step size value, since for both species, it maximized the number of annotated C/D box snoRNAs predicted by SnoBIRD, while minimizing the total number of predictions (**Supplementary Figure S3**). This is by design a conservative threshold, and the user can easily increase the step size value to up to 10 without risking losing too many relevant predictions (using the -s option). The net impact of increasing the step size is an expected linear decrease in terms of prediction time (as the total number of windows to be predicted on decreases by a factor of *s*, where *s* is the step size).

After predicting on the sequences of interest with SnoBIRD’s first model (identification step), a merging strategy was implemented, serving the purpose of merging overlapping entries and selecting significant predictions (**Supplementary Figure S2**). The following filters were included in this merging strategy in this order: 1) keep predicted windows that have a probability greater than or equal to the defined *p1* threshold (default: 0.999, or 0.9 for BED input); 2) merge and keep blocks of at least *w* consecutive positive windows (default: 10); 3) return the centered 194 nt window within each block if it is also predicted positively. To define the snoRNA start and end within the resulting positive windows, a Shapley Additive Explanation (SHAP)-based approach was implemented using the *SHAP* package (69) set with the partition algorithm and max_evals=50. Briefly, SHAP values are computed for each nucleotide (average (mean) SHAP value across all k-mers containing the given nucleotide) in the input window. Peaks of high SHAP values are identified along the window sequence using the savgol_filter and find_peaks function from *scipy* (v1.13.0). C (RTGATGA) and D (CTGA) motifs are searched for amongst the peaks that fit within the respective range of C and D motif location (C and D motifs being located respectively before and after the half-window). The best C-D pair limiting the number of mutations with regards to their consensus sequence is then chosen. The best C’ and D’ box are then identified by selecting the best C’-D’ pair (i.e. limiting the number of mutations with regards to their consensus sequence) amongst all possible pairs (where C’ is downstream of D’) located between the previously identified C and D motifs. The predicted snoRNA start and end are then defined as the fifth nucleotide upstream of the C box and the fifth nucleotide downstream of the D box, respectively, based on the distribution of start/end positions with regards to the C and D motifs of all annotated C/D box snoRNAs present in the initial collected dataset (**Supplementary Figure S4**).

The following refinement step was implemented to predict if the C/D box snoRNAs identified by the first model are expressed or pseudogenes. Briefly, SnoBIRD’s second model is run on the identified windows and returns a probability associated with either the expressed C/D or pseudogene prediction. The predictions are then separated in the two following categories based on the probability threshold *p2* (default: 0.999): high confidence (if probability ≥ *p2*) or low confidence prediction (probability < *p2*). The high and low confidence predictions are then filtered differently based on known snoRNA expression determinants, namely the C and D box score (Hamming distance, i.e. the number of mutations compared to their consensus sequence), the cumulative box score (sum of Hamming distances of the C, D, C’ and D’ boxes), the terminal stem score (terminal stem length multiplied by the terminal stem secondary structure stability computed with RNAcofold (from *ViennaRNA* v.2.7.0 (70)), as previously described in (24)) and the structural score (snoRNA secondary structure stability computed with RNAfold normalized by the snoRNA length). Based on feature distributions of expressed C/D box snoRNAs and snoRNA pseudogenes shown in the third main figure, different threshold values were defined as favorable (or not) for snoRNA expression and applied to the low and high confidence predictions. Namely, the low confidence predictions were subject to the following filters in this order: 1) the prediction is considered as a snoRNA pseudogene if its C box score ≥ 2 and its D box score ≥1; 2) the prediction is considered as expressed if its terminal stem score ≤ -25 kcal*nt/mol; 3) the prediction is considered as a snoRNA pseudogene if less than 2 features are favorable for its expression. For high confidence predictions of snoRNA pseudogenes, the prediction is overturned (thereby ultimately considered as an expressed C/D box snoRNA) if at least 4 of its features are favorable for its expression. Overall, by design, the refinement step was elaborated to favor expressed C/D box snoRNA prediction over snoRNA pseudogene prediction (in order to limit false negatives of the expressed C/D box snoRNA class, i.e. expressed C/D box snoRNAs that would be dismissed as pseudogenes, and by the same token increase precision on the snoRNA pseudogene class). By default, SnoBIRD was implemented so that both models are run, but using the --first_model_only option, the user can decide to run only the first model and obtain C/D box snoRNA predictions in the general sense (**Supplementary Figure S2**). In addition, although SnoBIRD can run on CPUs alone, its standard recommended usage is to be run on GPUs where its overall speed is vastly accelerated, because of its utilization of BERT models and SHAP value computations. Finally, SnoBIRD was designed to output the predictions in any of the following format: FASTA, BED, GTF or tab-separated value (TSV) files.

### Other tool usage

To compare SnoBIRD’s performance with existing tools, Snoscan, Snoreport2 and Infernal_rfam were run on the test set of the first model as well as on the whole FASTA sequence of *S. pombe* and *H. sapiens* genomes using 2 CPU cores (no GPU was provided to any of the tools as by design, none of these tools can benefit from GPU acceleration). As the running time increased heavily with genome size for all these tools, their job runtime was limited at maximum 60 h when they ran for the time comparison analysis in order to limit unnecessary energy consumption and CO_2_ emission. Snoscan was run using default parameters, and the following rRNA sequences from the NCBI Gene database were fed as mandatory input for predicting in the different species present in the test set and/or for the species whole genome predictions: *S. pombe* (Z19578.1), *S. cerevisiae* (LC756999.1), *N. crassa* (FJ360521.1), *C. albicans* (AACQ01000295.1), *A. fumigatus* (KC119199.1), *O. tauri* (Y15814.1, AY586729.1), *O. sativa* (LC086814.1), *A. thaliana* (X52320.1), *O. anatinus* (AJ311679.1, XR_003764684.1), *T. thermophila* (X54512.1), *G. lamblia* (XR_005248692.1, XR_005248693.1, 7PWO_1), *L. major* (XR_002460811.1, OP829811.1), *D. discoideum* (AM168071.1, HQ141459.1, JF931128.1), *C. elegans* (X03680.1), *D. melanogaster* (X15707.1), *M. musculus* (NR_003279.1, NR_003278.3, NR_003280.2), *G. gallus* (KT445934.2), *M. mulatta* (KX061890.1) and *H. sapiens* (NR_146144.1). Snoreport2 was run with the following options (-CD --positives) on the input sequences as well as on the reverse complement of these sequences which needed to be manually generated since Snoreport2 predicts only on the + strand. Infernal_rfam was run using Infernal’s cmscan submodule with Rfam covariance models v15.0 with default parameters except the following added options: -- cut_ga –rfam --nohmmonly. The output of all these tools was filtered and formatted using custom python scripts to extract pertinent information for comparison with SnoBIRD.

Clustal omega was run with default parameters on its web server (71) to multi-align SnoBIRD predicted C/D box snoRNA sequences within the *SF3B3* /*Sf3b3* locus in *D. melanogaster*, *D. rerio*, *G. gallus* and *M. mulatta*. The resulting Newick tree file was then used to generate the phylogenetic tree.

### Filtering of predictions

After applying SnoBIRD (and the other tools) across the diverse genomes mentioned above, predictions were filtered using *pybedtools* (v0.10.0) (72) to find which predictions overlapped annotated C/D box snoRNAs as well as expressed regions in the genomecov-generated bedGraph files (see the section “TGIRT-Seq library preparation and analysis” below for more details). The annotated C/D box snoRNAs were obtained from Ensembl (31) GTF files (Protists v55, Animals v108, Plants v55 and Fungi v55) supplemented with RNAcentral (61) and Rfam (65) snoRNAs. Different filters were used to filter real expressed predictions (potential new C/D box snoRNAs) using custom python scripts: a prediction had to be covered by reads on the same strand on at least 50 % of its length, with more than 5 reads on average (mean), and the regions flanking the prediction (20 nt on both sides) should not have more than a third of the average number of reads covering the prediction. Following these filters, each potential candidate was manually inspected to remove predictions overlapping expressed exons of the same size and to filter out predictions with low uneven coverage, thereby constituting the final candidate snoRNAs represented in the Venn diagrams of relevant figures. As the *D. rerio* TGIRT-Seq datasets that we used to identify expressed predictions were size-selected for RNAs ≤ 100 nt (63), a maximal 100 nt length filter was applied to *D. rerio* (on top of the 164 nt length filter) to remove annotated or predicted snoRNAs that exceeded that length.

### xxFigure generation

All figures were generated with custom python scripts using *matplotlib* (v3.8.4), *seaborn* (v0.13.2), *pandas* (v2.2.2) or Integrative Genome Viewer (IGV; v2.4.18) (73) exported images.

### Statistical analyses

The following equations were used to calculate the different scores included in this study:

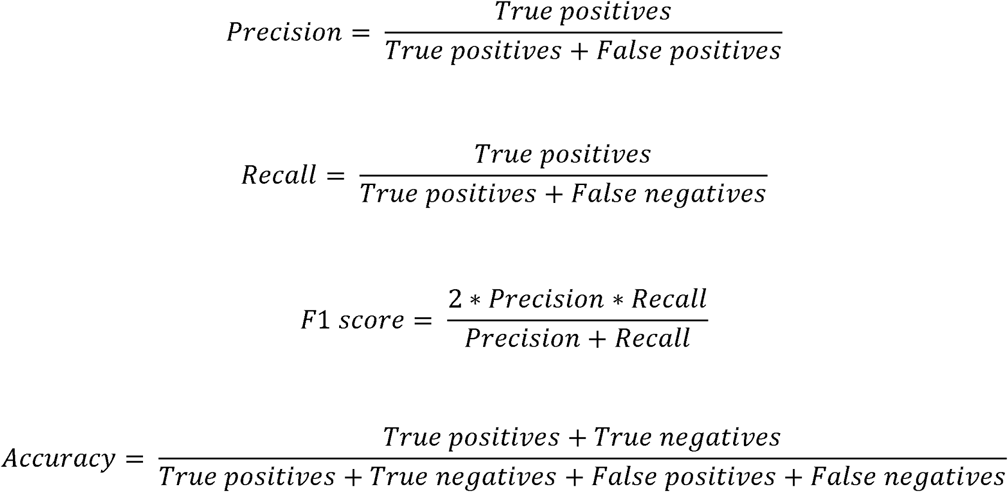

### Environmental impact

The training, optimization and inference steps with SnoBIRD, as well as other high-throughput analyses, were run on different high performance computing clusters of the Digital Research

Alliance of Canada (DRAC). The total environmental impact of the whole study is estimated at 0.821 kg of CO_2_ emission based on CPU and GPU usage across the DRAC clusters (information retrieved from the DRAC cluster-specific portals).

### Fission yeast culture for RNA coimmunoprecipitation (RIP)

The following *S. pombe* strains were used in order to perform RIP assays: wild type (WT) untagged (ID: FBY106; genotype: h+ ade6-M216 leu1-32 ura4-D18 his3-D1 (74)) and NOP58- Tandem Affinity Purification (TAP) (ID: FBY2417; genotype: h+ ade6-M216 leu1-32 ura4-D18 his3-D1 Nop58-TAP::natMX6). The addition of C-terminal TAP tag at the endogenous nop58 locus was performed by PCR-mediated gene targeting using the lithium acetate method for yeast transformation, as previously described (75). Proper gene targeting was confirmed by Western blotting. Fission yeast cells were routinely grown at 30°C in Yeast Extract Medium Supplemented (YES) with adenine, uracil, histidine, and leucine. For each condition, 50□mL of yeast cultures were grown to an OD600 of ∼0.5 at 30°C in YES and frozen in liquid nitrogen. Cell pellets from 50 mL cultures were thawed and resuspended in 500□μL of cold lysis buffer (50□mM Tris-HCl [pH 7.5], 140□mM NaCl, 2 mM MgCl2, 1% Triton X-100 and 1 mM dithiothreitol [DTT]) that was supplemented with a protease inhibitor cocktail (1 mM PMSF, 1X PLAAC, 1X complete protease inhibitor cocktail [Roche]) and RNase inhibitor (40 U/mL). Cells were disrupted vigorously with glass beads in a FastPrep-24™ (MP biomedical) for 3 cycles of 30□s at 6.5□m/s. 150□μL of cold lysis buffer with proteases and RNase inhibitors was then added to increase the volume, and samples were sonicated for 3 cycles of 5□s at 20% amplitude with a Branson digital sonifier. 50 μL of sonicated lysate (10% of IP volume) was kept for total RNA extraction (input fraction) and 15 μL was used for verification of IP efficiency by western blot. 500 μL of sonicated lysates were then incubated for 2 hours at 4°C with 50□μL of IgG Dynabeads (Life Technologies, 11041) in the case of TAP-tagged protein. For the anti-fibrillarin purification, beads were coupled with fibrillarin (ab4566, abcam) or flag (as a negative control) antibodies. Beads were then washed three times with cold lysis buffer supplemented with proteases and RNase inhibitors. After washing steps, beads were resuspended in 150 μL of lysis buffer and 15 μL of bead slurry was kept for western blot analysis. RNA was extracted using TRI Reagent (Sigma-Aldrich, T9424). RNA pellets were resuspended in 20 µL of RNase-free water for the input sample and 10 μL for the IP sample. 1 μL of diluted input RNA and the total of immunoprecipitated RNA was treated with 1 unit of RNase-free DNase RQ1 (Promega, M6101) for 30 min at 37°C and inactivated with 1 µL of 25 mM EDTA for 10 min at 65°C. Reverse transcription reactions were performed in 20 µL using random hexamers and the SuperScript III reverse transcriptase (Invitrogen). Quantitative PCR (qPCR) reactions were performed in triplicates on a LightCycler 96 system (Roche) in a final volume of 15 µL using 6 µL from a 1:20 dilution of each cDNA, 0.15 µM of forward and reverse primers, and 7.5 µL of the 2×PerfeCTa SYBR Green SuperMix from Quantabio. The primers used in the qPCR experiments are listed in **Supplementary Table S1**.

### Fission yeast culture, RNA extraction and size-selection for RNA-Sequencing (RNA-Seq)

Wild type yAS99 *S. pombe* cultures were grown to mid-log phase in 25 mL of YES media at 32°C. Cultures were pelleted at 4°C and 1900 x g for 10 min, and pellets were washed with 25 mL of sterile ddH_2_O, then re-pelleted at the previous conditions. Pellets were resuspended in 500 µL of nuclease-free H_2_O and transferred to sterile microfuge tubes. Cells were re-pelleted using the previous conditions and pellets were subsequently washed with 250 µL of complete RNA Extraction Buffer A (50 mM NaOAc pH 5.2, 10 mM EDTA pH 8.0, 1% SDS). 750 µL of warm Buffer A-saturated acid phenol was then added to each sample. The samples were vortexed to resuspend the pellet in the buffers and were incubated at 65°C for 4 min, vortexing occasionally. The samples were subsequently centrifuged at maximum speed for 3 min and the aqueous layer was transferred to a new sterile microfuge tube. An additional 250 µL of complete RNA Extraction Buffer A was added to the initial tubes with cells and a second round of RNA extraction was performed as described above, adding the aqueous layer to the corresponding tubes and discarding the phenol layer. Phenol-chloroform RNA extraction and precipitation was then performed on the aqueous samples, by adding an equal volume of phenol:chloroform:isoamyl alcohol (25:24:1) to each sample and vortexing until thoroughly mixed. The aqueous layer obtained from centrifugation at 14,000 x g for 10 min was transferred to a sterile microfuge tube. To each aqueous sample, 2 μL of GlycoBlue coprecipitant (Invitrogen cat. #AM9515), 10% of the isolated sample volume of 3M NaOAc pH 5.2, and 2.5X the isolated sample volume of ice-cold 100% ethanol was added. RNA was left to precipitate overnight at - 80°C. Precipitated RNA was pelleted at 4°C and 14,000 x g for 10 minutes. Pellets were washed with 500 µL of ice-cold RNase-free 70% ethanol and re-pelleted at the previous conditions. Pellets were air-dried and resuspended in 7 µL of nuclease-free H_2_O and 7 µL 2X formamide dye (80% deionized formamide, 10 mM EDTA, 0.06% bromophenol blue, 0.06% xylene cyanol). RNA samples were heated at 95°C for 5 minutes and snap-cooled on ice prior to separation on a 10% denaturing urea gel (8 M urea, 10% acrylamide, 1X TBE) at 100 V until the xylene cyanol dye front reached the bottom of the gel. Following staining with EtBr in 1X TBE (1 µL/5 mL), the gel was visualized under UV light and RNAs were size selected (20-150 nucleotides) by cutting each lane between the bands corresponding to the 5S and 5.8S rRNAs and retaining the lower halves in microfuge tubes. RNA was eluted from the gel pieces by overnight rotation at 4°C in 150 mM NaOAc pH 5.2 and 50% phenol:chloroform:isoamyl alcohol (25:24:1). The aqueous layer obtained by centrifugation at 4°C and 20,000 x g for 10 minutes was precipitated, and precipitated RNA for all samples was pelleted, washed, and air-dried, as described above. Pellets were finally resuspended in 20 µL nuclease-free H_2_O.

### Budding yeast culture, RNA extraction and rRNA depletion for RNA-Seq

Wild type 10H3 *S. cerevisiae* (genotype: MATa his□200 leu2□0 ura3□0 lys2□0) cultures were grown in Yeast Extract Peptone Dextrose (YEPD) rich media at 26°C up to an OD600 between 0.6 and 0.8. 50 mL of culture were centrifuged for 1 min at 4°C and 6000 rpm. The pellet was resuspended with 10 mL sterile water at 4°C, then centrifuged a final time at 6000 rpm for 1 min. The supernatant was discarded, and the pellet was frozen at -80°C. The pellet was resuspended with 300 µL of LETS buffer (0.1M LiCl, 0.01M Na-EDTA pH 8.0, 0.01 M Tris-HCl pH 7.5, 0.2% SDS) and incubated for 2 min on ice. 0.7 ml of glass beads were added to the samples, to enable mechanical cell lysis, followed by 300 µL of phenol. Samples were vortexed 7 times for 30 s, with 30 s breaks on ice. Supernatant was collected and beads were washed with 150 µL of LETS buffer, before centrifuging the samples at 14,000 rpm for 3 min. The supernatant from the washed beads was added to the previously collected fraction, followed by 2 RNA extractions with phenol:chloroform:isoamyl alcohol (25:24:1) and one with chloroform alone. RNA was then precipitated with 3 volumes of 95% ethanol and 2% KOAc, stored at -80°C for 30 min, followed by centrifugation at 14,000 rpm for 10 min. The resulting pellet was then washed 2 times with 95% ethanol and air-dried. RNA pellet was resuspended in 100 µL of RNAse-free water and underwent DNAse I treatment using the RNeasy kit (QIAGEN). The DNAse-free RNA was then ribodepleted using the Yeast Ribo-Zero Gold rRNA Removal Kit (Illumina) as previously described (19). The size profile and concentration of the ribodepleted RNA were validated on an Agilent RNA ScreenTape (Agilent Technologies 5067-5576).

### Mouse brain RNA extraction for RNA-Seq

Brain samples were extracted from adult female BALB/c mice and directly frozen until RNA extraction. Samples were first cryo-ground into powder prior to total RNA extraction. Total RNA was extracted using TRIzol Reagent (Invitrogen) according to the manufacturer’s recommendations. Briefly, cryo-ground samples were resuspended in 1 mL of TRIZol Reagent and 200 μL of chloroform were then added followed by a thorough mix. Samples were incubated 2 min at 4°C before a 15 min centrifugation at 14,000 g and at 4°C. The aqueous phase was then collected and transferred to RNAse-free microtubes. 1.5 μL of Glycoblue carrier (Invitrogen) and 600 μL of propan-2-ol were added and samples were thoroughly vortexed and incubated 10 min at room temperature. RNA pellets were then collected after a 15 min centrifugation at 14,000 g and at 4°C, and washed with 75% ethanol. RNA pellets were air dried 10 min before being resuspended in RNAse-free water (Life Technologies). Nanodrop (Thermo Scientific) was used to quantify sample concentrations and evaluate purity ratios. The RNAse-free RNA was then ribodepleted using the Ribo-Zero Gold rRNA Removal Kit (Illumina) as previously described (19).

#### Size-selection for RNAs of M. mulatta and G. gallus samples

10 µg of total RNA from Universal Chicken (Biochain R4C34565-1) and Universal Monkey (Biochain R4534565-1) were dried and resuspended in 10µL formamide dye (94% deionized formamide, 0.05% Bromophenol blue and 0.05% xylene cyanol). RNA samples were heated at 95°C for 2 min and then snap-cooled on ice for an additional 5 min. The RNA was separated on a 10% denaturing urea gel (8M Urea, 10% acrylamide: bisacrylamide (19:1), 1X TBE at pH 8.3) at 100V in a Mini Trans-Blot Cell (Bio-Rad 153BR) until the bromophenol dye reached the bottom of the gel. Gels were stained with SYBR gold (Thermo Fisher Scientific S-11494) in 1X TBE buffer (1:5000). The RNAs were size selected (≤ 300 nucleotides) based on a size marker and split into two 0.6mL microtubes. The gel was crushed through a pierced hole of the 0.6mL microtube into a 1.7mL microtube. 500 µL of 150 mM NaOAc pH 5.1 was added to each microtube, which were vortexed and frozen at -80°C for 1h. 500 μL of 50% phenol:chloroform:isoamyl alcohol (25:24:1) was added, and RNA was eluted from the gel pieces by overnight rotation at 4°C. Samples were centrifuged at 4°C and 20,000 x g for 15 min, and the obtained aqueous phase was precipitated in 2% KOAc ethanol with 2 µL glycogen. RNA was pelleted and washed twice in 70% ethanol, and air dried. RNA was resuspended in 20 µL RNase/DNase-free H20. RNA size profile and concentration were validated on an Agilent RNA ScreenTape (Agilent Technologies 5067-5576).

### TGIRT-Seq library preparation and analysis

TGIRT-Seq libraries were prepared as previously described (40, 41) for the *S. pombe*, *S. cerevisiae*, *M. musculus*, *G. gallus* and *M. mulatta* samples by the RNomics platform at Université de Sherbrooke. Their subsequent analysis was performed using a snoRNA-sensitive detection pipeline that we previously published (19), with the addition of stranded bedGraph generation using genomecov from *bedtools* (v2.31.0) (76) with default parameters except for the following added flags: -bg -split.

## RESULTS

### Current annotations do not reflect the complexity of snoRNA repertoires, and no tool is currently suited to improve reliably these annotations

As mentioned above, it was recently highlighted that the latest snoRNA annotations are not systematically comprehensive amongst species, with clear gaps across several large eukaryote kingdoms and no consideration for the annotation of snoRNA pseudogenes (i.e. annotated snoRNAs that are not expressed) (8). As snoRNA pseudogenes exhibit characteristics that hinder their expression and which are very distinct from those of expressed C/D box snoRNAs (**Supplementary Figure S5**) (24), we sought to explore the extent to which snoRNA pseudogenes were distributed across a diverse panel of eukaryotes. By reanalyzing available and newly generated TGIRT-Seq datasets in human (*H. sapiens*), mouse (*M. musculus*), fruit fly (*D. melanogaster*) and budding yeast (*S. cerevisiae*), we observe a widespread distribution of pseudogenes amongst the currently annotated C/D box snoRNAs, varying from ∼65 % in human to 0 % in the unicellular eukaryote *S. cerevisiae* (**Figure 1A**). As none of the currently available snoRNA prediction tool (Snoscan, Snoreport2 and Infernal_rfam) can differentiate between expressed C/D box snoRNAs and snoRNA pseudogenes (**Table 1**), we sought to create a new tool that includes this relevant characteristic while maintaining beneficial features from the past approaches, eliminating or limiting their biases and enhancing the overall user-friendliness. SnoBIRD was thus developed such that a simple DNA sequence would be sufficient as input, thereby creating a tool which can theoretically identify any snoRNA, unlike Snoscan and Infernal_rfam that are limited, respectively, to discover canonical snoRNAs (i.e. with a rRNA target) and snoRNAs that are part of an Rfam family (**Table 1**). In terms of practical use, SnoBIRD was designed to predict by default on both strands of the input sequence (unlike Snoreport2’s predictions which are restricted to the positive strand), to be parallelized using GPU/CPU multiprocessing to reduce runtime (unlike Snoscan and Snoreport2 that are inherently not parallelizable), and to be reproducibly and easily installed via python’s virtualenv or Conda (unlike Snoreport2 which presents several dependency issues during installation) (**Table 1**). Furthermore, SnoBIRD’s output format was implemented to facilitate its downstream usage, which is not the case for the existing tools which often embed relevant information in a cumbersome format (**Table 1**). By default, a tab-separated file is produced by SnoBIRD to easily extract relevant information (e.g. genomic location, sequence, boxes, etc.), but additional FASTA, BED or GTF formats are also available as output.

**Figure 1.**
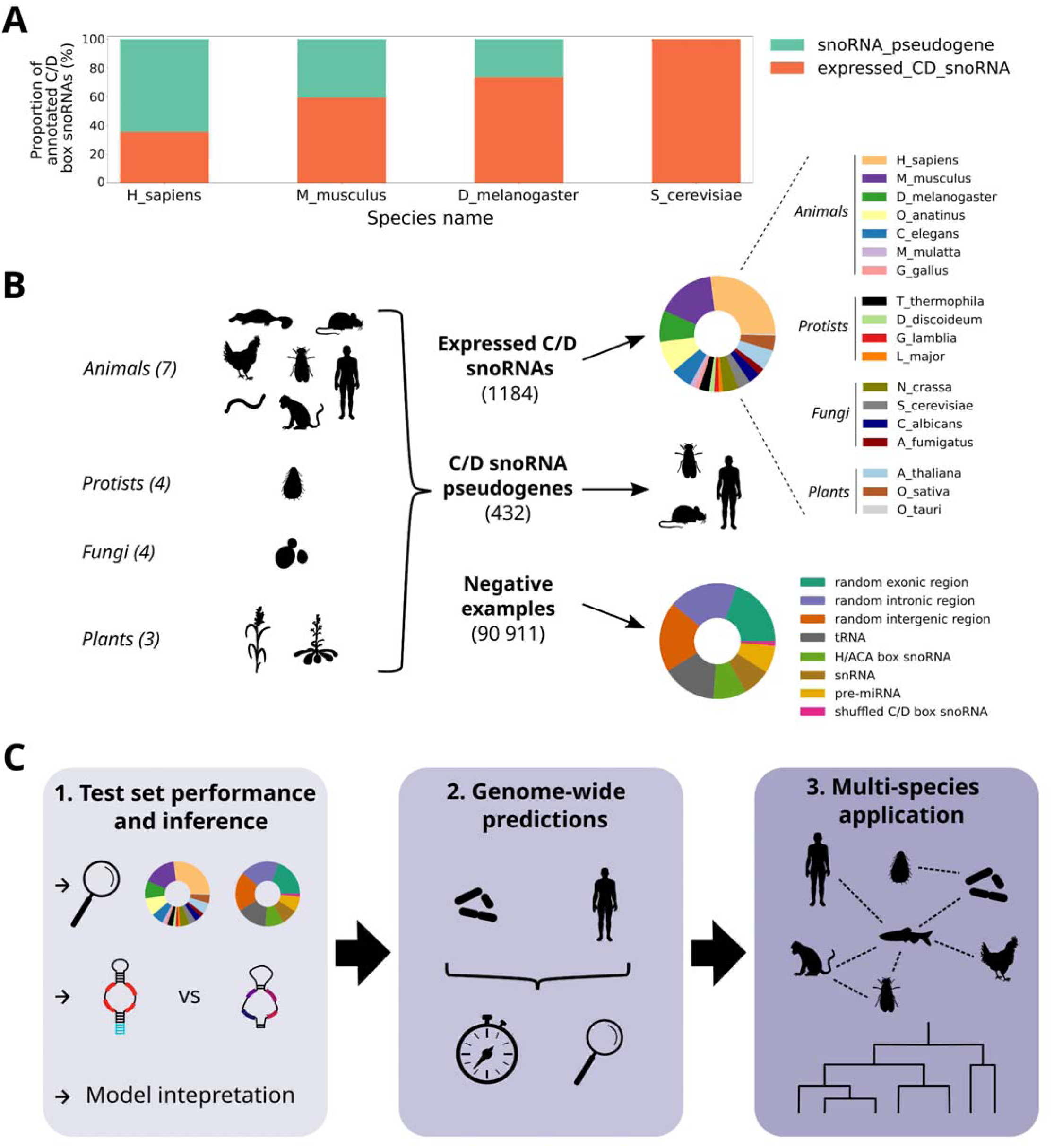
Current annotations do not reflect snoRNA complexity, and a new reliable method needs to be developed to alleviate this problem across all eukaryotes. **(A)** Proportion of the annotated C/D box snoRNAs that are expressed or pseudogenes based on TGIRT-Seq datasets in different eukaryotes and on the currently annotated C/D box snoRNA genes in Ensembl. **(B)** Composition of the initial dataset that was used to build SnoBIRD, a novel C/D box snoRNA predictor, in terms of species and gene biotype (for positive and negative examples, respectively). **(C)** 3-step strategy that was employed to compare SnoBIRD’s performance to existing methods. The first step consists of evaluating the performance of SnoBIRD on a held-out test set and compare its performance to existing tools. This step also includes snoRNA pseudogene prediction testing, which only SnoBIRD can perform as of today, and model interpretation to infer the learned patterns during training. The second step aims at evaluating SnoBIRD’s performance and that of its competitors in a genome-wide manner on a small and large genome (*Schizosaccharomyces pombe* and *Homo sapiens*, respectively), considering time efficiency and identification of known and novel C/D box snoRNAs. The third step consists of using SnoBIRD to predict C/D box snoRNAs across the genome of diverse eukaryotes with matched TGIRT-Seq samples, facilitating inference across species in terms of snoRNA evolution.

With these various desirable improvements in mind, we first gathered examples from the literature to form the initial dataset used for the training of SnoBIRD. This initial dataset was constituted of 1184 expressed C/D box snoRNAs and 432 C/D box snoRNA pseudogenes after filtering according to their length, with snoRNAs originating from more than 18 different species distributed across all eukaryote kingdoms (**Figure 1B**). As BERT models use preferentially fixed input sizes, only snoRNAs with a length ≤ 164 nt were comprised in the initial dataset to include as much examples as possible (∼95%) without shifting too much the size distribution towards outlier snoRNAs (**Supplementary Figures S1A** and **S6**). A fixed input size of 194 nt was therefore chosen to include minimally 30 nt surrounding the snoRNA (**Supplementary Figure S1B),** since these flanking regions (15 nt upstream and downstream) are known to form the terminal stem structure, which is an important determinant of C/D box snoRNA expression (23, 24, 77, 78). In addition to these positive examples, negative examples sampled from the same species were included in the initial dataset **(Figures 1B** and **S7**): other types of midsize noncoding RNAs (tRNAs, H/ACA box snoRNAs, snRNAs and pre-miRNAs), the shuffled sequence of C/D box snoRNAs and randomly selected regions in exons, introns and intergenic regions not overlapping with annotated C/D box snoRNAs. These regions were included in a greater proportion compared to positive examples to mimic the reality of genomes in which the vast majority of the sequences SnoBIRD will encounter will not include C/D box snoRNAs. Having assembled the initial dataset, we then devised a 3-step strategy to assess SnoBIRD’s performance following its optimization and training (**Figure 1C**). Firstly, the performance on C/D box snoRNA identification was compared between SnoBIRD’s and that of three other tools on a controlled held-out test set representative of all eukaryote kingdoms. As SnoBIRD is the only tool designed to identify snoRNA pseudogenes, its performance (but not that of other tools) was evaluated for the snoRNA pseudogene prediction task on the held-out test set. Following test set prediction, SnoBIRD’s model was interpreted using SHAP values (69) in order to infer and assess the reliability of the learned features during training. Secondly, SnoBIRD and the other tools were applied on two genomes with very different length (the fission yeast *S. pombe* and *H. sapiens*) to evaluate their scalability as well as the relevance of their predictions. Thirdly, SnoBIRD was applied genome-wide across several species with matched TGIRT-Seq datasets to identify novel C/D box snoRNAs and to highlight SnoBIRD’s potential in facilitating comparative evolutionary studies of snoRNA across species.

### SnoBIRD outperforms existing tools on a test set representative of all eukaryote kingdoms

Following the collection of the initial dataset, we used a data augmentation strategy to synthetically increase the representation of positive examples in our dataset (**Figures 2A** and **S1C**), a commonly adopted strategy known to improve training efficacy and overall performance of machine learning models (79). The data-augmented dataset was then split in three independent sets (respectively 10 %, 70 % and 20 % of all examples): the tuning set was used for hyperparameter tuning with a 3-fold cross-validation strategy, the training set was used for SnoBIRD’s training with a 10-fold cross-validation strategy and the held-out test set was used to assess SnoBIRD’s and the existing tools’ performance (**Figure 2A**). We next proceeded to SnoBIRD’s optimization and training as a way to fine-tune the BERT model (DNABERT from (43)) on the data-augmented dataset, reaching adequate performance as depicted by high F1-score over the epochs of training (**Supplementary Figure S8**). Intrinsically, SnoBIRD is designed to work in two main steps: the identification and the refinement steps, respectively to predict the presence of C/D box snoRNAs in the general sense and to predict if these snoRNAs are either expressed or pseudogenes (**Figure 2B**). Briefly, the input sequence is first split into 194 nt windows which are next converted into 6-mers (i.e. the DNABERT input format) and then transformed into dense vector embeddings. These embeddings are used by the first BERT model to perform the identification step (**Figure 2B**). Given a positive C/D box snoRNA prediction, the initial window is then fed to a second BERT model, which was trained to perform the specific refinement task, and passed through additional filters based on known snoRNA expression determinants (**Figure 2B**).

**Figure 2.**
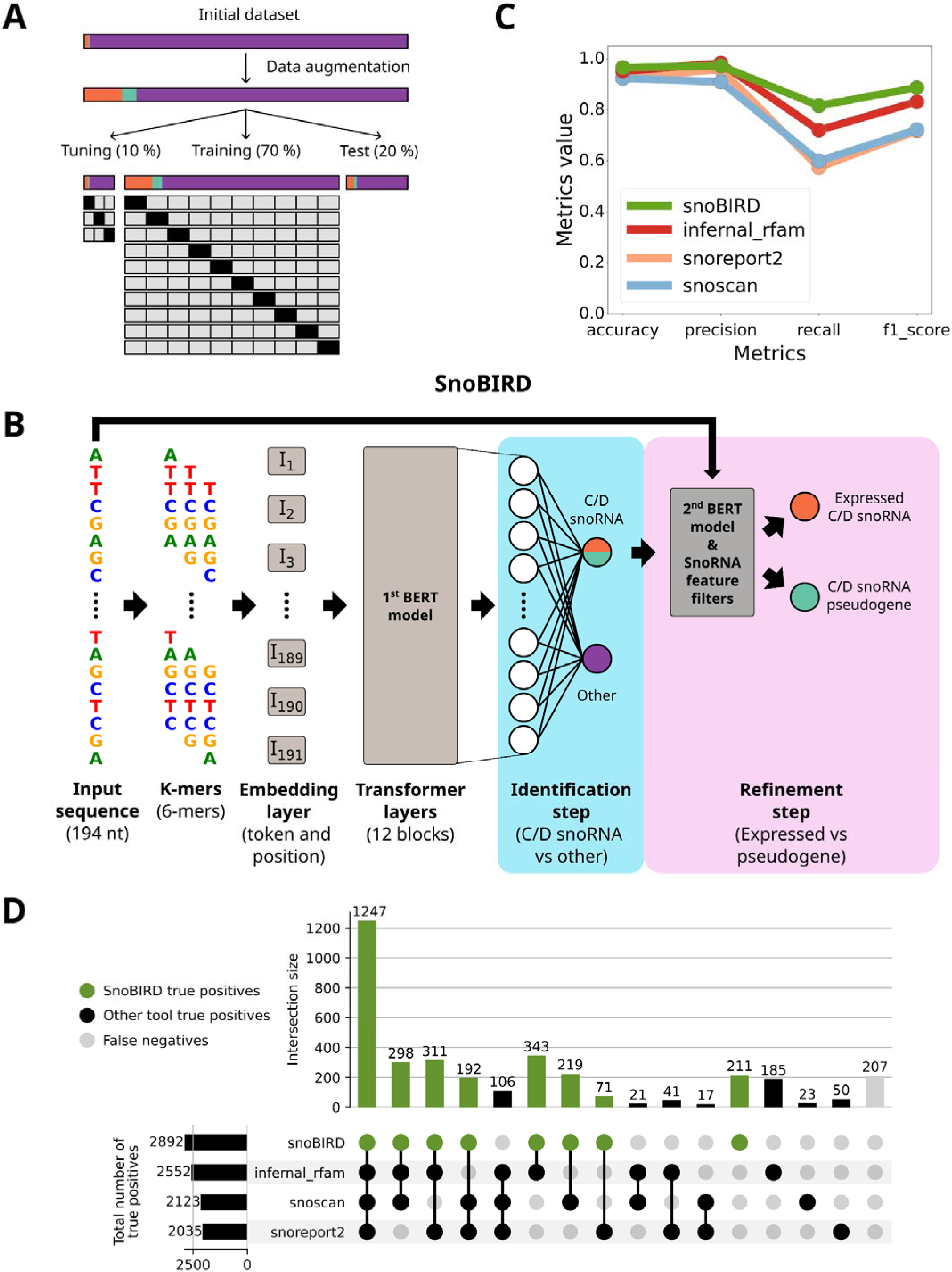
SnoBIRD outperforms existing tools in a controlled test set environment. **(A)** Schematic representing how the initial dataset composed of expressed C/D box snoRNA (orange), C/D box snoRNA pseudogenes (teal) and negative examples (violet) was data augmented for the snoRNA examples and then split into the tuning, training and test datasets of SnoBIRD’s first model (respectively 10, 70 and 20 % of the initial dataset). A stratified 3-fold cross-validation strategy was used to define the best hyperparameters for SnoBIRD (tuning set), whereas a stratified 10-fold cross-validation strategy was employed to train SnoBIRD (training set). The held-out test set was used to compare the performance of SnoBIRD and the other tools in a controlled environment. **(B)** Architecture of SnoBIRD which uses two models in respectively the identification and refinement steps (light blue and pink areas). First, the input sequence is converted into windows of 194 nt which are converted into k-mers (6 nt) and then converted into dense vectors by the embedding layer. These embeddings are then fed to the first model of SnoBIRD, a fine-tuned BERT model, which predicts, in the identification step, if the input sequence contains a C/D box snoRNA (in the general sense) or not. Finally, the refinement step consists in predicting if the C/D box snoRNA is expressed or a pseudogene, using a second fine-tuned BERT model combined with snoRNA feature filters (box conservation, global structure and terminal stem stabilities). **(C)** Line plot comparing different performance metrics (accuracy, precision, recall and F1-score) of SnoBIRD’s first model and other tools in predicting the C/D box snoRNAs present in the held-out test set. **(D)** Upset plot representing the intersection of test set predictions between the tools regarding the positive class (C/D box snoRNA). The green dots and vertical bars represent SnoBIRD’s true positives (a C/D box snoRNA predicted as such), the black dots and vertical bars represent the other tools’ true positives, whereas the gray dots and vertical bars represent false negatives (C/D box snoRNAs that were not predicted as such). The black horizontal bars represent the total number of true positive predictions per tool.

Interestingly, SnoBIRD outperforms all other tools on the held-out test set for the specific identification task based on the accuracy, recall and F1-score metrics, while being the second best after Infernal_rfam for the precision metrics (**Figure 2C**). These results also stand true when looking exclusively at the expressed C/D box snoRNAs present in the test set, whereas we observe an overall decrease in performance across tools when only considering snoRNA pseudogenes in the identification task, which is yet acceptable for SnoBIRD and Infernal_rfam but markedly low for Snoscan and Snoreport2 (**Supplementary Figure S9A-B**). Looking at the concordance of predictions between tools, we observe a great overlap between true positives (i.e. C/D box snoRNAs predicted as such), with SnoBIRD displaying the highest number of overall true positives (2892) and the highest number of true positives found by only one tool (211) (**Figure 2D**). In addition, the results were highly similar between tools for the true negative predictions (i.e. examples from the negative set not predicted as a C/D box snoRNA) (**Supplementary Figure S9C**). Finally, when considering the prediction errors of the tools across the different species present in the test set, SnoBIRD demonstrates a remarkably low number of prediction errors that seems unbiased across species, which is less the case for other tools (**Supplementary Figure S10**). For instance, Snoscan shows a high level of false positive predictions in *M. mulatta* (i.e. C/D box snoRNA predictions which are not snoRNAs in reality), whereas Infernal_rfam and Snoreport2 display a higher number of false negatives (actual C/D box snoRNAs that are missed by the tools) across protist and fungal species (**Supplementary Figure S10**, right half of the bar plots). Altogether, these results indicate that SnoBIRD is the most reliable tool to predict C/D box snoRNAs across all eukaryote kingdoms.

### SnoBIRD uses relevant biological signal to differentiate between expressed C/D box snoRNAs and snoRNA pseudogenes

Following SnoBIRD’s impressive performance on the test set, we sought to explore which features were learned during its training as a way to validate the biological relevance of its predictions. To do so, we interpreted SnoBIRD’s model by computing SHAP values (69) for all the C/D box snoRNAs accurately predicted from the initial dataset of 1616 snoRNAs (**Figure 3A**). Briefly, we computed an average SHAP value per nucleotide in the input sequence fed to SnoBIRD, thereby representing the importance of each nucleotide for the given prediction, with higher SHAP values correlating with higher importance (see Material and Methods for details). Strikingly, we observe that SnoBIRD tends to recognize two regions of importance for its predictions in the input sequences: one overlapping with the C box and the other overlapping with the D box (**Figure 3A**). These results not only indicate that SnoBIRD learned the C and D motifs that typify C/D box snoRNAs, but also that SnoBIRD can adjust the position of the found C and D boxes depending on the snoRNA length (as these important regions shift from the outer limits of the input window towards the center of the window as the snoRNA length decreases) (**Figure 3A**). Moreover, when clustering the snoRNAs by the resemblance of their SHAP profiles instead of sorting by their length, we notice that the C box region is in general more important than the D box region for the predictions (which is expected as the C motif is longer and thus more recognizable than the D motif), while SnoBIRD uses both the C and D box equally for some snoRNA predictions (**Supplementary Figure S11**). As these SHAP profiles clearly highlighted C and D box positions, we included the SHAP value computations as an integral part of the SnoBIRD pipeline in order to identify box motifs as well as the start and end of the predicted snoRNAs (**Supplementary Figure S2**). Indeed, as most snoRNA boundaries start 5 nt upstream of the C box and end 5 nt downstream of the D box (**Supplementary Figure S4**), we applied this same rule to the SnoBIRD predictions. When comparing SnoBIRD-defined snoRNA boundaries to those defined by the annotation for all C/D box snoRNAs accurately predicted in the initial dataset, we observe a strong overlap, suggesting that SnoBIRD reliably annotates the start and end of snoRNA predictions (**Supplementary Figure S12**).

**Figure 3.**
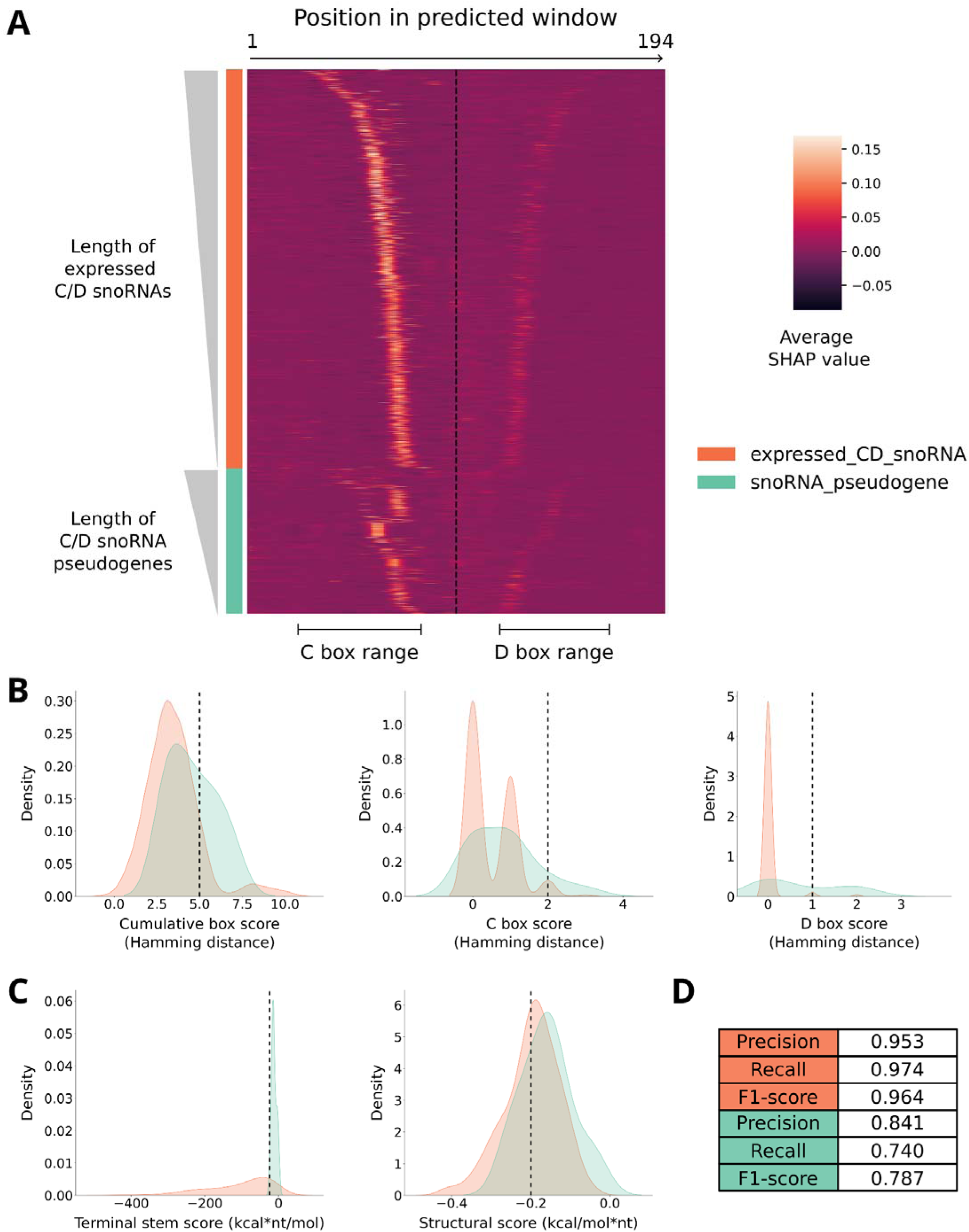
SnoBIRD uses biologically meaningful signal to predict and differentiate between expressed C/D box snoRNAs and snoRNA pseudogenes. **(A)** Heatmap showing the importance of each nucleotide (nt) in the input sequence (194 nt window) for the identification step of SnoBIRD (prediction of C/D box snoRNAs in the general sense). Stacked horizontal lines represent all the C/D box snoRNAs in the initial dataset that were predicted as such by SnoBIRD. Expressed C/D box snoRNAs were separated from snoRNA pseudogenes (represented by the left color bar), and both classes were sorted by increasing length of snoRNA within their fixed 194 nt window length (represented by the left grey gradient triangles). Vertical lines in the heatmap represent the first to last nucleotide (from left to right respectively) in the window encompassing the snoRNA. The dotted vertical line marks the center of the window. The nucleotide importance is color-shaded and based on its average SHAP value across all 6-mers containing that given nucleotide (the legend is on the right). The C box and D box range is underlined below the heatmap based on high SHAP values. **(B)** Density plots highlighting differences in snoRNA features between expressed C/D box snoRNAs (orange) and snoRNA pseudogenes (teal) in the test set based on the boxes that SnoBIRD found in the predicted snoRNAs. From left to right are shown the distributions of cumulative box score (sum of all mutations, i.e. Hamming distance, across the C, D, C’ and D’ boxes compared to their consensus sequence), C and D box scores (respectively the number of mutations, i.e. Hamming distance, in C and D boxes compared to their consensus sequence). Each vertical dotted line represents the chosen threshold value per feature used to filter expressed C/D box snoRNAs from snoRNA pseudogenes in SnoBIRD’s refinement step. **(C)** Same as (B), but the distributions represent the terminal stem score (predicted structural stability of the terminal stem multiplied by its length) and the structural score (predicted whole snoRNA structure stability normalized by the snoRNA length). **(D)** Metrics showing SnoBIRD’s performance for its refinement step (prediction of expressed C/D box snoRNAs vs snoRNA pseudogenes) on test set examples of each class (shaded by the same color code). Only SnoBIRD’s performance is represented as no other tool has the capacity to perform this task.

As the SHAP profile distributions differed between expressed snoRNAs and pseudogenes, with snoRNA pseudogenes depicting more asymmetrical C and D regions with regards to the center of the input window (**Figure 3A**), we decided to train a second BERT model to differentiate between these two subclasses of snoRNAs. While this second model identified C and D box regions as important for the expressed C/D box snoRNAs (although in a fainter way than with the first BERT model), it lacked a clear biological signal for the snoRNA pseudogene class predictions (**Supplementary Figure S13**). Therefore, to improve overall performance and biological relevance of this refinement step, we decided to include additional filters based on features known to distinguish expressed snoRNAs from snoRNA pseudogenes (see Material and Methods for details). Based on the boxes and snoRNA limits found through the SHAP value computation step of SnoBIRD, we could calculate a cumulative box score (sum of all mutations across the C, D, C’ and D’ boxes) and individual box scores that were, as expected, higher for snoRNA pseudogenes than for expressed C/D box snoRNAs (**Figures 3B** and **S14**). Similarly, we computed a terminal stem score (length of the terminal stem multiplied by its structural stability) and a structural score (snoRNA structural stability normalized by its length) which were, as expected again, lower (and therefore more stable) across expressed C/D box snoRNAs compared to snoRNA pseudogenes (**Figure 3C**). Combining the second BERT model predictions with the additional feature filters, SnoBIRD achieved high performance on the expressed C/D box snoRNA class (F1-score=0.964) and respectable yet lower performance on the snoRNA pseudogene class (F1-score=0.787) (**Figure 3D**). These data underline that, by design, SnoBIRD was optimized to maximize the performance on the expressed C/D box snoRNA class as we argue that it is preferable to consider a snoRNA pseudogene as expressed than to misclassify an expressed C/D box snoRNA as a pseudogene (and thereby potentially lose its biological relevance). Taken together, these results emphasize that SnoBIRD accurately predicts C/D box snoRNAs and is the only tool capable of distinguishing expressed from snoRNA pseudogenes, all of this relying on solid and meaningful biological signal.

### Genome-wide comparisons underscore SnoBIRD’s enhanced scalability and reliability to identify known and novel C/D box snoRNAs

After assessing SnoBIRD’s performance in a controlled test set environment, we next wondered how scalable and reliable SnoBIRD would be when deployed across a whole genome. To investigate this, we applied SnoBIRD and the other existing predictors across two genomes of very different size: the fission yeast *S. pombe* (∼12 Mb) and the human genome (∼3000 Mb). Upon usage on a small genome, SnoBIRD’s overall runtime of ∼85 min is reasonable and similar to that of other tools (except for Snoscan which already displays a longer runtime of ∼218 min, which is more than twice of SnoBIRD’s runtime) (**Figure 4A**, upper panel). When applied on the large human genome, SnoBIRD really outperforms its competitors with an overall runtime of less than 13h, thanks to its high parallelizability capacity (**Figure 4A**, lower panel). Its closest competitor, Snoreport2, takes more than 3.5 days (>87 h) to complete its predictions, whereas Infernal_rfam and Snoscan take more than 12 and 51 days respectively. To validate if these gigantic runtimes were specific to the whole human genome sequence, we applied all these tools on genomes of increasing lengths (**Supplementary Figure S15**), making sure to cap the runtime at maximally 60 h to limit unnecessary energy consumption and carbon emission. Similar to the previous results, only SnoBIRD prediction time scales efficiently with genome size, with the other tools becoming unpractical and almost unusable with genome sizes greater than 500 Mb (**Supplementary Figure S15**). Interestingly, SnoBIRD’s cumulative runtime is most affected by the genome prediction step (identification step with the first BERT model) for genomes of smaller sizes, whereas the SHAP value computation accounts for most of the total runtime on genomes of greater size (**Supplementary Figure S16**). Overall, these results indicate that SnoBIRD, but not its competitors, was optimized to be run on sequences of any size in a reasonable amount of time.

**Figure 4.**
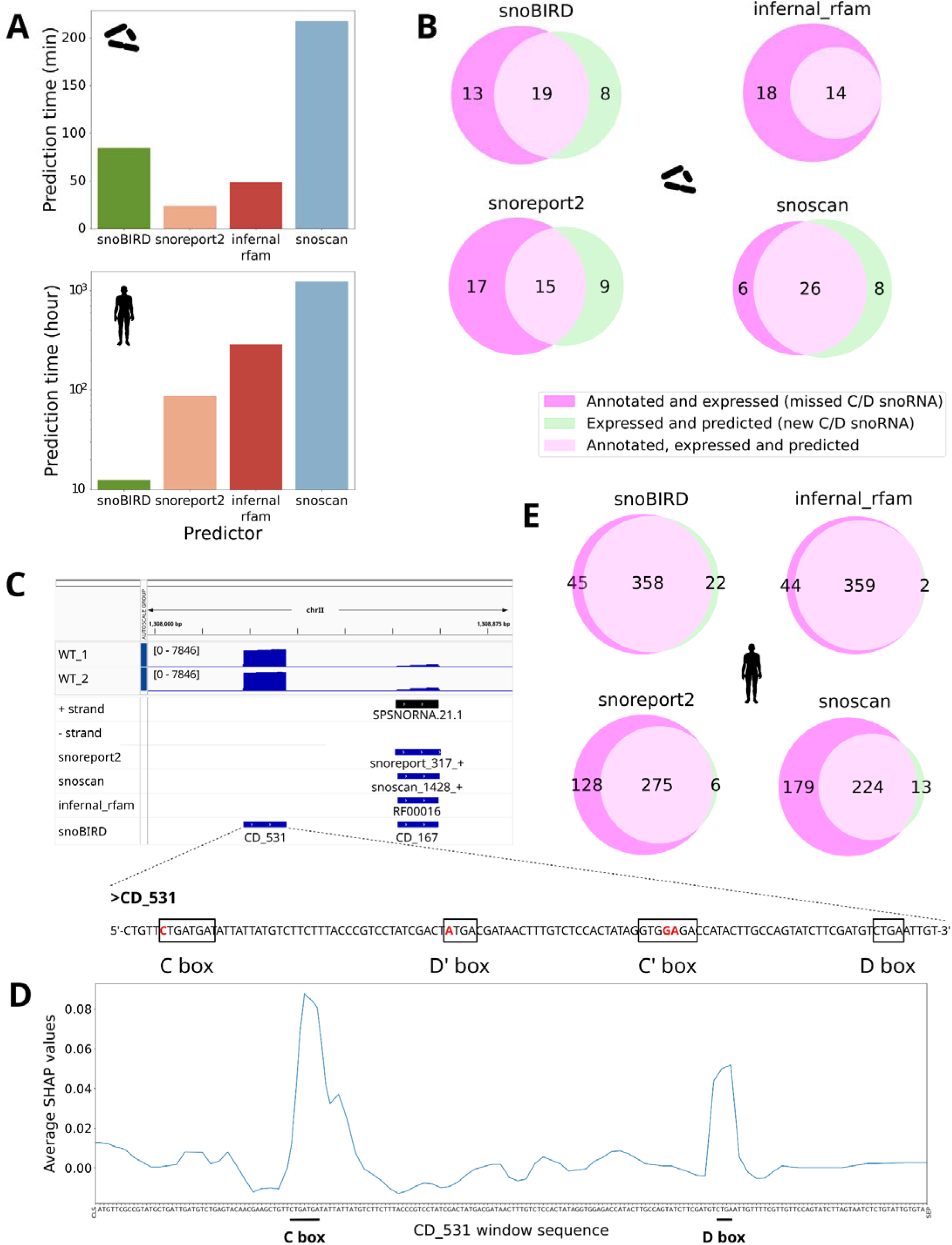
Genome-wide comparison between predictors underlines SnoBIRD’s efficiency and reliability to identify known and novel C/D box snoRNAs. **(A)** Bar plots displaying total runtime of the different C/D box snoRNA predictors on a small (top panel, *Schizosaccharomyces pombe*, genome size of ∼12 Mb) and a large genome (bottom panel, *Homo sapiens*, genome size of ∼3000Mb). **(B)** Venn diagrams showing the intersection between annotated or novel expressed C/D box snoRNAs and the different tools’ predictions in *S. pombe* (the expression is based on TGIRT-Seq samples). **(C)** Genome viewer screenshot depicting two TGIRT-Seq datasets of wild type (WT) *S. pombe* samples at the specific chrII:1,308,000-1,308,875 locus. The + and – strand tracks represent the currently annotated snoRNAs in *S. pombe*. The following tracks below represent the predictions made by each tool, including SnoBIRD as the lowest track. The sequence of the novel C/D box snoRNA only predicted by SnoBIRD (CD_531) is shown below the screenshot, with the SnoBIRD-inferred C/D’/C’/D motifs highlighted by black rectangles and mutated nucleotides (with regards to their consensus sequence) written in red. **(D)** Line plot showing the importance of each nucleotide in SnoBIRD’s prediction with regards to CD_531, the new snoRNA found in (C). The importance is assessed based on the average SHAP value (across k-mers containing the given nucleotide) along the 194 nt-long window sequence. The C and D boxes are underlined to highlight which peaks were used by SnoBIRD to define the presence of relevant C/D box snoRNA motifs. SHAP values were generated using the parameter max_evals=500 within the SHAP partition explainer for clarity purposes in this figure. **(E)** Same as in (B), but with *H. sapiens*.

Having predicted with the different tools on both *S. pombe* and *H. sapiens* genomes, we next sought to evaluate their potential in identifying known and novel expressed snoRNAs in these presumably well-annotated species. Out of the 32 annotated and expressed C/D box snoRNAs in *S. pombe*, SnoBIRD identified 19 of these (∼59%), which represents the second-best result after Snoscan (26/32 (81%) and an improvement over Snoreport2 and Infernal_rfam which found less than half of all annotated C/D box snoRNAs (**Figure 4B**). The higher recall for Snoscan is to be expected as it was trained specifically on yeast snoRNAs. Interestingly, SnoBIRD identified eight novel expressed C/D box snoRNA candidates, a similar number to what Snoreport2 and Snoscan found, whereas Infernal_rfam did not find any new C/D box snoRNA (**Figure 4B**). As a starting point, we decided to validate experimentally in *S. pombe* four of the eight SnoBIRD candidates using an RNA coimmunoprecipitation assay followed by qPCR to assess their binding to core proteins of the C/D box snoRNP. The four tested candidate snoRNAs showed a strong binding enrichment to both NOP58 and Fibrillarin (as observed with the positive controls, i.e. known C/D box snoRNAs), arguing in favor of these candidates being bona fide C/D box snoRNAs (**Supplementary Figure S17**). Further investigating these candidates, we focused on CD_531, a candidate which is amongst the most highly expressed ones in wild type *S. pombe* TGIRT-Seq samples (**Figures 4C**, top and **S18**). While being located in a potential snoRNA cluster close to the annotated *snoU14* (*SPSNORNA.21.1*), only SnoBIRD predicts this candidate as a C/D box snoRNA (**Figure 4C**, middle). This could be explained by the fact that its characteristic motifs display some alterations compared to their consensus sequence (**Figure 4C**, bottom), which could hinder motif search (a direct step implemented in all competing tools) whereas SnoBIRD, being agnostic to direct motif search, is more permissive in its identification of snoRNAs. Interestingly, when looking at the important regions in CD_531 sequence used by SnoBIRD to make its prediction, we observe that not only it finds the C and D motif regions as important but also extends beyond these regions downstream of the C box and upstream of the D box, highlighting potential important nucleotides that could serve to stabilize the snoRNA structure (**Figure 4D**).

Extending our analysis to human, we find that SnoBIRD identified 89 % (358/403) of all annotated and expressed C/D box snoRNAs, which is almost equal to Infernal_rfam (359/403), but considerably higher than Snoreport2 (275/403) and Snoscan (224/403) (**Figure 4E**). Notably, SnoBIRD identified 22 novel C/D box snoRNA candidates that are expressed in human TGIRT-Seq samples, which is more than all the other tools’ novel candidates combined (**Figure 4E**). Interestingly, the only two novel snoRNAs predicted by Infernal_rfam are also predicted by all the other tools, and overlap the longer annotated *SNORD91A* and *SNORD91B* (**Figures 4E** and **S19**). Strikingly, we can observe that SnoBIRD as well as Infernal_rfam and Snoreport2 nicely identify the most plausible start and end of these two snoRNAs with regards to read accumulation, which is not the case for Snoscan (as it frequently outputs multiple overlapping predictions with varying lengths) (**Supplementary Figure S19**). As other examples of reannotation, we can also observe that SnoBIRD helps in *S. pombe* to redefine the right start and end of snoRNAs that are either misannotated as longer snoRNAs, long noncoding RNAs or snRNAs in current annotations, which is not consistently the case for the other tools (**Supplementary Figure S20**). Another interesting case is SnoBIRD’s prediction CD_134 in *S. pombe* that is located close to the H/ACA box snoRNA *snR90* (*SPSNORNA.42.1*) (**Supplementary Figure S20D**). This prediction was previously identified as the C/D box snoRNA *snR80* (80), but was never added, to our knowledge, to the *S. pombe* gene annotation. Overall, these results underscore that SnoBIRD is also useful to correct and improve current snoRNA annotations.

As a final comparison between the different tools, we looked at the total number of predictions as well as their overlap with annotated genomic elements (**Supplementary Figure S21**). While SnoBIRD presents a relatively high number of total predictions (739 and 276 087 for *S. pombe* and *H. sapiens*, respectively), it still represents a major improvement compared to other snoRNA-specific approaches: Snoscan’s total predictions amount to more than twice those of SnoBIRD (1315 and 555 813 for *S. pombe* and *H. sapiens*) whereas Snoreport2’s totals amount to ∼1.5 time those of SnoBIRD (1118 and 360 790 for *S. pombe* and *H. sapiens*) (**Supplementary Figure S21**). To ensure a fair comparison for the total predictions, Infernal_rfam was treated in a different category as it is an annotation tool used by large annotation consortia such as Ensembl (31) and thereby not a tool made for the discovery of snoRNAs per say. This signifies that by design, Infernal_rfam is very stringent in its output predictions (as they need to fit very well within the covariance models, otherwise they are not returned as output), but this is to the detriment of identifying new snoRNAs, as depicted by its high precision and low recall (**Figure 2C**) as well as its very low number of newly identified snoRNAs (**Figure 4B,E**). To highlight potential biases in the genomic context of predicted snoRNAs for each tool, we then intersected all predictions with the Ensembl gene annotation for both *S. pombe* and *H. sapiens*, separating the genomic regions into exonic, intronic or intergenic (**Supplementary Figure S21**). We note that SnoBIRD, Snoscan and Snoreport2 can predict on all types of genomic regions, with a varying proportion that reflects each genome’s specificities (the *S. pombe* genome consisting largely of exons, whereas introns and intergenic regions dominate the human genome) (**Supplementary Figure S21**). Unexpectedly, we observe a strong bias for Infernal_rfam which does not predict any intergenic C/D box snoRNA in *S. pombe* and only 6 in human (although there are hundreds of intergenic snoRNAs annotated amongst these species (8, 60)) (**Supplementary Figure S21**). This is likely due to intergenic snoRNAs not being part of Rfam covariance models. Altogether, the results presented in this section indicate that SnoBIRD is the only C/D box snoRNA predictor that can be reliably applied to any genome and run in a reasonable amount of time to identify both expected and novel relevant snoRNA genes.

### Applying SnoBIRD across a broad panel of eukaryotes enables multi-species evolutionary studies

To prove SnoBIRD’s usefulness beyond single-species analyses, we applied it across a panel of varied eukaryotes with matched TGIRT-Seq datasets that we either generated within this study or that were recently published. In addition to *S. pombe* and *H. sapiens*, we predicted C/D box snoRNAs with SnoBIRD in the protists *T. thermophila* and *P. falciparum* as well as in the animals *D. melanogaster*, *D. rerio*, *G. gallus* and *M. mulatta*. In general, we observe that the proportion of predicted snoRNA pseudogenes increases and correlates with organismal complexity, apart from *P. falciparum* (**Figure 5A**). The proportion of expressed snoRNA predictions is higher than that of snoRNA pseudogene predictions for all species, which is an expected feature of SnoBIRD which favors by design the identification of expressed C/D box snoRNAs over that of snoRNA pseudogenes. Looking at the distribution of features known to influence snoRNA expression, we observe in these predictions similar trends across species to those found in the initial dataset: predicted expressed C/D box snoRNAs display lower cumulative, C and D box scores as well as lower terminal stem and structural scores compared to predicted snoRNA pseudogenes (compare **Figures 3B-C** and **S14** to **Supplementary Figure S22**). These observations highlight again the biological importance of differentiating between these two subclasses of snoRNAs using SnoBIRD.

**Figure 5.**
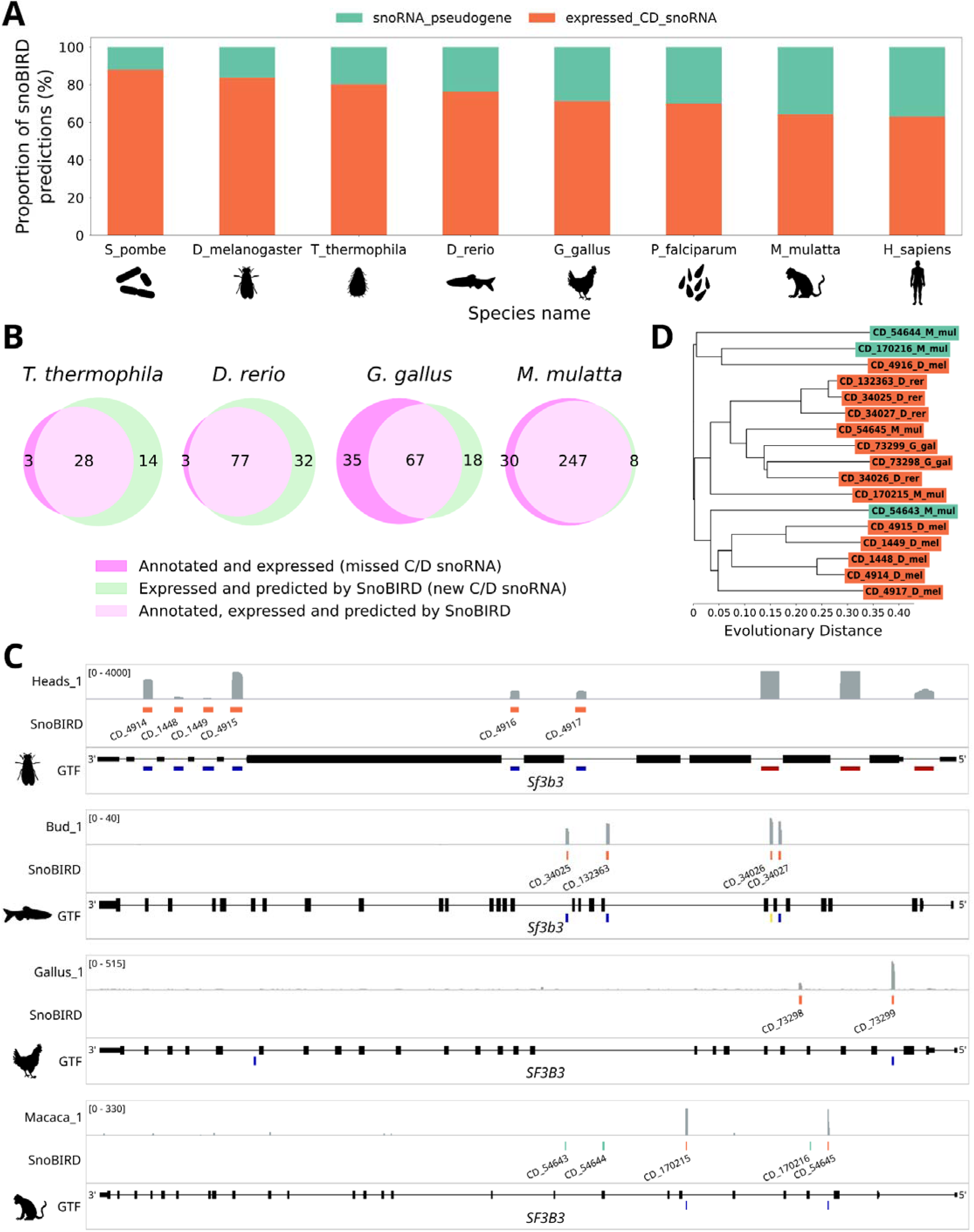
SnoBIRD enables multi-species inference, highlighting snoRNA divergent paths during evolution. **(A)** Stacked bar plot displaying the proportion of the different types of C/D box snoRNA predictions obtained after running SnoBIRD across a wide range of different eukaryote species. Species are ordered by increasing percentage of C/D box snoRNA pseudogene predictions. **(B)** Venn diagrams showing the intersection between annotated or novel expressed C/D box snoRNAs and SnoBIRD’s predictions across species (the expression is based on TGIRT-Seq samples). **(C)** Genomic view of the *SF3B3* gene locus across different species (fruit fly (*D. melanogaster*), zebrafish (*D. rerio*), chicken (*G. gallus*) and monkey (*M. mulatta*), from top to bottom respectively). Each subpanel shows in the GTF track the exons and introns of *SF3B3* as black rectangles and lines respectively, as well as annotated snoRNA genes as colored rectangles below (blue: C/D box snoRNAs; red: H/ACA box snoRNAs; yellow: snoRNA of unknown type). The SnoBIRD track shows SnoBIRD predictions, with orange and teal rectangles representing expressed C/D box snoRNAs and snoRNA pseudogenes, respectively. The top track in each subpanel shows read accumulation (bedGraph) along the locus for one representative sample per species. **(D)** Phylogenetic tree suggesting evolutionary distance between the C/D box snoRNAs predicted by SnoBIRD across the different species *SF3B3* genes (i.e. those represented in (C)). Expressed C/D box snoRNA and snoRNA pseudogene predictions are colored in orange and teal, respectively. Species names are abbreviated such as M_mul for *Macaca mulatta* after the prediction id for each leaf in the tree.

We next confirmed that SnoBIRD identified the majority of expressed and already annotated C/D box snoRNAs across these species, ranging from 65 % in *G. gallus* to more than 96 % in *D. rerio* (**Figures 5B** and **S23**). In addition to these known C/D box snoRNAs, SnoBIRD identified several new expressed C/D box snoRNA candidates across the different species (with the extreme case of 32 new candidates in *D. rerio*) (**Figure 5B**), highlighting its potential to supplement and expand current snoRNA annotations eukaryote-wide. These new predictions as well as those in *S. pombe* and *H. sapiens* were included in the **Supplementary Table S2** . We then assessed the genomic element overlapping these SnoBIRD predictions, also comparing side by side with the actual genomic proportion (**Supplementary Figure S24**). Interestingly, we see an enrichment for SnoBIRD predictions to overlap with introns across the genomes of animals (i.e. more intronic predictions than the actual intronic proportion of the genome), which is not observed in the other tested eukaryote kingdoms (**Supplementary Figure S24**). This suggests that SnoBIRD potentially learned that the typical snoRNA location in animals is in introns (8, 20), likely due to its larger 194 nt window encompassing not only the snoRNA but its genomic surroundings. Having predicted in all these different species, we could also note recurring overlap trends across species such as the peculiar location of several SnoBIRD predictions overlapping with 5.8S rRNA genes (**Supplementary Figure S25**).

As a final case study, we analyzed SnoBIRD predictions across a snoRNA host gene that is recurrently present amongst the different species covered in this study: *SF3B3* (or *Sf3b3* in *D. rerio* and *D. melanogaster*). This gene encodes a core protein component of the U2 small nuclear ribonucleoprotein complex (snRNP) and harbors a varying number of intron-embedded snoRNAs depending on the species (**Figures 5C** and **S26**). *D. melanogaster* ’s *Sf3b3* harbors six C/D box snoRNAs as well as an additional three H/ACA box snoRNAs (**Figure 5C**), which is more snoRNAs than any of the other species considered. To decipher if these *D. melanogaster* snoRNAs shared a possible common origin with *SF3B3* -hosted snoRNAs in other species, we applied SnoBIRD on all the considered species. Not only did SnoBIRD correctly identify the expressed C/D box snoRNAs annotated in the different species, but it also helped uncover one novel expressed C/D box snoRNA in *G. gallus* (CD_73298) as well as reannotate one previously annotated snoRNA of unknown box type in *D. rerio* (CD_34026) (**Figure 5C**). Additionally, SnoBIRD predicted the presence of three C/D box snoRNA pseudogenes in *M. mulatta*, which made us wonder if these could have a common origin with any of the *D. melanogaster* snoRNAs. Based on the sequence identity between SnoBIRD predictions across the different species, we constructed a phylogenetic tree to potentially recapitulate the evolutionary path of the different C/D box snoRNAs. Surprisingly, most *D. melanogaster* C/D box snoRNAs clustered together, along with CD_54643, a snoRNA pseudogene predicted in *M. mulatta* (**Figure 5D**, lower cluster). The other annotated C/D box snoRNA in *D. melanogaster* (predicted by SnoBIRD as CD_4916) also clustered but this time with the two other predicted snoRNA pseudogenes in *M. mulatta* (**Figure 5D**, top cluster). These relationships suggest some gene duplication events unique to *D. melanogaster*, and that these *M. mulatta* snoRNA pseudogenes might represent remnants of ancient snoRNAs only still functional in *D. melanogaster* and lost or no longer expressed in the three other organisms. Interestingly, the expressed C/D box snoRNAs from the remaining species all clustered in one group (**Figure 5D**, middle cluster), suggesting a more recent common origin for these snoRNAs differing from *D. melanogaster* ’s snoRNAs. In summary, these results demonstrate yet another useful application of SnoBIRD’s potential, this time unveiling possible evolutionary paths of snoRNAs across multiple species.

## DISCUSSION

In this study, we have highlighted major issues that current eukaryote snoRNA annotations present, such as their incompleteness with regards to the number of included snoRNAs as well as their widespread lack of specificity in terms of snoRNA type (i.e. C/D vs H/ACA box snoRNA not always specified) and snoRNA pseudogene annotation. To address this, we have designed SnoBIRD, a novel tool which predicts C/D box snoRNAs in any eukaryote genome and classifies them as expressed or pseudogenes. We have shown that not only does SnoBIRD outperform its competitor tools in a controlled test set representative of all eukaryote kingdoms, but that it does so using meaningful and reliable biological signal recognized within the input sequence as the characteristic C and D motifs. We have also demonstrated that SnoBIRD, but none of the other tools, can accurately distinguish between expressed C/D box snoRNAs and their pseudogenes, further refining current annotations. Applied on the whole genome of the fission yeast and human, we have underlined that only SnoBIRD is suited to identify genome-wide known and novel relevant candidate C/D box snoRNAs in a reasonable amount of time, while the other tools can take up to several weeks to generate their oftentimes incomplete predictions. Finally, we have applied SnoBIRD across a diverse panel of eukaryote species with matched TGIRT-Seq samples, allowing for the discovery of several new C/D box snoRNA candidates and enabling the study of snoRNA evolutionary paths across different species.

Although several C/D box snoRNA predictors have been released in the last decades (32–36), most of these tools have either become obsolete or carry questionable design choices, since they were often constrained by the availability of datasets and information at their time of conception. These drawbacks include the rigid search for the presence of motifs, structural features and guide sequence complementarity (in the case of Snoscan) which make these tools miss relevant C/D box snoRNAs (**Figures 4B-E**, **S17** and **S18**). While SnoBIRD benefits from greater prediction flexibility, other more generic approaches like Infernal_rfam that use covariance models are confined to only predicting snoRNAs which were previously identified in other species and characterized enough to be included in a given Rfam family. Albeit a tool with great value, Infernal_rfam thus remains more of a reannotation tool rather than a tool for discovery. It is also not adapted to all species (**Figures 4B** and **S10**), as less studied eukaryotes such as protists, fungi and plants are not as well represented in Rfam families, and seems biased against specific snoRNA location such as intergenic snoRNAs (**Supplementary Figure S21**).

Prioritizing user-friendliness during SnoBIRD’s elaboration, we have improved over past approaches in the fields of reproducible installation (automatic environment creation and model downloads), ease of use (expanded input and output types (**Supplementary Figure S2**), automated job submission, etc.) and scalability (parallelization strategy to work as well on small and large sequences (**Figures 4A** and **S15**)). Moreover, SnoBIRD can scan entire sequences (e.g., genes, chromosomes, genomes, etc.) using the FASTA format as input or only consider specific genomic regions suspected to harbor C/D box snoRNAs (e.g., regions enriched for the binding of an RBP following cross-linking and immunoprecipitation (CLIP) experiments, unannotated regions with blocks of read accumulation in RNA-Seq experiments, etc.) using the BED format as input (**Supplementary Figure S2**). This SnoBIRD-specific BED option enables its seamless integration in experimental pipelines designed for the discovery of novel genes or interactors, and it increases even more its prediction speed (as it will run only on the specified regions and not the whole genome sequence). SnoBIRD’s improvements also lie in the quality of output information it provides compared to the other tools. Indeed, SnoBIRD is the only approach that annotates all four boxes (especially the D and D’ motifs which are useful to locate the target base-pairing region on the snoRNA), that provides snoRNA pseudogene annotation as well as snoRNA features (box conservation, terminal stem and structural stability scores) and that organizes this information into an easy-to-read format such as TSV, FASTA, BED or GTF (**Table 1**). Such output formats can then be effortlessly integrated into subsequent analyses (e.g., finding the overlap of BED entries with other genes, computing the read abundance in RNA-Seq experiments, etc.). Finally, another output feature specific to SnoBIRD is that it limits the number of overlapping predictions through its merging step (**Supplementary Figure S2**). This is oftentimes not the case for the other tools (as depicted in **Supplementary Figures S18A-B**, **S19** and **S20A,D**), thus complexifying subsequent analyses by forcing the user to choose which overlapping prediction should be used as the potential “real” snoRNA gene.

Throughout this study, SnoBIRD has been applied to a wide range of different eukaryote species. Remarkably, SnoBIRD’s usage expanded the repertoires of both poorly characterized species and well annotated ones such as *S. pombe* and human, by either reannotating misannotated C/D box snoRNAs (**Supplementary Figures S19** and **S20**) or by identifying more than a hundred novel C/D box snoRNAs (**Supplementary Table S2**, **Figures 4B,E** and **5B**). We have also validated in *S. pombe* multiple C/D box snoRNA candidates (**Supplementary Figure S17** and we support their addition as expressed C/D box snoRNA genes in the next annotation release of PomBase (81). Interestingly, one of these validated SnoBIRD prediction was also recently characterized in parallel by another group (82). Indeed, the SnoBIRD prediction CD_531 (**Figure 4C-D**), identified as *snR107*, was shown to be hosted in a previously described longer noncoding RNA called *mamRNA* (83). This C/D box snoRNA was shown to exhibit two crucial functions: 1) to guide a site-specific 2’-O-methylation on the 25S rRNA to regulate ribosome production; 2) to interact with specific RBPs to regulate gametogenesis (82). The concurrent discovery of this same snoRNA by their approach and ours reinforces that SnoBIRD identifies relevant regulatory C/D box snoRNAs that are missing from current annotations. Additionally, further experiments are needed to confirm that the predictions in the other species are genuine snoRNAs (**Supplementary Table S2**), but based on the *S. pombe* results, we are confident that many of these high-confidence candidates are also bona fide C/D box snoRNAs.

In addition to displaying high performance and ease-of-use, SnoBIRD maintains a high degree of interpretability by integrating SHAP values as an intrinsic component of the pipeline (**Figure 3**). Interestingly, these SHAP values remain available as intermediate results for each prediction in the final output, allowing the user to dig deeper in the specifics of each SnoBIRD prediction (as highlighted in **Figure 4D**). This increased interpretability also represents one of few examples where large language models (LLMs) such as BERT can be interpreted and where they can be freed from their usual label as “black-box” models (84). Therefore, our approach could serve as a blueprint for other interpretable genomic tools (e.g., a H/ACA box snoRNA predictor) as it exploits the enhanced prediction performance of LLMs while providing a precise biological explanation for each prediction. Furthermore, SnoBIRD’s refinement step is similar to other approaches that try to distinguish expressed from pseudogene copies of different types of noncoding RNAs, e.g. tRNAs (85, 86). For instance, tRNAscan-SE, the most widely adopted tRNA predictor, includes a tRNA pseudogene detection module which classifies, based on secondary structure, if a given tRNA prediction is likely functional or a retrotransposed degenerate copy (86). These tRNA pseudogenes are also included in the affiliated GtRNAdb tRNA database. Thus, a similar strategy for snoRNA pseudogene prediction, such as the one implemented in SnoBIRD, is highly desirable as these snoRNA pseudogenes are different from the expressed snoRNAs in their expression level, sequence, structure and functionalities (**Supplementary Figure S5**) (24).

As SnoBIRD was purposefully trained to limit the number of false positives, it achieved by far the lowest number of total predictions amongst the different snoRNA-centric discovery tools (**Supplementary Figure S21**). Nonetheless, the total number of SnoBIRD predictions remains relatively high, with an increased proportion of snoRNA pseudogenes correlating with organism complexity (**Figure 5A**), raising the question as to why so many predictions. One hypothesis is that there might have been a high number of snoRNA duplication events that have occurred in ancestral genomes. Most of these duplicated snoRNAs would have integrated loci that are unfavorable to their expression (as hypothesized previously (24)), leading to their degeneration into the remnants identified by SnoBIRD as pseudogenes. Future studies could analyze in a systematic way (similar to our analysis of **Figure 5C-D**) the overall evolutionary trajectory of all SnoBIRD predictions across diverse eukaryote genomes, leading to better insights into the potential origin of these snoRNA predictions.

In terms of limitations, SnoBIRD only detects, by design, snoRNAs with a maximal length of 164 nt, which still corresponds to more than 95 % of annotated C/D box snoRNAs eukaryote-wide (**Supplementary Figures S1A** and **S6**). The excluded longer snoRNAs are usually *SNORD3* copies which also exist in shorter versions (and these shorter copies are predicted by SnoBIRD). Another drawback is that the performance of SnoBIRD’s refinement step is not as high as for the identification step. A potential explanation is that we did not include the host gene expression status as additional feature for the second BERT model (despite being reported as the fourth most important expression determinant of snoRNAs (24)). This design choice was driven by the fact that host gene expression levels cannot be inferred directly from the sequence, unlike the other used features (motif conservation, terminal stem and structural stabilities). By relying uniquely on the sequence as input, we simplified the input requirements, while still attaining reasonable performance on the pseudogene prediction task. We believe that a future iteration of SnoBIRD including as input such data (e.g., transcriptomic or even epigenetic datasets in addition to the sequence) could help improve this refinement step and provide more accurate snoRNA pseudogene predictions.

To conclude, SnoBIRD represents the most reliable tool to predict C/D box snoRNAs and classify them as expressed or pseudogenes across any eukaryote genome in a reasonable time. As snoRNAs are central regulators of several physiological and pathological processes (6–8), it is only through a better and more refined annotation that snoRNAs will be investigated thoroughly, leading to new discoveries regarding human and other eukaryotes molecular processes.

## DATA AVAILABILITY

The TGIRT-Seq datasets generated in this study (for *G. gallus*, *M. mulatta, M. musculus*, *S. cerevisiae* and *S. pombe*) were deposited on the Gene Expression Omnibus and are accessible under the id GSE290579. SnoBIRD tuning, training and test sets are available on Zenodo at https://zenodo.org/records/14927289. SnoBIRD training procedure as well as the analyses presented in this study are organized in a Snakemake workflow accessible at https://github.com/etiennefc/cd_predictor. Finally, SnoBIRD as a tool is freely accessible on GitHub at https://github.com/etiennefc/snoBIRD and on Zenodo at https://zenodo.org/records/14927205/files/snoBIRD.zip.

## SUPPLEMENTARY DATA

Supplementary Data are available at NAR online.

## AUTHOR CONTRIBUTIONS

Étienne Fafard-Couture: Conceptualization, Data Curation, Formal Analysis, Investigation, Methodology, Software, Validation, Writing—original draft, Writing—review & editing. Cédric Boulanger: Investigation, Resources, Validation. Laurence Faucher-Giguère: Resources. Vanessa Sinagoga: Resources. Mélodie Berthoumieux: Resources. Jordan Hedjam: Resources. Virginie Marcel: Supervision. Sébastien Durand: Supervision. Mark A. Bayfield: Supervision. François Bachand: Supervision. Sherif Abou Elela: Supervision. Pierre-Étienne Jacques: Conceptualization, Supervision, Writing—review & editing. Michelle S. Scott: Conceptualization, Funding Acquisition, Supervision, Writing—review & editing.

## ACKNOWLEDGMENTS

We would like to thank Benjamin Gibert and Maëva Hervieu for carrying out mice maintenance and obtaining ethical approval. We would also like to acknowledge Mathieu Catala’s help with sample and antibody selection as well as manuscript revision. We also extend our gratitude to the DRAC and Calcul Québec for providing support and access to advanced research computing resources. Finally, we would like to thank members of the different labs for their thoughtful insights and help in optimizing SnoBIRD for its public release.

