## Supplementary Figures for "SnoBIRD: A tool to identify C/D box snoRNAs and refine their annotation across all eukaryotes"

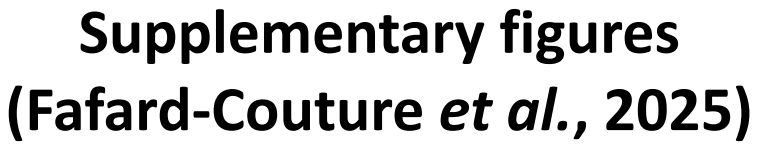

**Supplementary figures**
**(Fafard-Couture *et al.*, 2025)**

**A**

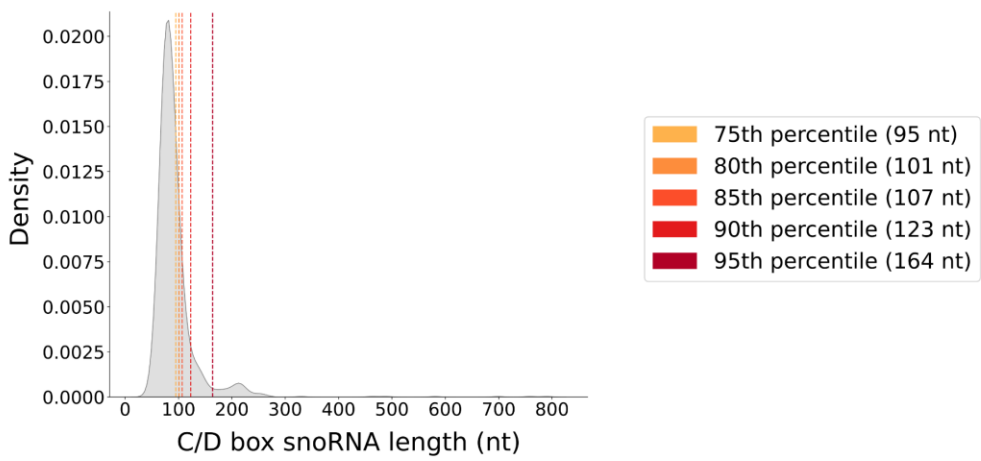

**B**

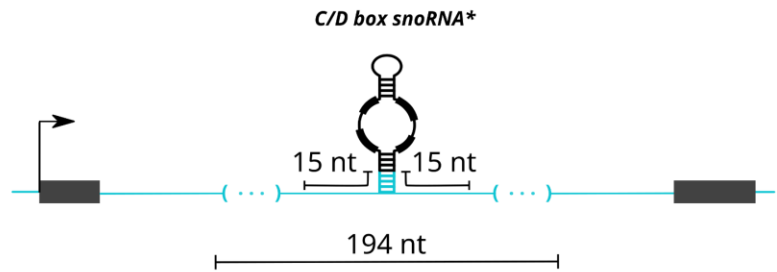

**C**

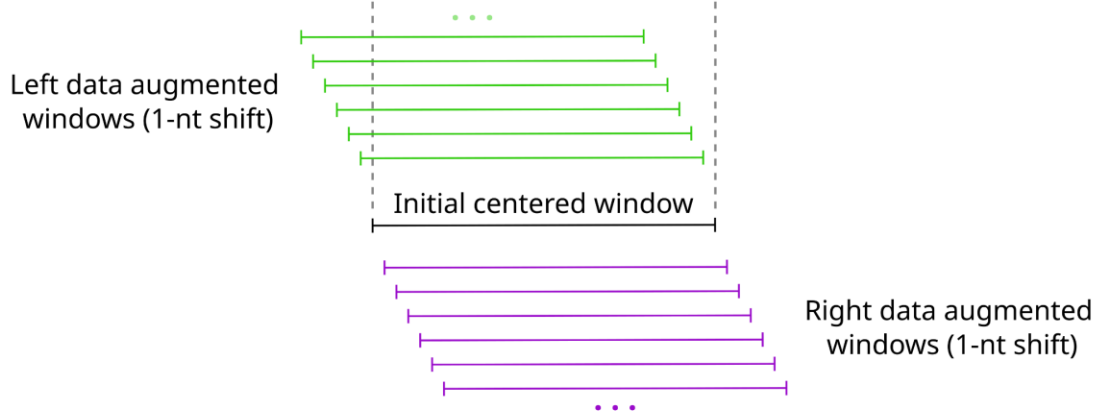

**Figure S1 (Supplemental to Methods and Figures 1 and 2). SnoBIRD's training window selection and data augmentation.** **(A)** Length distribution of the collected C/D box snoRNAs (no filter applied). Vertical dotted lines represent the length thresholds from the 75<sup>th</sup> to the 95<sup>th</sup> percentile. The 164 nt threshold was used to keep all C/D box snoRNAs below or equal to that threshold (i.e. 1184 expressed C/D box snoRNAs and 432 C/D box snoRNA pseudogenes). **(B)** The choice of SnoBIRD's input window size length was fixed at 194 nt, which comprises the centered C/D box snoRNA with its flanking 15 nucleotides. If the snoRNA is smaller than 164 nt, the window is further extended within surrounding nucleotides (represented by the blue dots in parentheses). The \* represents the fact that this window extension strategy was also applied to all other midsize noncoding RNAs smaller than 164 nt and present in the initial dataset as negative examples. **(C)** Data augmentation was applied by shifting the initial centered window both left and right by 1 nt increments to create new synthetic examples of the C/D box snoRNAs examples only. The dataset used for SnoBIRD's first model tuning/training/test was data augmented by a factor of 10 (5 shifts on both sides), whereas the dataset for SnoBIRD's second model was data augmented by a factor of 10 for expressed C/D box snoRNAs and by 30 (15 shifts on both sides) for C/D box snoRNA pseudogenes to create an equal number of examples of both classes.

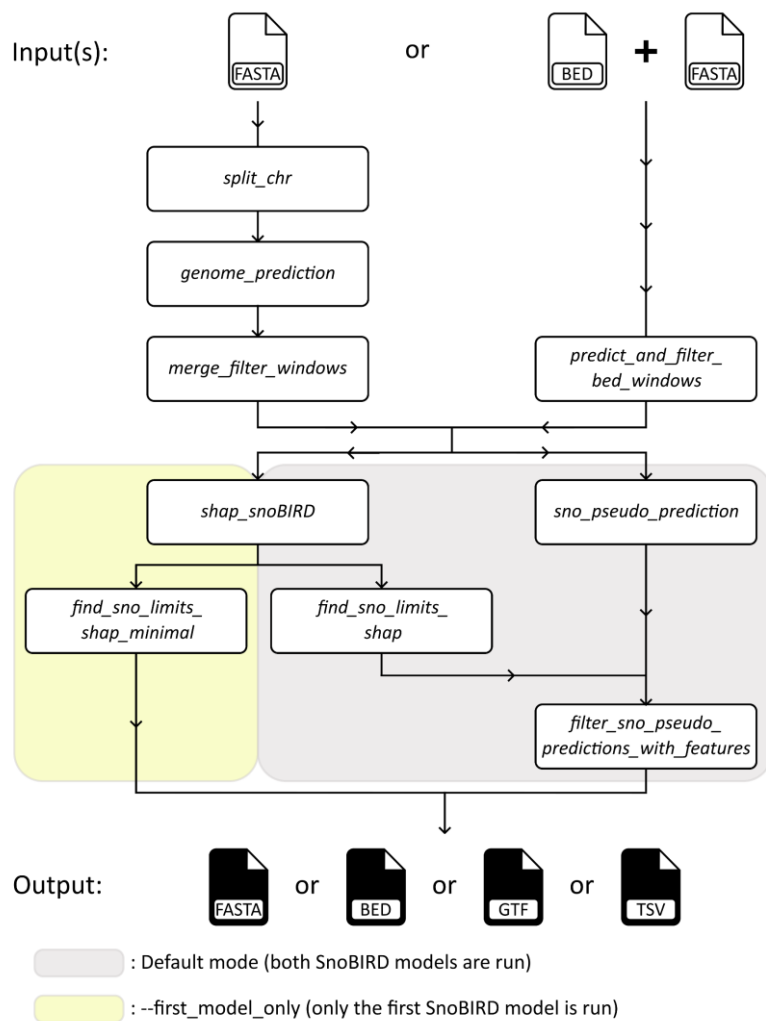

**Figure S2 (Supplemental to Methods and Figure 2). Overview of the main steps and options of SnoBIRD.**

The input given to SnoBIRD can either be a FASTA file (contain one or multiple sequences) or a BED file of regions of interest as well as a FASTA file of the input genome sequence (separated per chromosome). When using only a FASTA file as input, this input is split in different chromosome and/or smaller chunks of chromosome files (via *split\_chr*) to parallelize downstream jobs. These split sequences are fed to SnoBIRD for prediction of the presence of C/D box snoRNAs in the general sense (via *genome\_prediction*). The positive windows are next merged and filtered based on the *p1* (probability of the first model) and *w* (number of consecutive positive windows) parameters to return centered positive windows (via *merge\_filter\_windows*). If a BED input file is used, only the windows present in this input are predicted on by the first model of SnoBIRD and later filtered using the *p1* parameter to return centered positive windows (via *predict\_and\_filter\_bed\_windows*). Using the default parameters, SnoBIRD will next run the second model which will output a probability of being either an expressed C/D box snoRNA or a pseudogene (via *sno\_pseudo\_prediction*). In parallel, SHAP values will be computed for each centered positive window (output of the merging step after the first model) (via *shap\_snoBIRD*) and these SHAP values will be used to identify the C and D boxes as well as to define the start and end of the snoRNA within the 194 nt window (via *find\_sno\_limits\_shap*). Finally, the predictions of the second model combined with feature filters (based on the terminal stem score and structural score, as well as on mutations found in the different boxes) will be filtered and generated as the final output (via *filter\_sno\_pseudo\_predictions\_with\_features*). The final output is user-defined within the following choices: FASTA, BED, (gene transfer format) GTF or (tab-separated value) TSV file. Of note, if the *--first\_model\_only* option is used, the second model predictions will be skipped and only the predictions of the first model with found boxes based on the computed SHAP values will be produced as output (via *find\_sno\_limits\_shap\_minimal*).

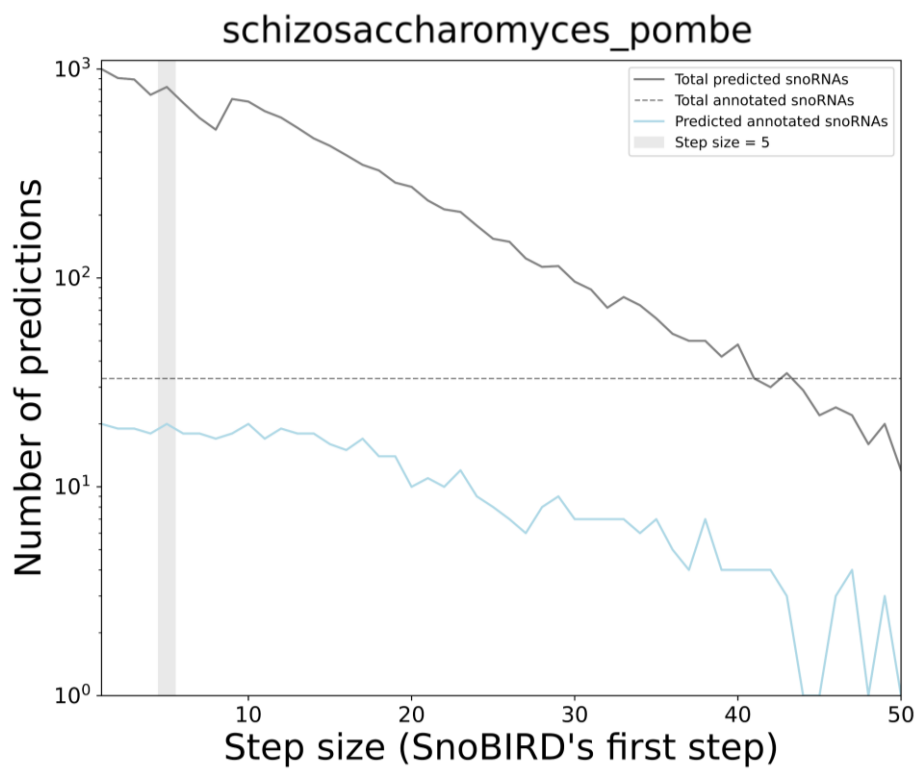

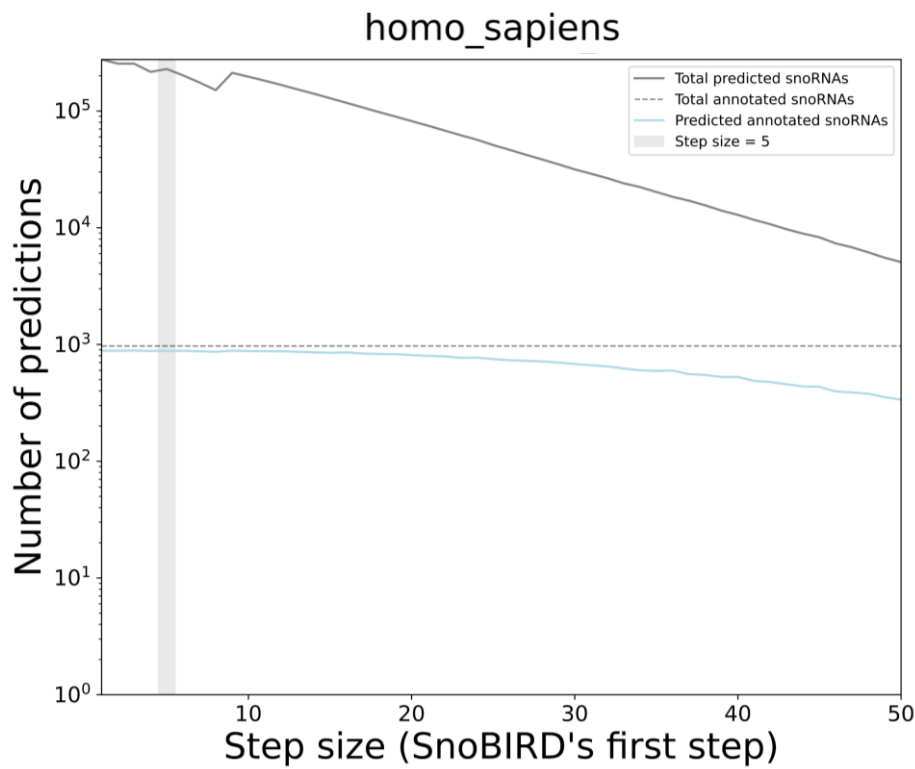

**Figure S3 (Supplemental to Methods). Effect of step size on SnoBIRD's prediction in *S. pombe* and *H. sapiens*.** Line plots depicting for *S. pombe* (top panel) and *H. sapiens* (bottom panel) the total number of SnoBIRD predictions (black line) and those overlapping annotated C/D box snoRNAs (blue line) as a function of the step size used when running SnoBIRD. The dotted horizontal line represents the number of C/D box snoRNAs annotated in that species. The shaded grey vertical region represents the chosen default step size value (step size=5) which is conservative but maximizes the number of predicted annotated C/D box snoRNAs while limiting the total number of predictions.

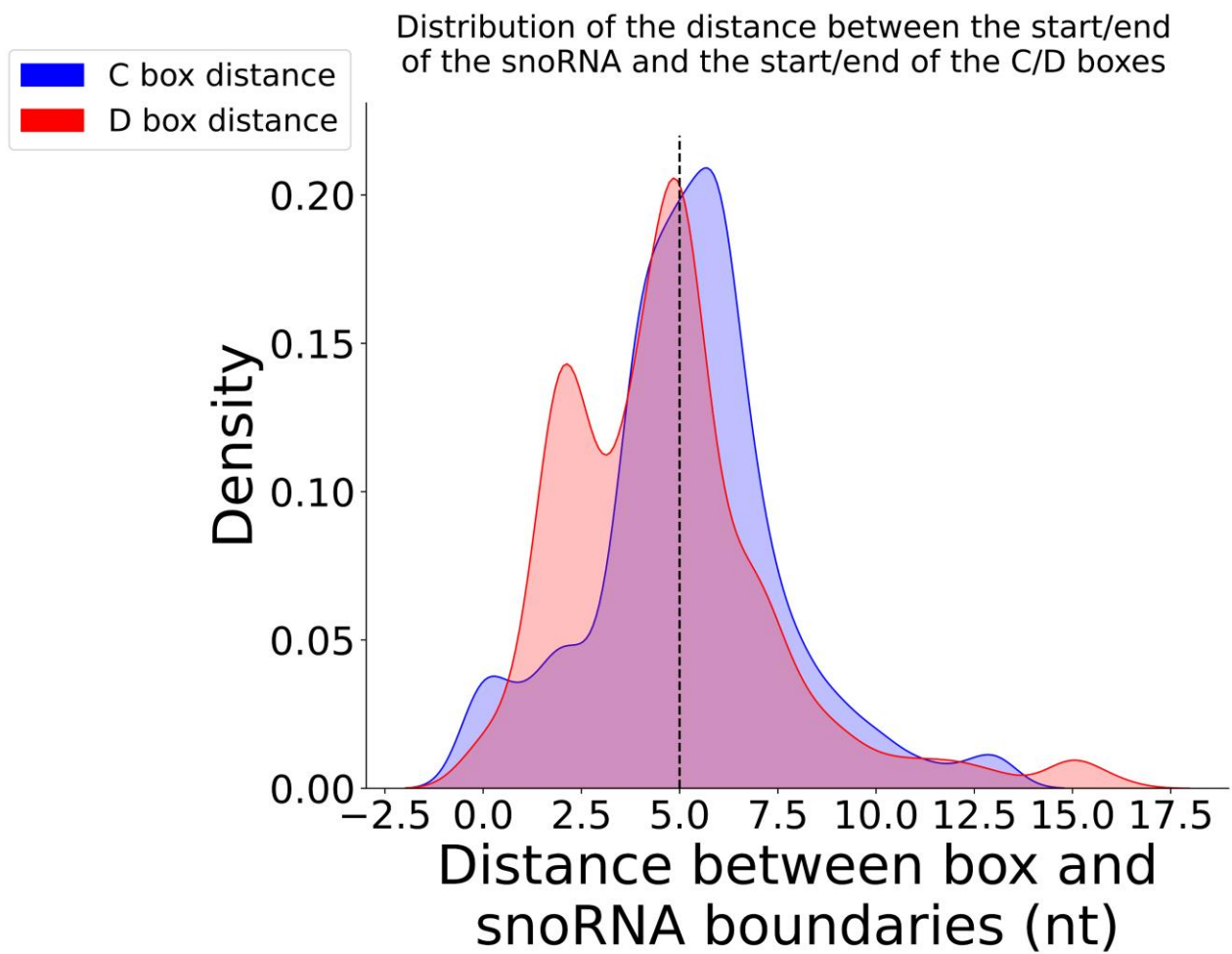

**Figure S4 (Supplemental to Methods and Figure 3). Comparison of the distance between annotated start/end and box positions of known C/D box snoRNAs.** Density plots showing the distribution of the distance between the annotated start of C/D box snoRNAs and the start of their C box (blue curve) or the distance between the annotated end of C/D box snoRNAs and the end of their D box (red curve). The vertical dotted line represents the threshold of 5 that was chosen, i.e. that snoRNAs predicted by SnoBIRD all have their start defined at 5 nt upstream of their C box and their end defined at 5 nt downstream of their D box.

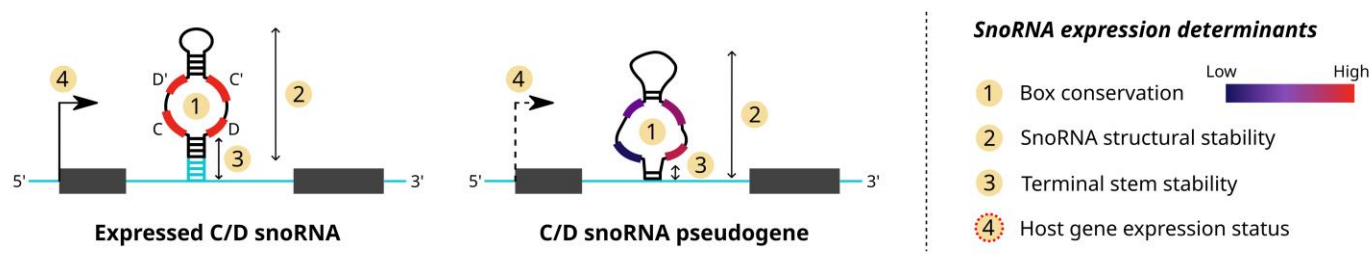

**Figure S5 (Supplemental to Figure 1). Expressed C/D box snoRNAs and C/D box snoRNA pseudogenes display different expression determinants.** Schematic representation of expressed C/D box snoRNAs and snoRNA pseudogenes. SnoRNAs should be divided in these two subclasses since they greatly differ in terms of their cellular function as well as their expression determinants. Expressed snoRNAs display conserved boxes, stable secondary structure and terminal stem and are usually located in expressed host genes. Conversely, snoRNA pseudogenes accumulate several mutations in their boxes, show unstable secondary structure and terminal stem and are generally located in non-expressed loci. Interestingly, except for the host gene expression status (red dotted circle), the other distinguishing features are all related directly to the snoRNA sequence.

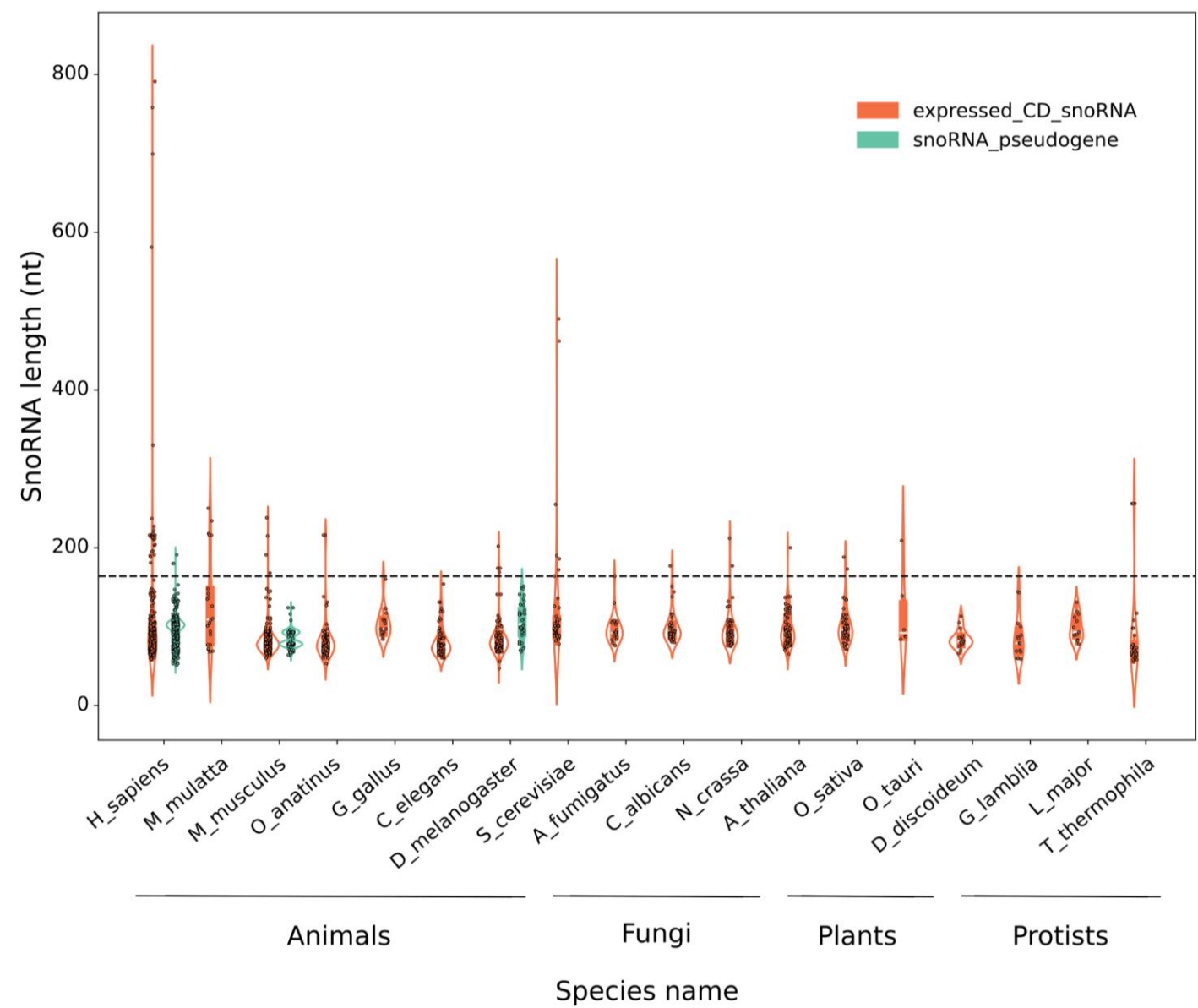

**Figure S6 (Supplemental to Figures 1 and 2). Length distribution of annotated C/D box snoRNAs across species and snoRNA subclasses.** Violin plots showing the length distribution of annotated expressed C/D box snoRNAs and snoRNA pseudogenes present in the initial dataset. C/D box snoRNA pseudogenes are present only for species with enough TGIRT-Seq datasets to reliably assess their non-expression. The horizontal dotted line represents the chosen maximal length threshold of 164 nt.

**Species distribution across**
**C/D snoRNA pseudogenes**

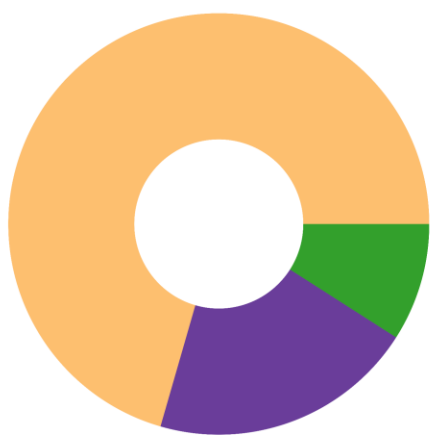

**Species distribution across**
**negative examples**

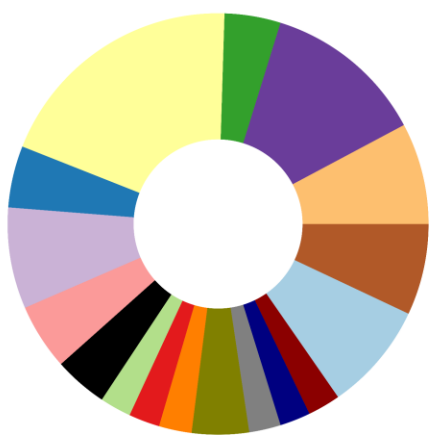

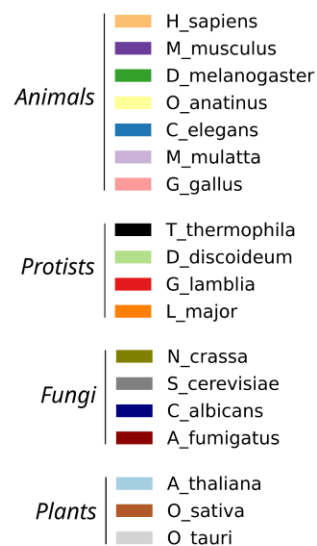

**Figure S7 (Supplemental to Figure 1). Species distribution across C/D box snoRNA pseudogenes and negative examples.** Donut charts representing species distribution from the initial dataset of the C/D box snoRNA pseudogenes class (left), i.e. for species with enough TGIRT-Seq datasets to reliably assess global snoRNA expression level, and of the negative examples (right), i.e. midsize noncoding RNAs, random exonic, intronic and intergenic regions and shuffled C/D snoRNA sequences.

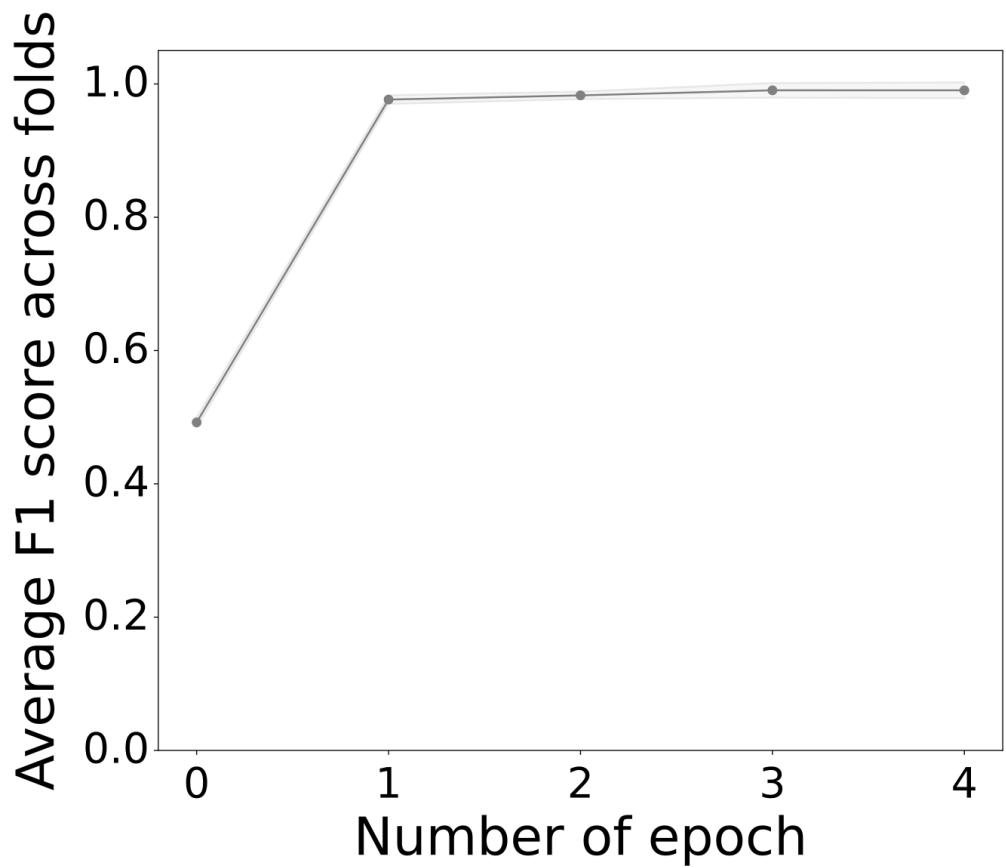

**Figure S8 (Supplemental to Figure 2). Training learning curve of SnoBIRD's first model.** Line plot depicting the average F1-score across the 10 folds during the 4 epochs of training of SnoBIRD's first model. The shaded grey cloud over and under the line represents the F1-score standard deviation across folds.

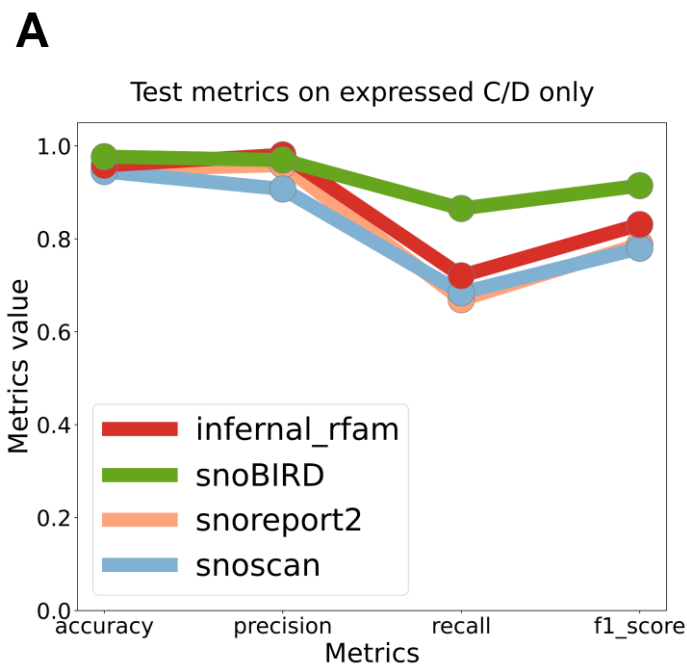

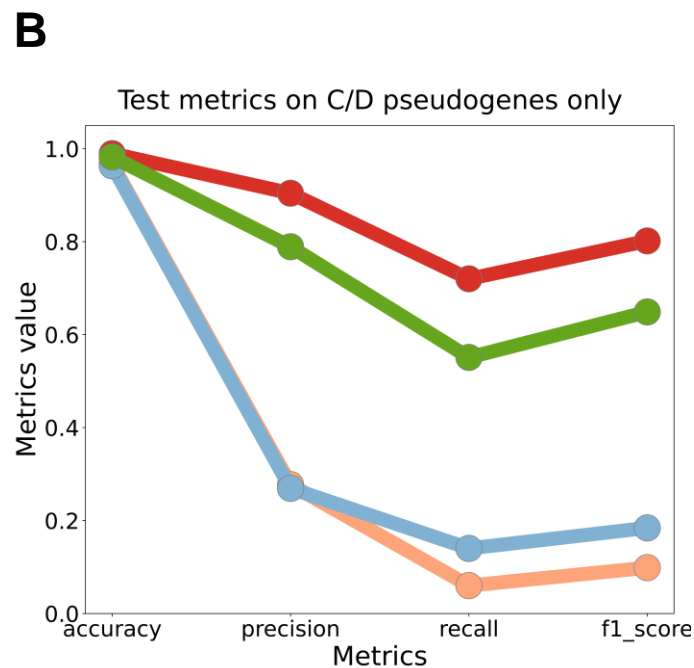

**C** Identification step true negative and false positive examples (in the test set) across tools

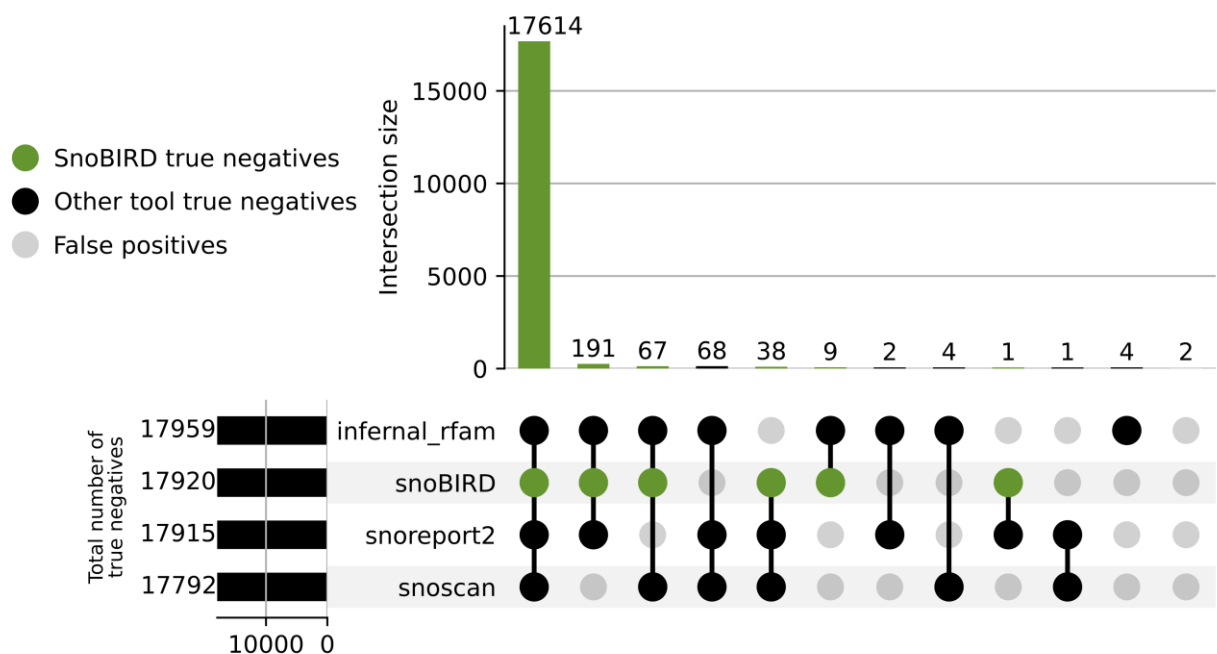

**Figure S9 (Supplemental to Figure 2). SnoBIRD outperforms most tools in terms of per subclass predictions and true negative detection.** (A) Test set metrics based on the expressed C/D box snoRNA and negative classes only. (B) Test set metrics based on the C/D box snoRNA pseudogene and negative classes only. (C) Upset plot representing the intersection of predictions between the tools regarding the negative class (i.e. not C/D box snoRNAs) predictions in the test set. The green dots and vertical bars represent SnoBIRD's true negatives (a negative predicted as such, i.e. not as a C/D box snoRNA), the black dots and vertical bars represent the other tools' true negatives, whereas the gray dots and vertical bars represent false positives (negative examples that were predicted as C/D snoRNAs). The black horizontal bars represent the total number of true negative predictions per tool.

Number of wrongly predicted examples in the test set

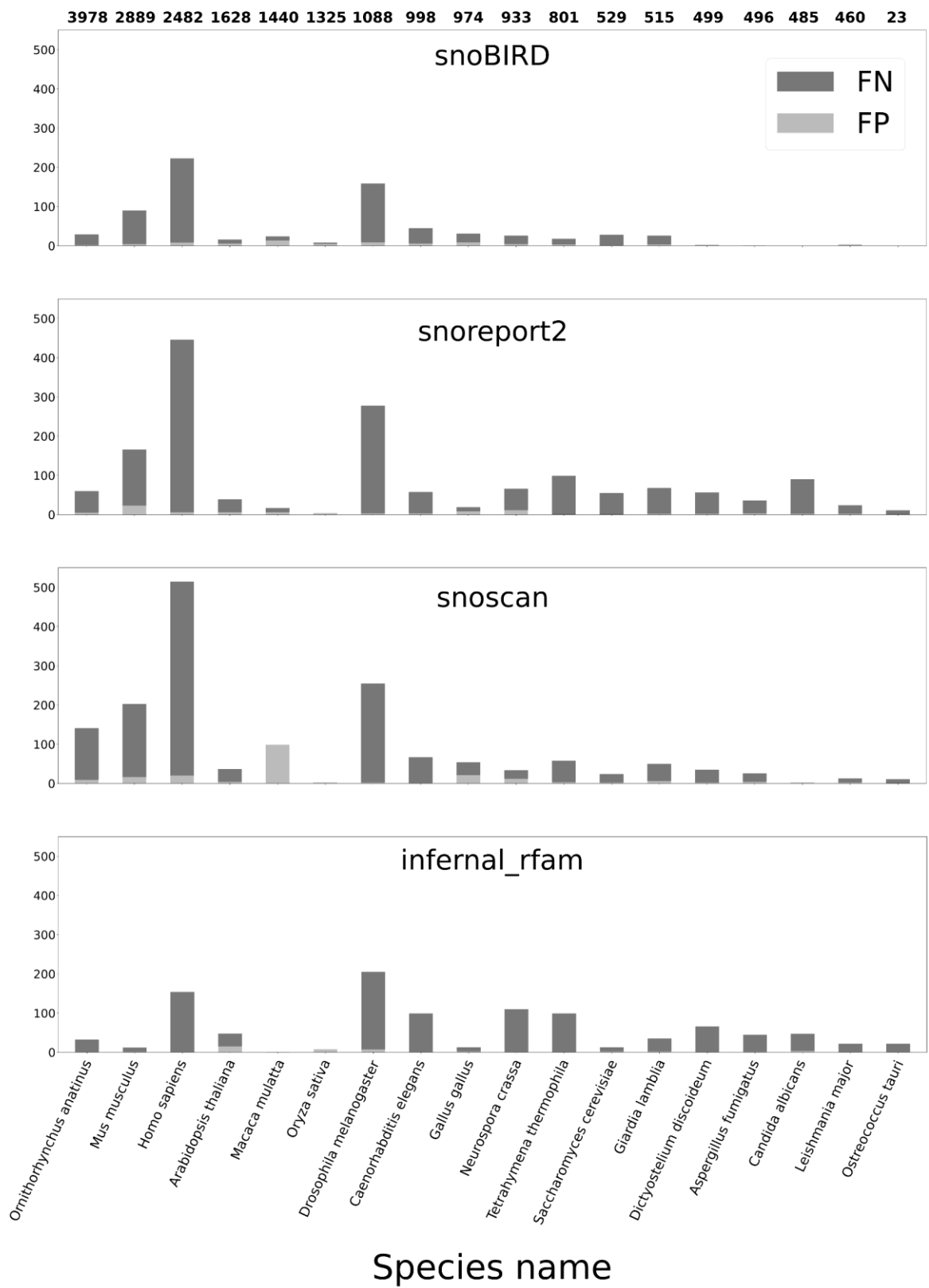

**Figure S10 (Supplemental to Figure 2). SnoBIRD shows less false negatives and across a greater variety of eukaryotes than the other tools.** Stacked bar plots showing the number of false negative (FN) and false positive (FP) examples from the test set across the different tools and per species. The total number of positive data-augmented examples per species present in the test set is represented as bold numbers above the topmost bar plot. Bars are ordered from the most to the least number of examples per species in the test set (from left to right respectively).

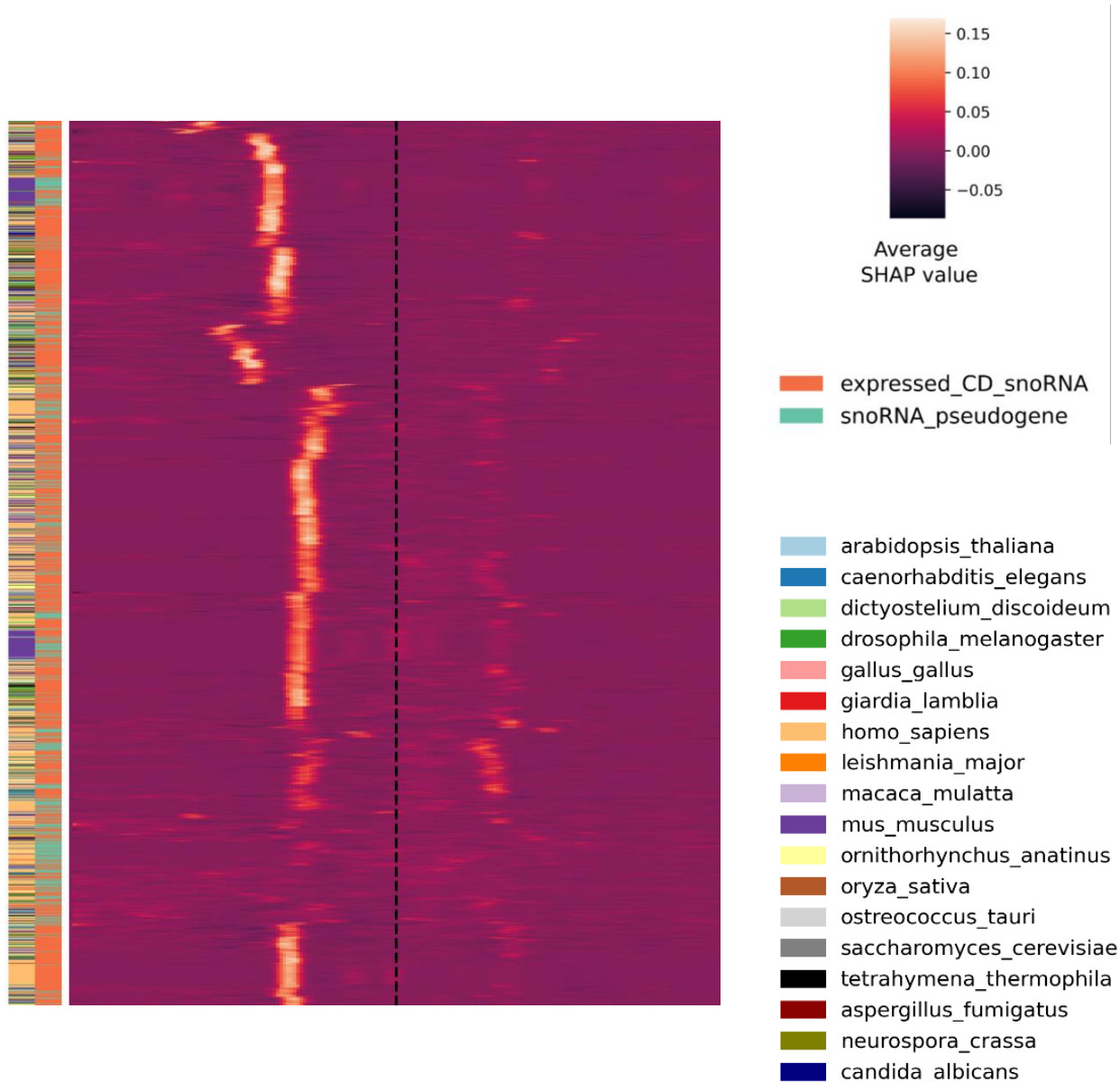

**Figure S11 (Supplemental to Figure 3). SnoBIRD’s prediction rely on different location and combination of C and D boxes.** Heatmap showing the importance of each nucleotide (nt) in the input sequence (194 nt window) for the identification step of SnoBIRD (prediction of C/D box snoRNA in the general sense). Stacked horizontal lines represent all the C/D snoRNAs in the initial dataset that were predicted as such by SnoBIRD and are clustered by SHAP profiles resemblance. Vertical lines in the heatmap represent the first to last nucleotide (from left to right respectively) in the window encompassing the snoRNA. The dotted vertical line marks the center of the window. The nucleotide importance is color-shaded and based on its average SHAP value across all 6-mers containing that given nucleotide (the legend is on the right). Species from which the snoRNAs originate are identified by the left most color bar. Expressed C/D box snoRNAs and snoRNA pseudogenes can be differentiated by the second color bar on the left.

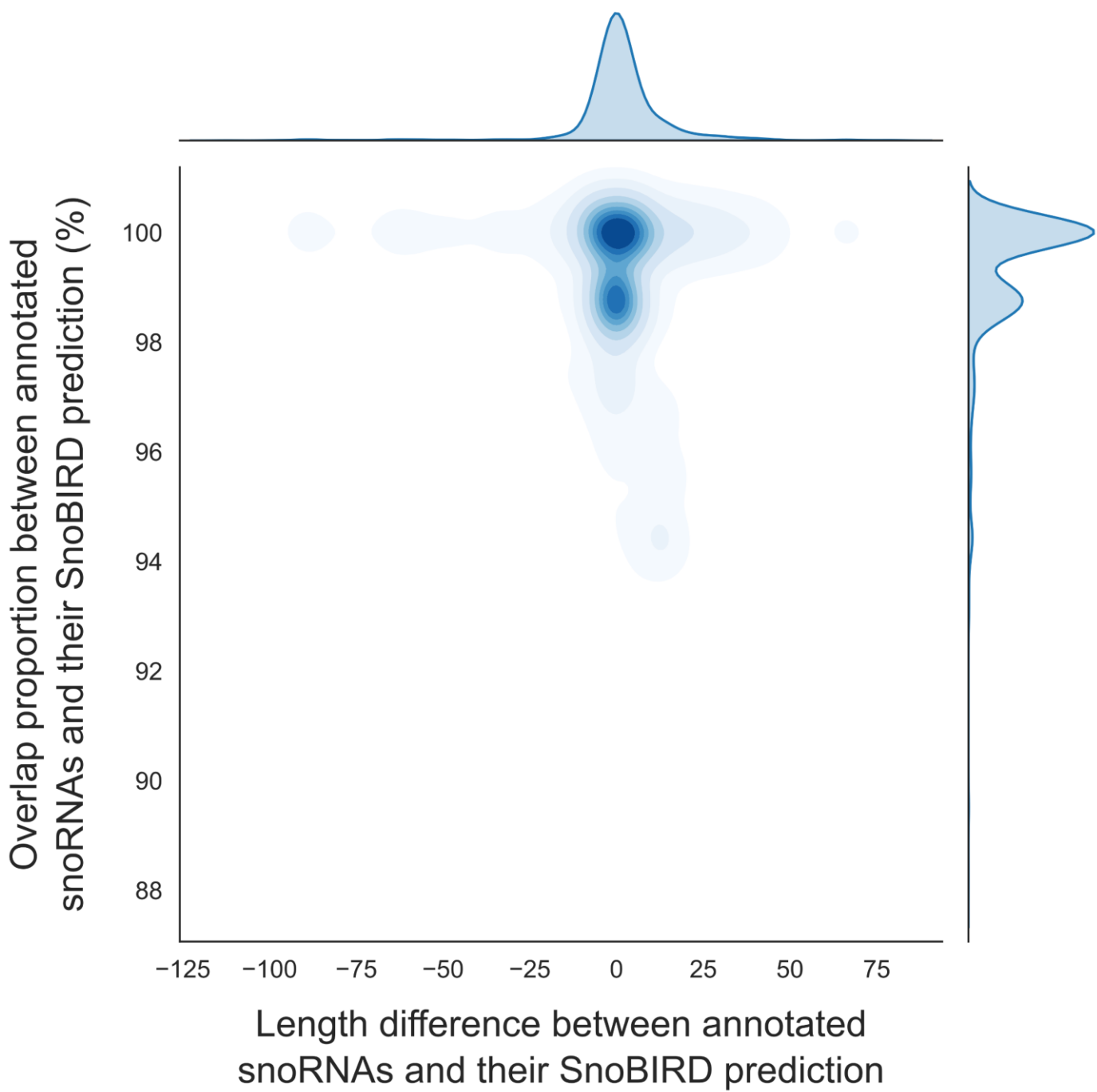

**Figure S12 (Supplemental to Figure 3). SnoBIRD predicted snoRNA limits are concordant with the annotated snoRNA limits.** Joint density plot depicting for the C/D box snoRNAs present in the initial dataset the length difference between SnoBIRD predictions and their known annotated length (x axis), as well as the overlap proportion between SnoBIRD predictions and their known annotated sequence (y axis).

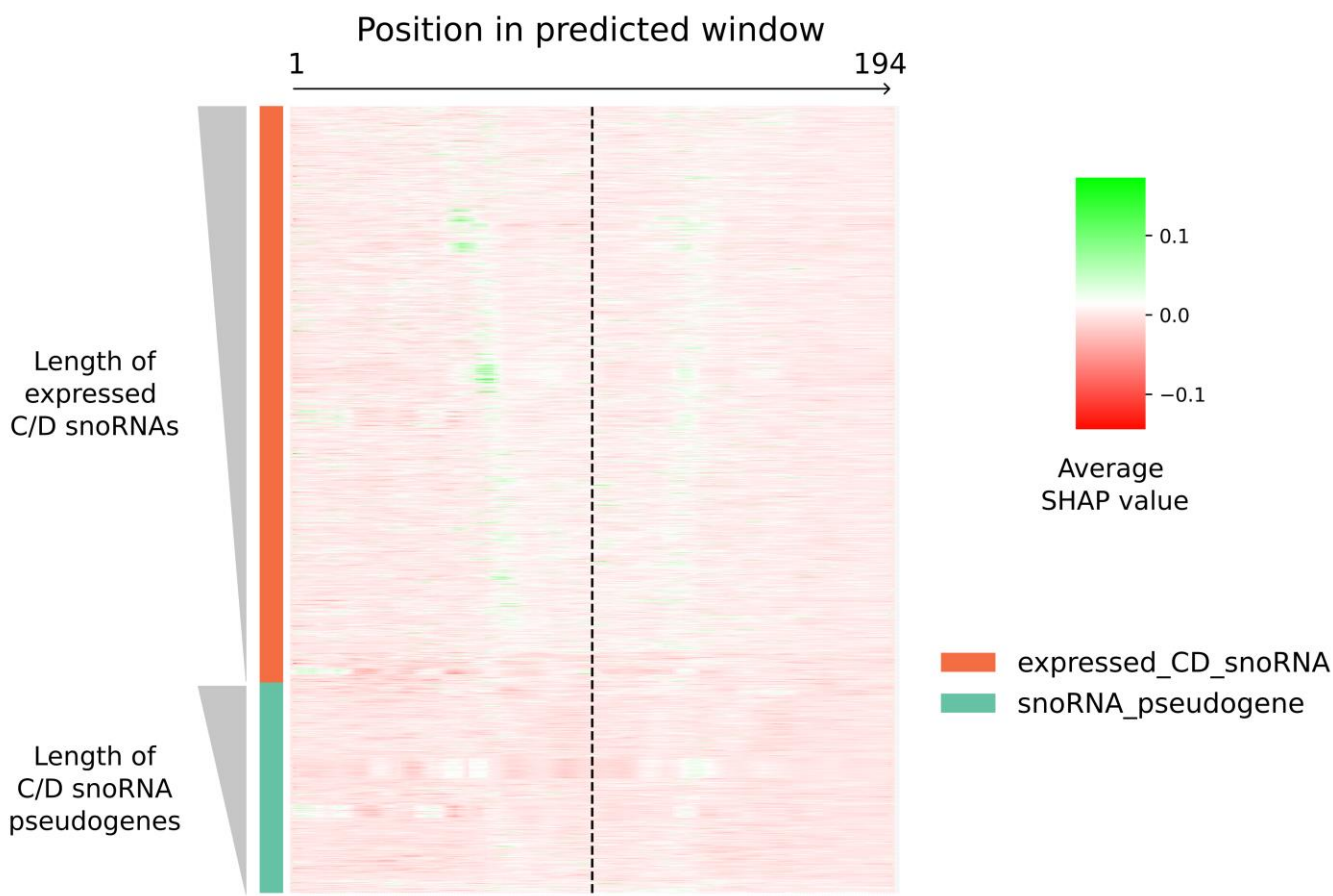

**Figure S13 (Supplemental to Figure 3). SnoBIRD's second model identifies some low-intensity biological signal.** Heatmap showing the importance of each nucleotide (nt) in the input sequence (194 nt window) for the refinement step of SnoBIRD (prediction of expressed C/D box snoRNA vs pseudogene). Stacked horizontal lines represent all the C/D box snoRNAs in the initial dataset that were predicted as such by SnoBIRD. Expressed C/D box snoRNAs were separated from snoRNA pseudogenes (represented by the left color bar), and both classes were sorted by increasing length of snoRNA within their fixed 194 nt window length (represented by the left grey gradient triangles). Vertical lines in the heatmap represent the first to last nucleotide (from left to right respectively) in the window encompassing the snoRNA. The dotted vertical line marks the center of the window. The nucleotide importance is color-shaded and based on its average SHAP value across all 6-mers containing that given nucleotide.

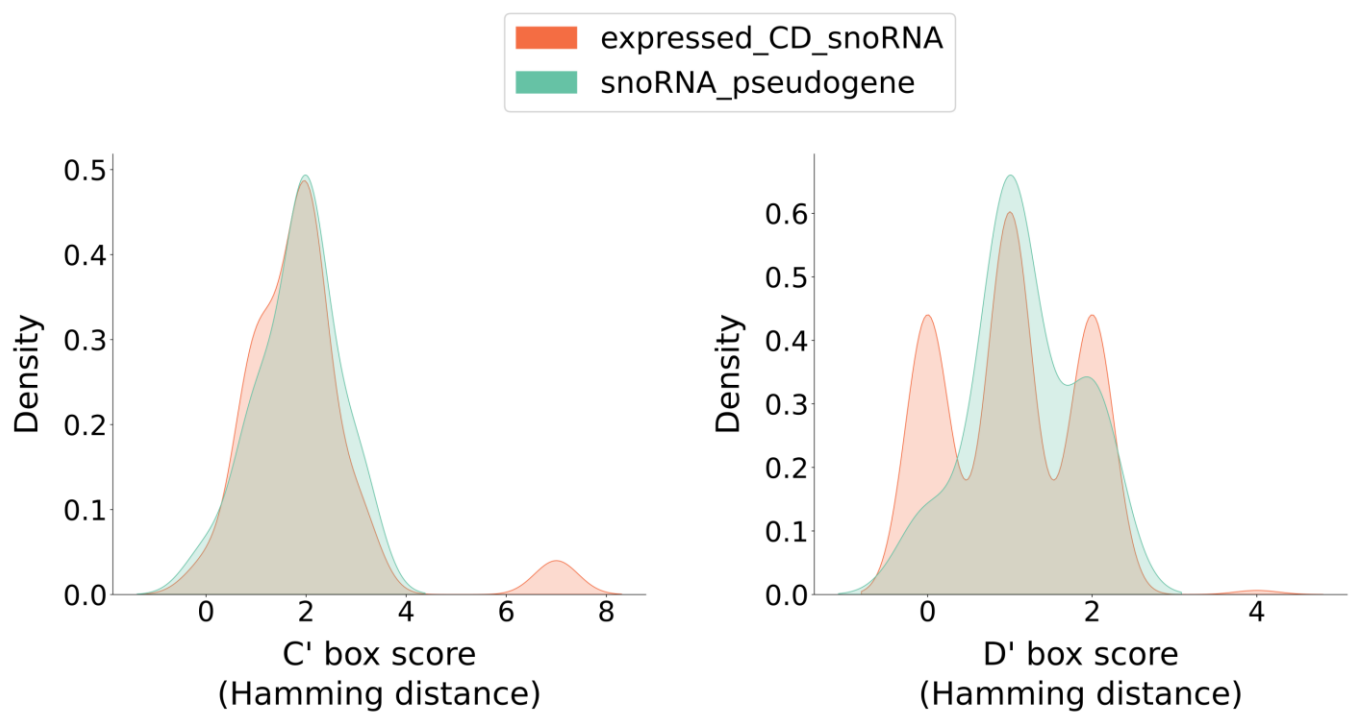

**Figure S14 (Supplemental to Figure 3). C' and D' box score distributions.** Density plots highlighting differences in snoRNA features between expressed C/D snoRNAs (orange) and snoRNA pseudogenes (teal) in the test set based on the boxes that SnoBIRD found in the predicted snoRNAs (left: C' box score; right: D' box score). Each score corresponds to the number of mutations (i.e. Hamming distance) with regards to the respective consensus sequence.

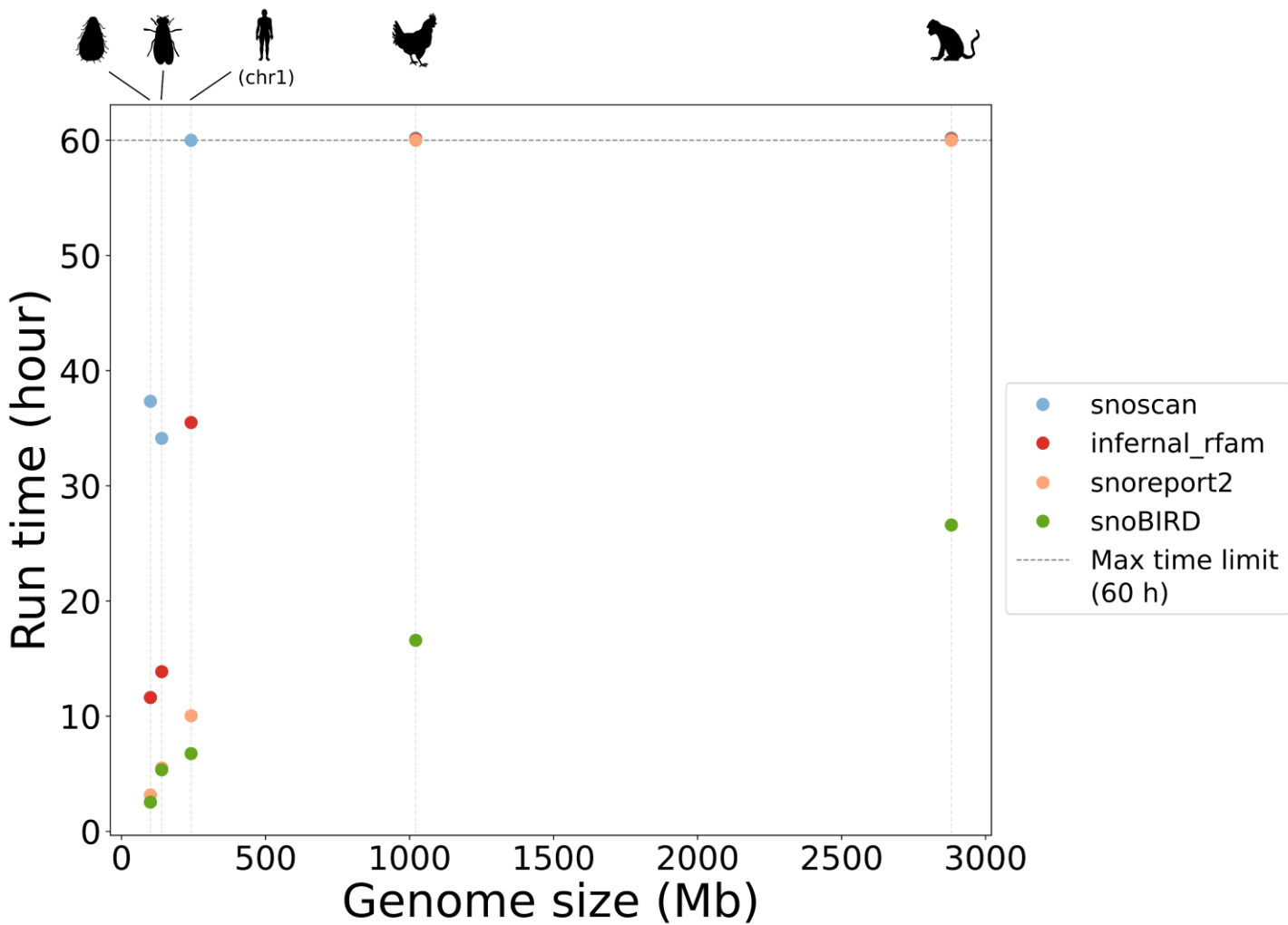

**Figure S15 (Supplemental to Figure 4). SnoBIRD is the only tool to scale well with increasing genome size.** Scatter plot showing the prediction overall run time per tool on genomes or sequences of varying length, namely *T. thermophila*, *D. melanogaster*, *H. sapiens'* chr1, *G. gallus* and *M. mulatta* genomes (each genome/sequence is marked by a light vertical dotted line). Prediction time was capped at maximally 60 hours (horizontal dotted line) to limit carbon emission as well as needless energy consumption, as only SnoBIRD clearly shows a reasonable overall run time. Of note, the human chr1 was included as control to show the performance on a single chromosome (compared to multi-chromosome sequences for the other species).

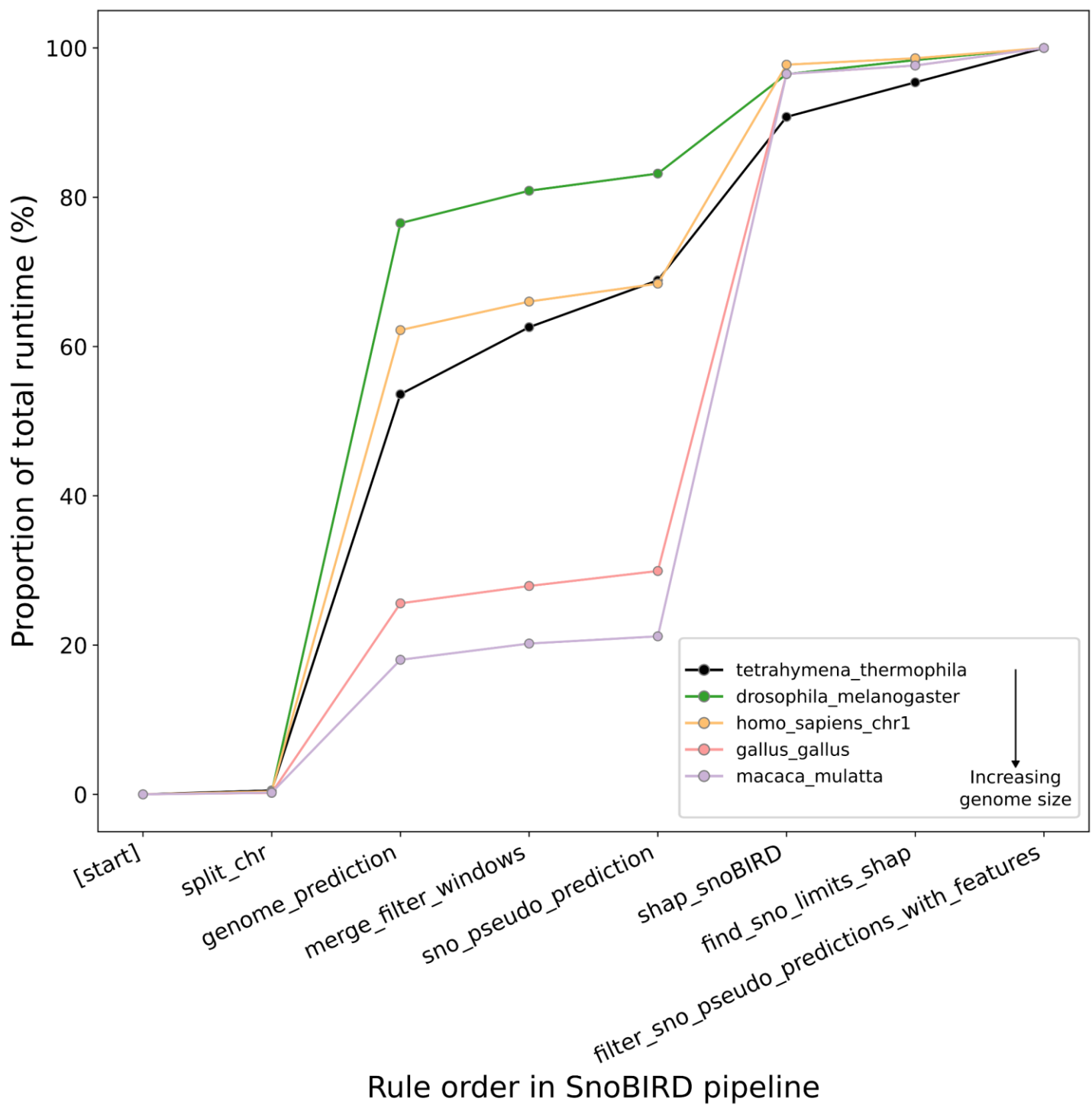

**Figure S16 (Supplemental to Figure 4). Run time per step in SnoBIRD's pipeline varies depending on input genome size.** Cumulative line plots highlighting the cumulative time (in percentage) for each step along SnoBIRD pipeline (steps are ordered on the x axis by their time of completion from the start to the end of the SnoBIRD's call) for different inputs of varying size. All SnoBIRD runs were with the same default parameters with the following updated options: -cs 10 -G V100 .

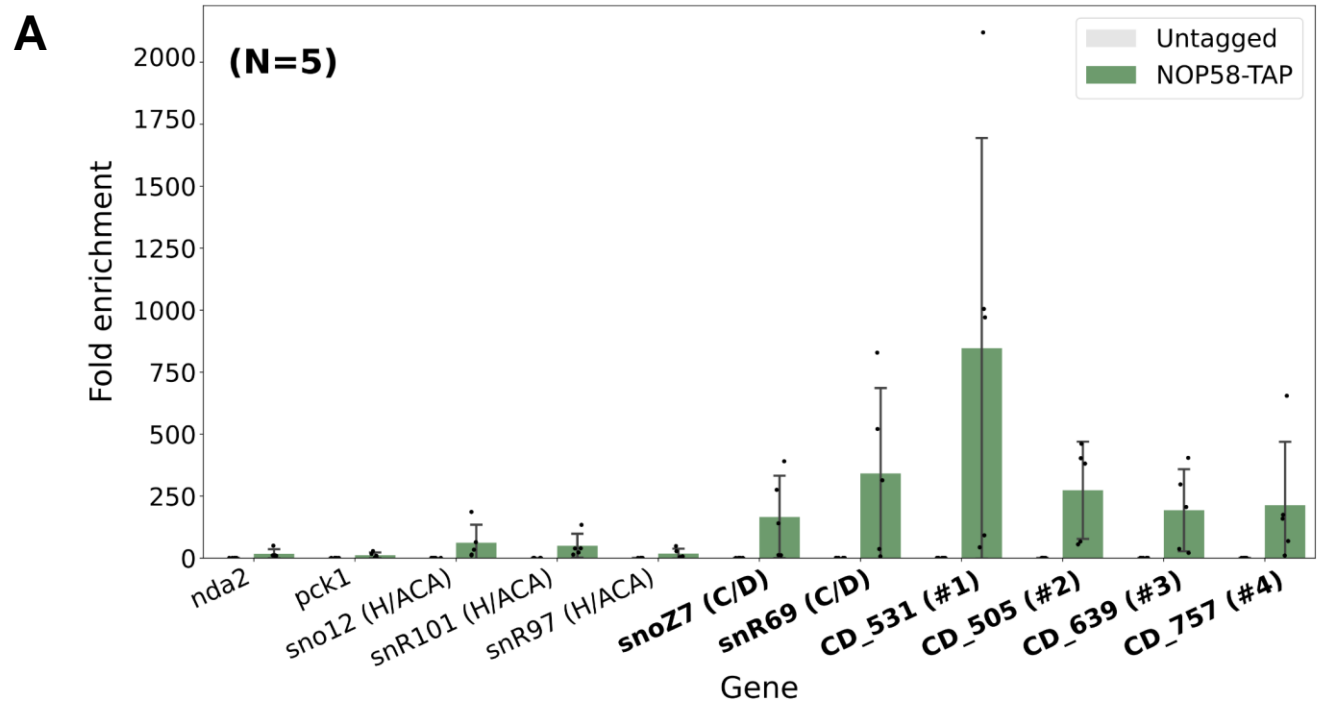

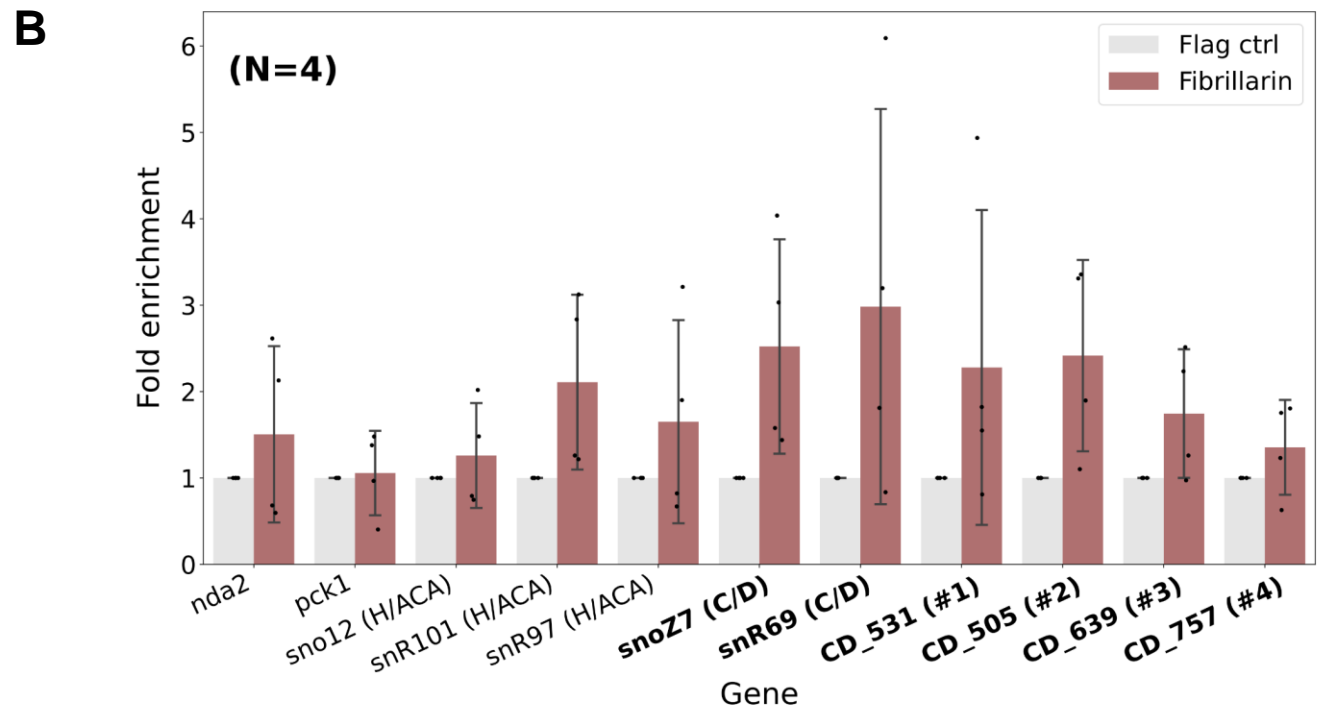

**Figure S17 (Supplemental to Figure 4). SnoBIRD predictions in *S. pombe* are experimentally validated by their immunoprecipitation with core proteins of the C/D box snoRNP. (A) Bar plot showing the fold enrichment between input and immunoprecipitation (IP) of NOP58-TAP-tagged samples of different RNAs in *S. pombe* (n=5). The right bars show the fold enrichment in the NOP58-TAP-tagged *S. pombe* strain whereas the left bars show the results in the control untagged strain. The expression level was assessed using quantitative PCR (qPCR). Negative gene controls include the protein-coding genes *nda2* and *pck1* as well as the H/ACA box snoRNAs *sno12*, *snR101* and *snR97*. Positive controls include the known C/D box snoRNAs *snoZ7* and *snR69*. Four different SnoBIRD predictions were tested (#1 to #4). (B) Same as (A), but the IP was done using a fibrillarin antibody (right bars) or a control flag antibody (left bars) (n=4). The positive controls' (C/D box snoRNAs) and the SnoBIRD-predicted C/D box snoRNAs' names are emphasized in bold.**

A

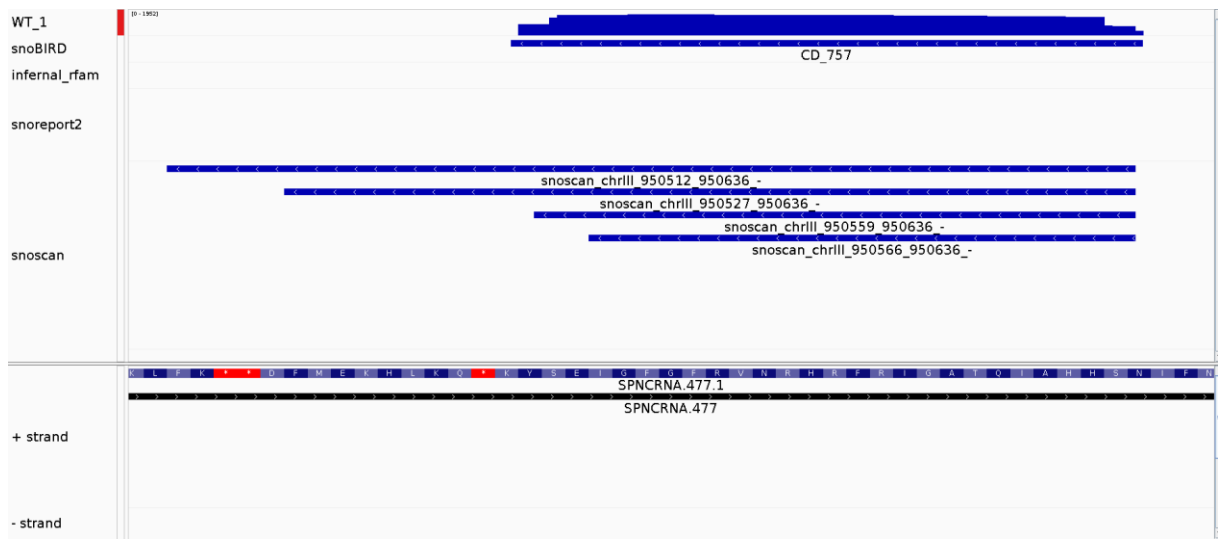

B

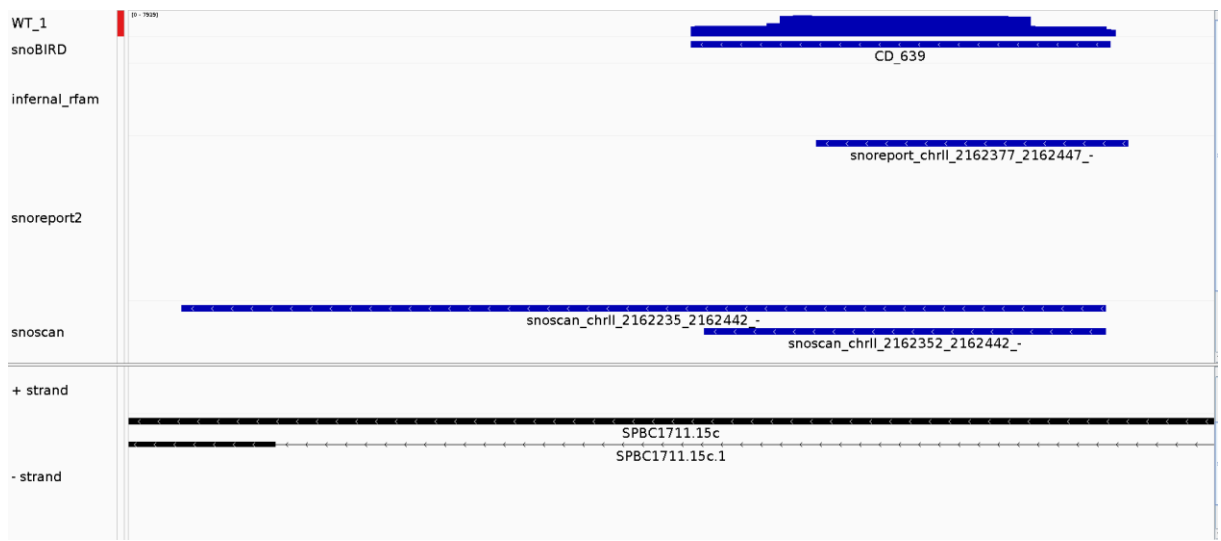

C

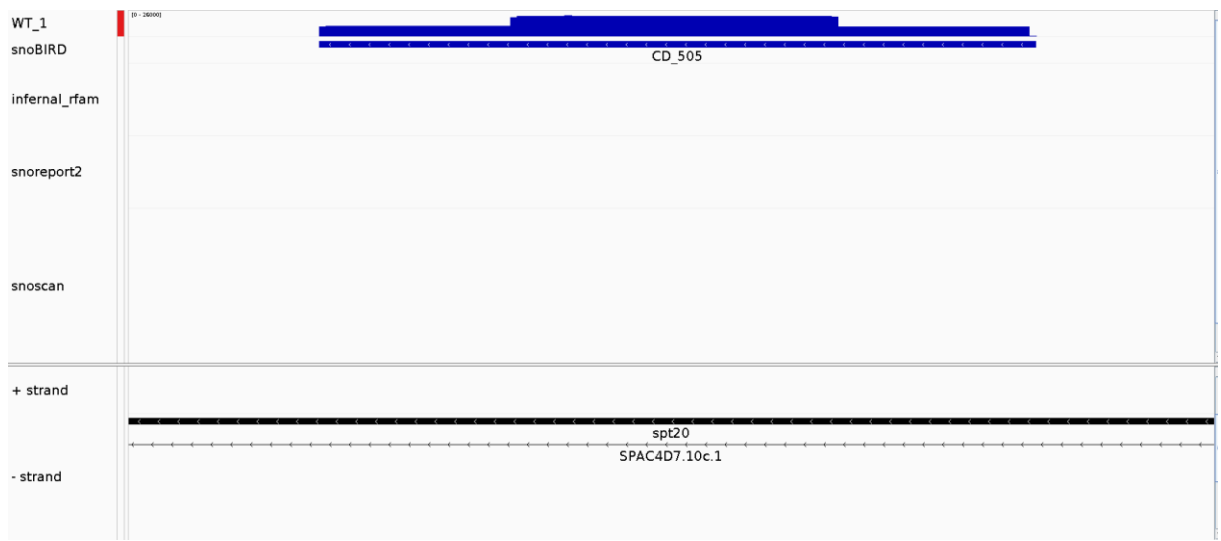

**Figure S18 (Supplemental to Figure 4). SnoBIRD's prediction in *S. pombe* identifies novel C/D box snoRNA candidates.** Integrative Genome Viewer (IGV) screenshots showing SnoBIRD'S predictions and C/D box snoRNA candidates CD\_757, CD\_639 and CD\_505, which were all validated to bind to core proteins of the C/D box snoRNP (Supplementary Figure S17). All images show as different tracks (from top to bottom) read accumulation in a wild type (WT) strain, predictions by the different tools and current genomic annotation.

A

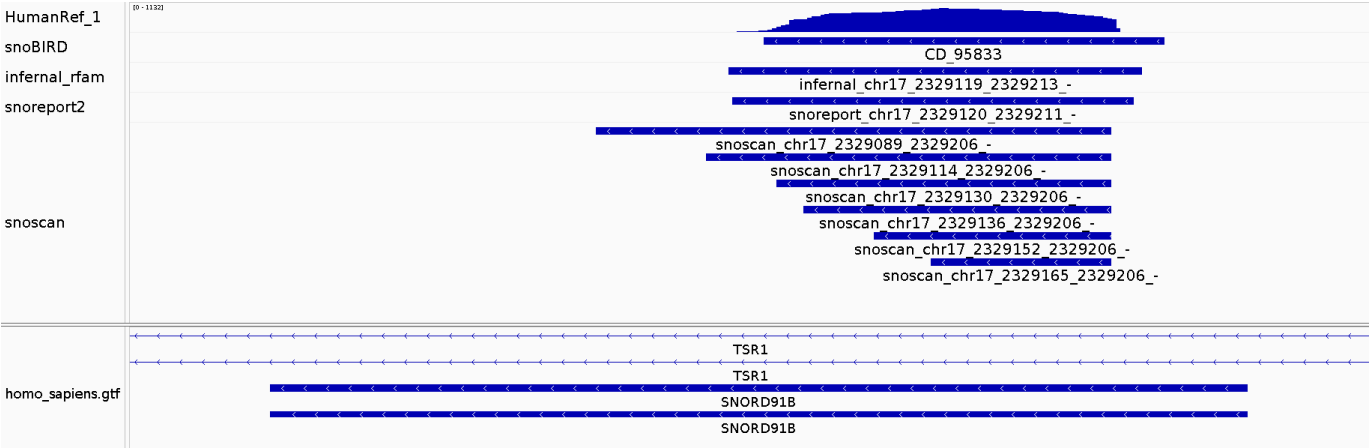

B

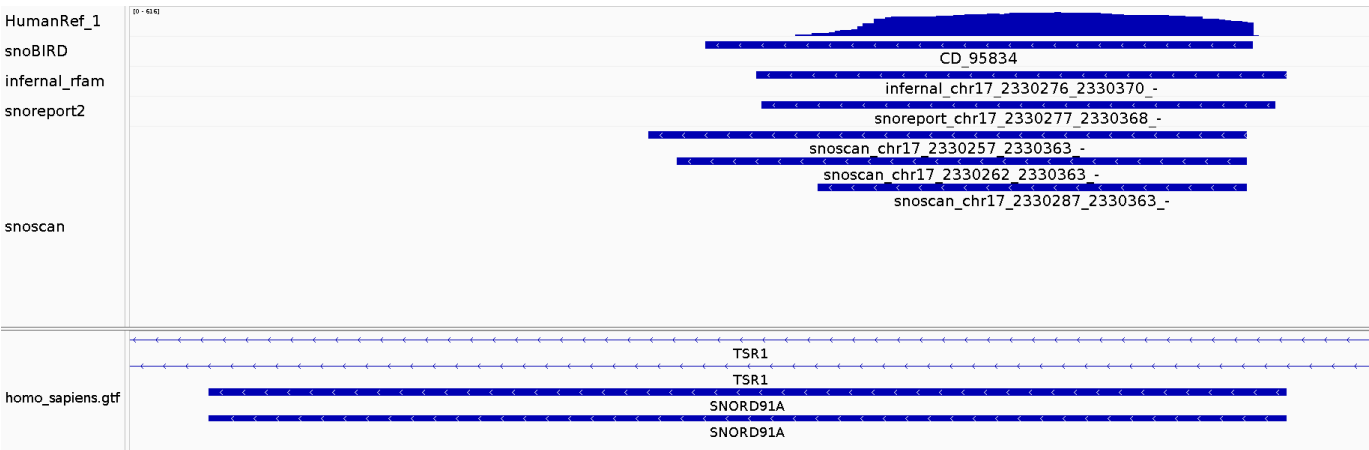

**Figure S19 (Supplemental to Figure 4). SnoBIRD can help reannotate existing C/D box snoRNAs.** Screenshots from the Integrative Genome Viewer (IGV) showing the current annotation of *SNORD91B* (**A**) and *SNORD91A* (**B**) in human and how SnoBIRD predictions can help reannotate accurate snoRNA limits according to read accumulation. For each panel, the following tracks are shown from top to bottom: read accumulation in the representative universal human reference sample 1 (HumanRef\_1) bedGraph, snoBIRD predictions, infernal\_rfam predictions, snoreport2 predictions, snoscan predictions and the current GTF annotation file showing both snoRNAs within their host gene *TSR1*.

A

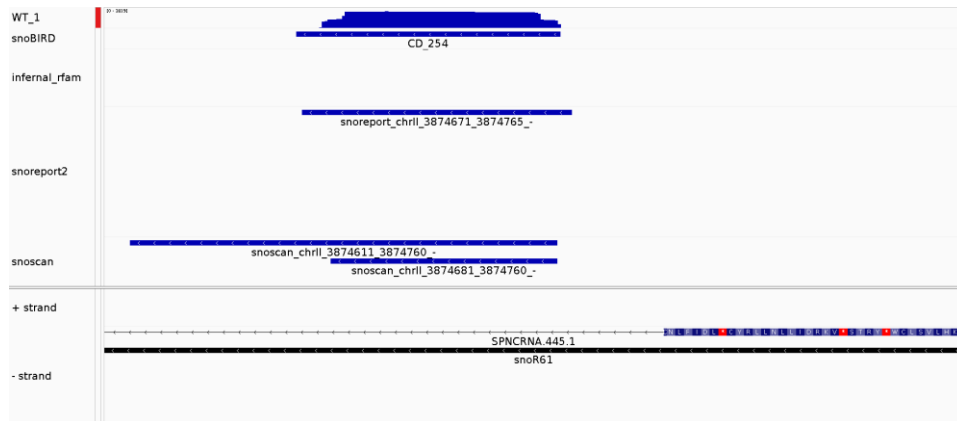

B

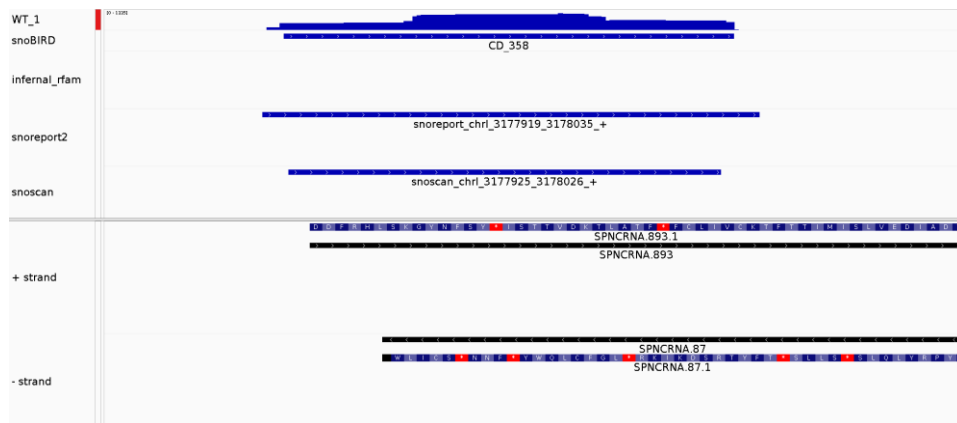

C

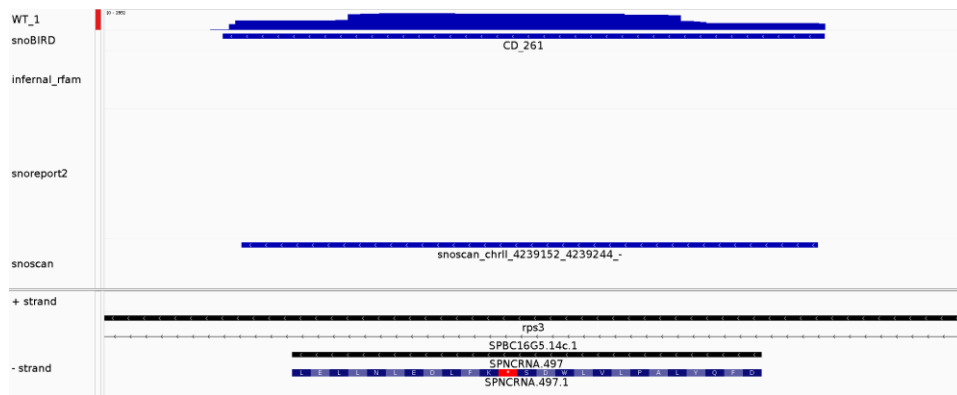

D

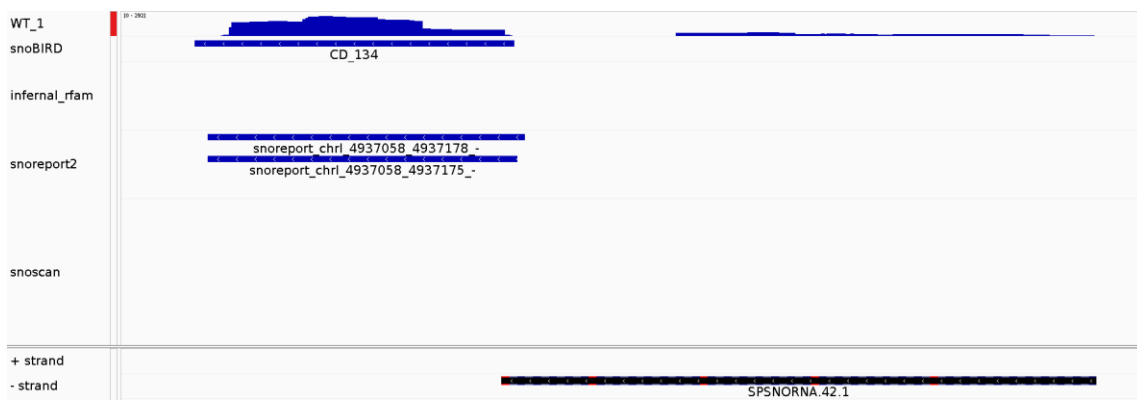

**Figure S20 (Supplemental to Figure 4). SnoBIRD's predictions in *S. pombe* help reannotate misannotated C/D box snoRNAs. (A)** Integrative Genome Viewer (IGV) screenshot showing CD\_254 within the long supposed snoR61 C/D box snoRNA. **(B)** IGV screenshot showing CD\_358 which overlaps with a supposed lncRNA. **(C)** IGV screenshot showing CD\_261 which overlaps with a supposed snRNA. **(D)** IGV screenshot showing CD\_134, a previously identified C/D box snoRNA missing from current annotations. All images include the tracks of read accumulation in a wild type (WT) strain, tools' predictions and current annotation.

### Overlap of predictions with genomic elements per tool and species

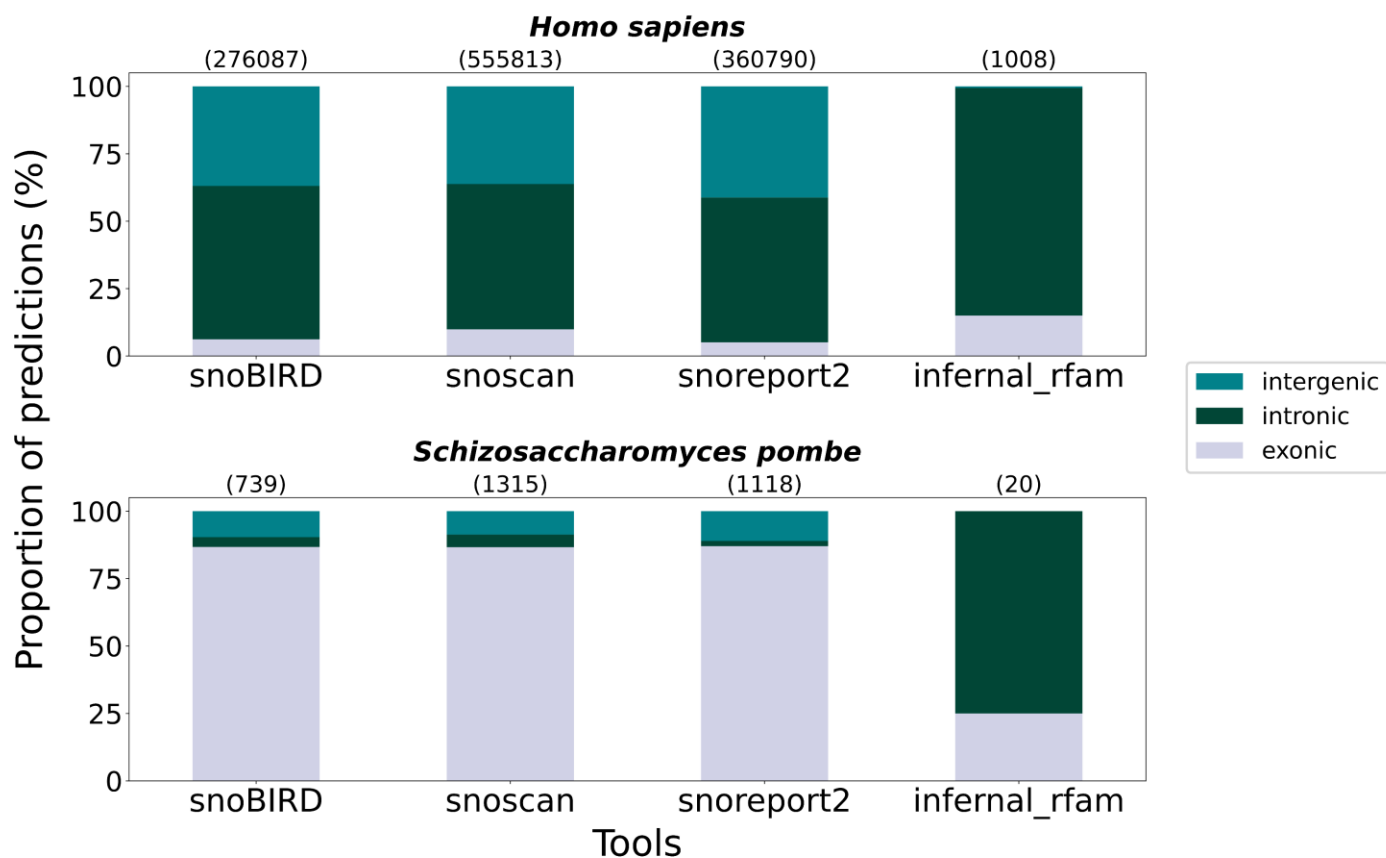

**Figure S21 (Supplemental to Figure 4). Overlap of predictions with genomic elements per tool for *H. sapiens* and *S. pombe*.** Stacked bar plots showing for *H. sapiens* (top panel) and *S. pombe* (bottom panel) the overlap of predictions with exonic, intronic and intergenic regions per tool. The total number of predictions per tool on a given genome is represented between parentheses above each bar.

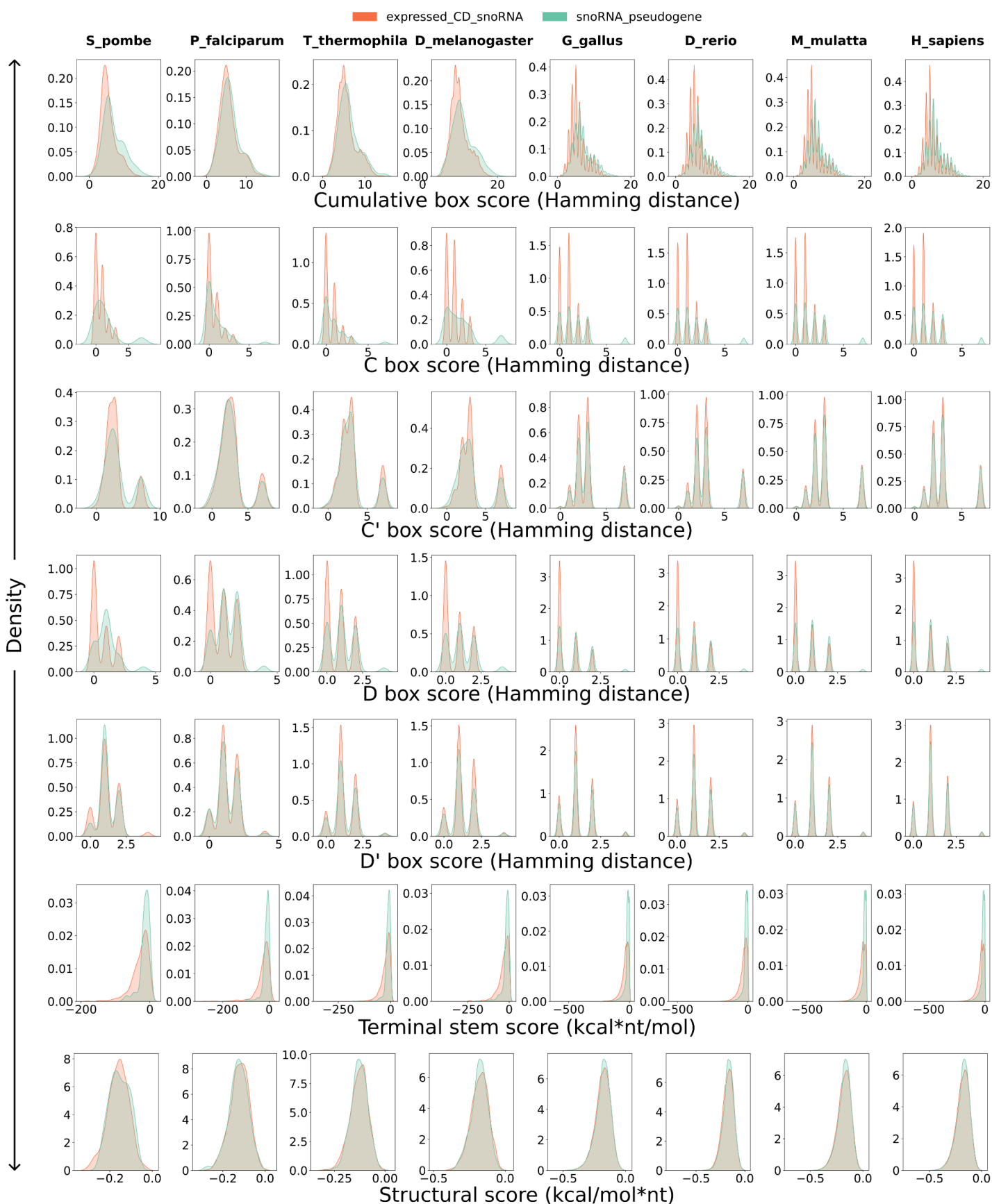

**Figure S22 (Supplemental to Figure 5). Feature distribution of the SnoBIRD predictions across different species.** Multi-density plots showing per row the different feature distributions between expressed C/D box snoRNA and snoRNA pseudogene predictions as a function of the species (i.e. per column). Hamming distance refers to the sum of mutations with regards to the respective consensus motif.

#### Overlap between expressed annotated C/D and expressed SnoBIRD predictions

*Drosophila melanogaster*

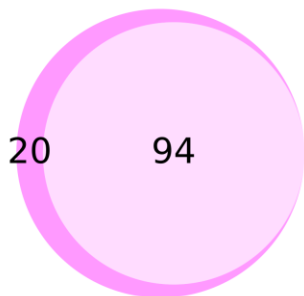

*Plasmodium falciparum*

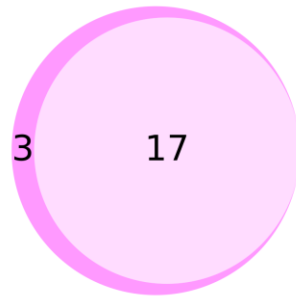

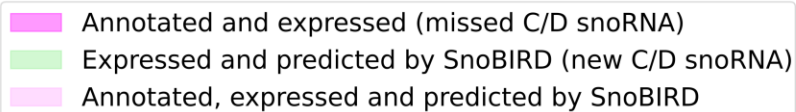

**Figure S23 (Supplemental to Figure 5).** SnoBIRD finds a large proportion of the annotated and expressed C/D box snoRNAs in *D. melanogaster* and *P. falciparum* but no new C/D box snoRNA candidates. Venn diagrams showing the intersection between annotated expressed C/D box snoRNAs and SnoBIRD's predictions across *D. melanogaster* and *P. falciparum* (the expression is based on TGIRT-Seq samples). No new C/D snoRNA was found by SnoBIRD in any of these species

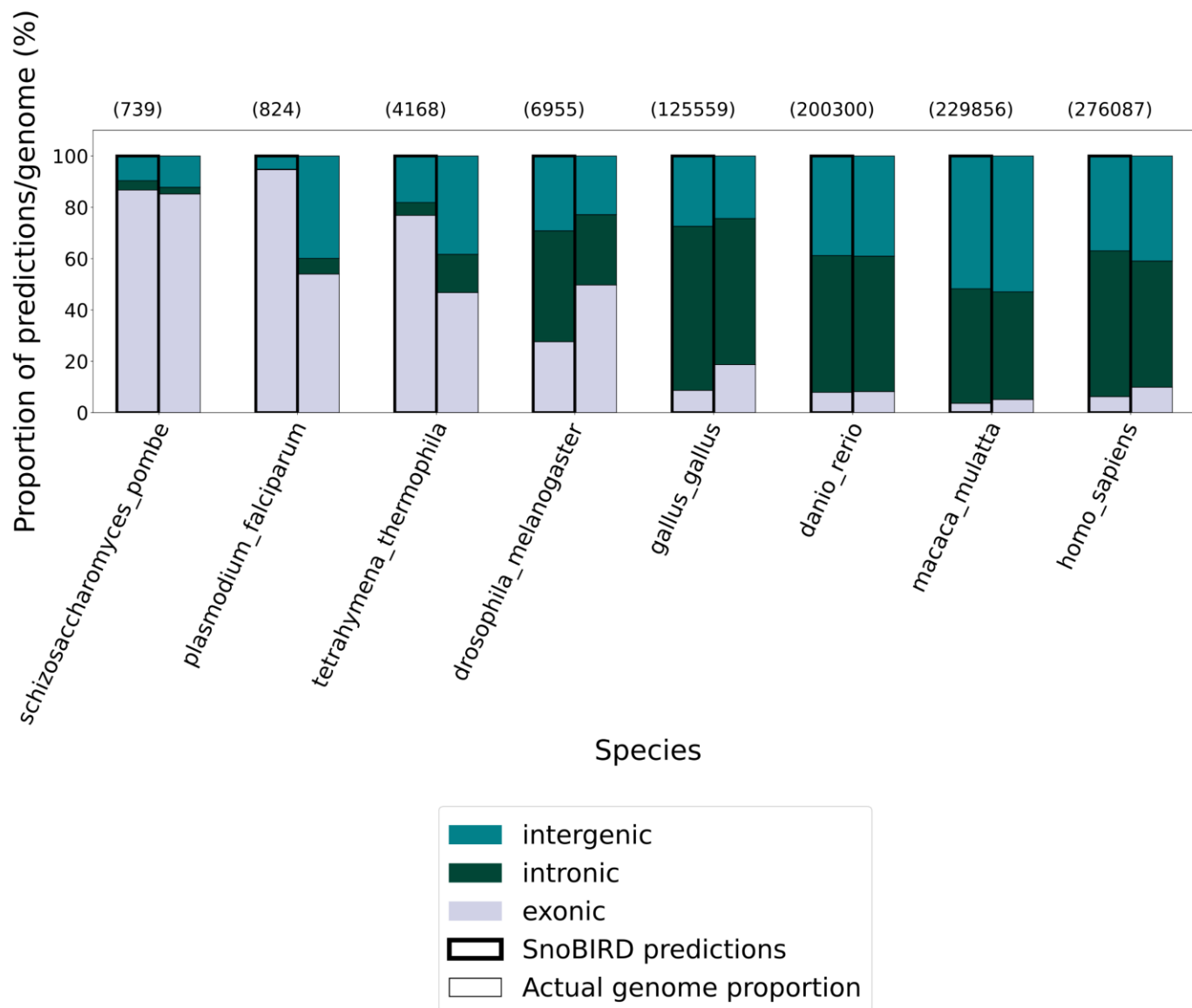

**Figure S24 (Supplemental to Figure 5). Overlap of SnoBIRD predictions with annotated genomic elements across different species genomes.** Stacked bar plots showing for different species the overlap of SnoBIRD predictions with exonic, intronic and intergenic regions (left bar with thick black borders). The actual genomic proportion of each element per species was obtained from Ensembl annotation files and is represented as the right bar with thin black borders. The total number of SnoBIRD predictions per species is represented between parentheses above each bar.

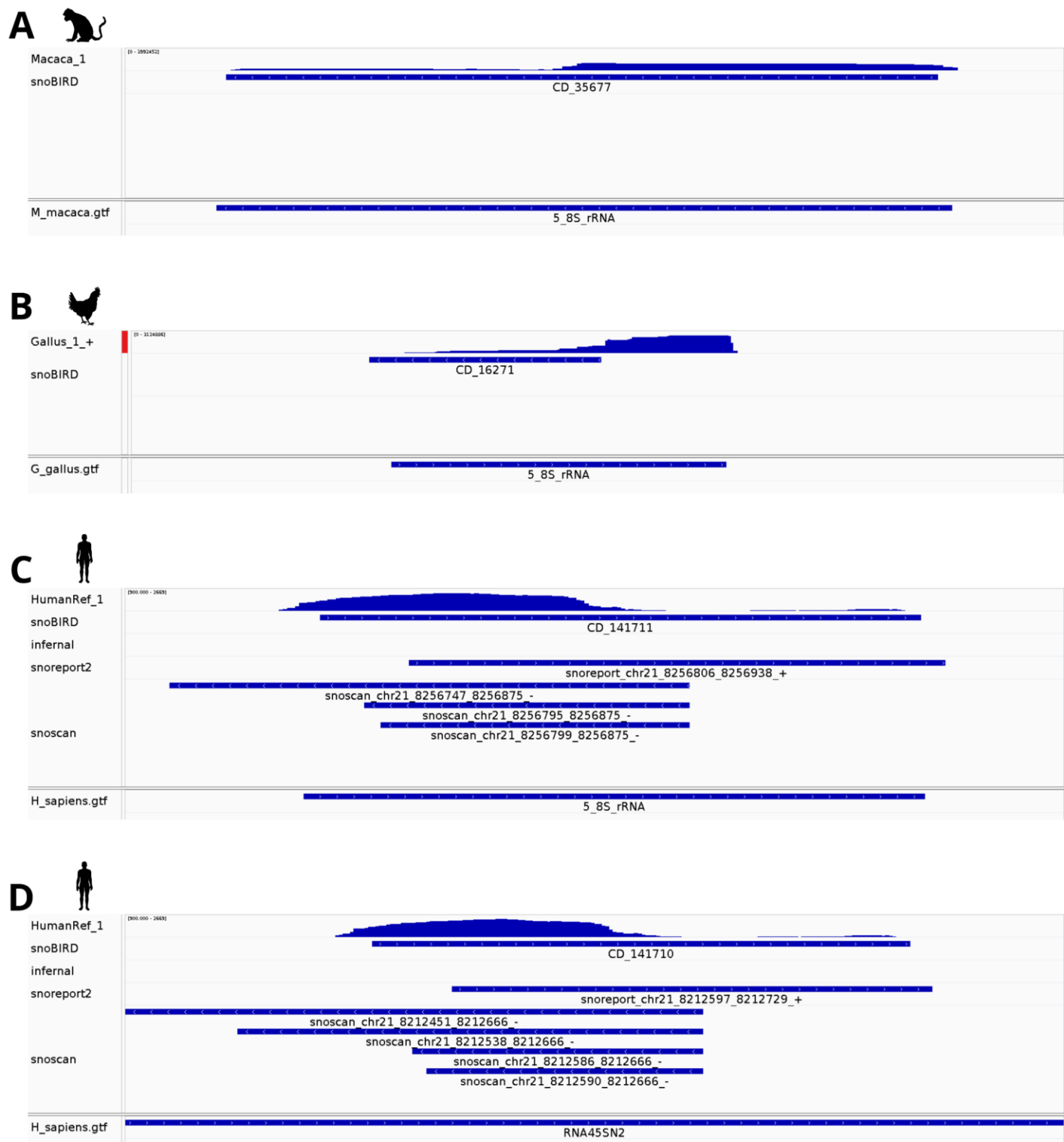

**Figure S25 (Supplemental to Figure 5). Several SnoBIRD predictions overlap with 5.8S ribosomal (r)RNA genes across different species.** Integrative Genome Viewer (IGV) screenshots of selected SnoBIRD predictions that overlap with annotated 5.8s rRNA genes (respectively in *Macaca mulatta* (**A**), *Gallus gallus* (**B**) and *Homo sapiens* (**C-D**), where in (**D**), the predicted snoRNA overlaps with the 5.8S portion of a 45S rRNA precursor gene). All images show as different tracks (from top to bottom) read accumulation in a representative sample per species, predictions by the different tools when applied in that species and current genomic annotation.

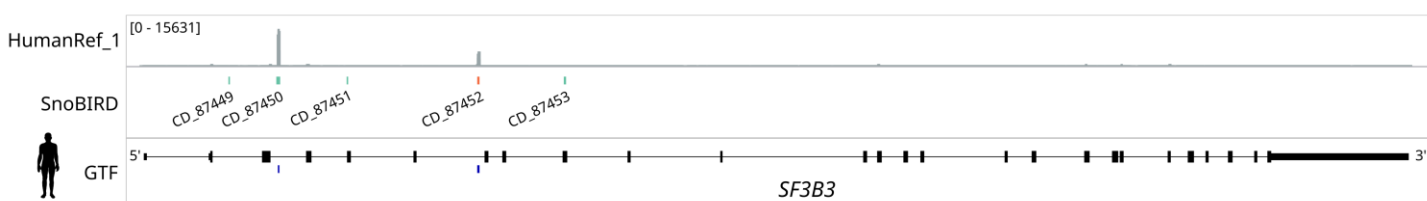

**Figure S26 (Supplemental to Figure 5). Genomic view of the conserved *SF3B3* gene locus in human.** The bottom track (GTF track) shows the exons and introns of *SF3B3* as black rectangles and lines respectively, as well as annotated C/D box snoRNA genes as blue rectangles below. The SnoBIRD track above the GTF track shows SnoBIRD predictions, with orange and teal rectangles representing expressed C/D snoRNAs and snoRNA pseudogenes, respectively. The top track shows read accumulation (bedGraph) along the locus for one representative sample, i.e. one universal human reference sample (HumanRef\_1).
